# The gerotherapeutic drugs rapamycin, acarbose, and phenylbutyrate extend lifespan and enhance healthy aging in house crickets

**DOI:** 10.1101/2025.08.25.671822

**Authors:** Gerald Yu Liao, Jenna Klug, Sherwin Dai, Swastik Singh, Elizabeth Bae, Warren Ladiges

## Abstract

The house cricket (*Acheta domesticus*) is a promising preclinical geroscience model due to its short lifespan, low maintenance, age-associated functional decline, and responsiveness to geroprotective drugs. Continuous dosing with rapamycin, acarbose, and phenylbutyrate extends lifespan; whether intermittent dosing offers similar benefits remains unknown. We tested 274 sex-matched crickets given 2-week intermittent dosing of each drug starting at mid-age (8-weeks), followed by behavioral testing at 10-weeks (geriatric stage). Assays included Y-maze olfactory discrimination, open-field exploration, and treadmill performance. Locomotor gaits were identified by velocity-based K-means clustering (silhouette > 0.5). A subset was monitored for post-treatment survival using Kaplan-Meier analysis. Olfactory preference was preserved by all drugs (*d*’s = −1.82 to −1.28, *P*’s < 0.01), with strongest effects in rapamycin-treated individuals. Rapamycin-treated males matched or exceeded juvenile locomotor activity; phenylbutyrate reduced male activity (*d* = 1.49, *P* < 0.05) and acarbose increased walking-to-running ratios (*d* = −0.75, *P* < 0.05). Rapamycin increased central exploration and freezing (*d* = −1.55, *P* < 0.0001), while acarbose and phenylbutyrate increased peripheral freezing (*d* = −0.76, *P* < 0.05). Rapamycin and phenylbutyrate extended maximum running time (*d*’s = −2.30 to −1.32, *P*’s < 0.0001), with sex-specific jumping gains in rapamycin-treated females and acarbose-treated males. Post-treatment lifespan was prolonged by rapamycin (*HR* = 0.42, *P* < 0.001) and reduced by acarbose in females (*HR*’s = 2.92 to 3.03, *P*’s < 0.05). Intermittent rapamycin preserved survival, cognition, and locomotion, while acarbose and phenylbutyrate produced selective benefits, supporting *A. domesticus* as a scalable model for geroprotective drug discovery.

## Introduction

The geroscience hypothesis posits that interventions targeting the biological mechanisms of aging can simultaneously delay or prevent multiple age-related diseases [1]. This framework has catalyzed the search for gerotherapeutics, agents designed to modulate the aging process itself rather than treat individual pathologies [2]. Among these, compounds such as rapamycin, acarbose, and phenylbutyrate have consistently extended lifespan and improved healthspan in mammalian models, demonstrating effects on conserved cellular pathways that regulate proteostasis, metabolism, and inflammation [3–5]. The translational challenge now lies in developing efficient, reproducible preclinical pipelines that bridge the gap between mechanistic discoveries and therapeutic application in humans [6].

Invertebrate models offer a powerful yet underexplored platform for this purpose. Unlike traditional mammalian models, they combine short lifespans, low maintenance costs, and high experimental throughput with evolutionary conservation of fundamental aging pathways [7–8]. The house cricket (*Acheta domesticus*) represents a particularly promising system. Previous work has demonstrated that crickets exhibit clearly defined morphological aging markers [9], measurable declines in locomotor (*manuscript in progress*) and cognitive performance [10–11], and histologically traceable age-associated changes (*manuscript in progress*). Furthermore, their modular nervous and organ systems allow for fine-grained behavioral and histological assessments [12], while their social and environmental responsiveness enables testing under conditions that closely mimic naturalistic aging dynamics [13]. Together, these features create an experimentally tractable platform that can interrogate both behavioral and cellular hallmarks of aging with high resolution.

Despite the evolutionary distance between insects and humans, core mechanisms of aging (e.g., metabolic dysregulation, neurodegeneration, immune senescence, and intestinal barrier failure) are remarkably conserved across species [8, 14]. This conservation raises the possibility that pharmacological interventions effective in mammals may also attenuate age-related decline in crickets, providing an intermediate preclinical step prior to vertebrate and human studies (Fig. 1). By validating whether FDA-approved geroprotective agents act on conserved pathways in crickets, we can leverage this model to accelerate drug screening and identify mechanisms with strong translational potential.

**Figure 1.**
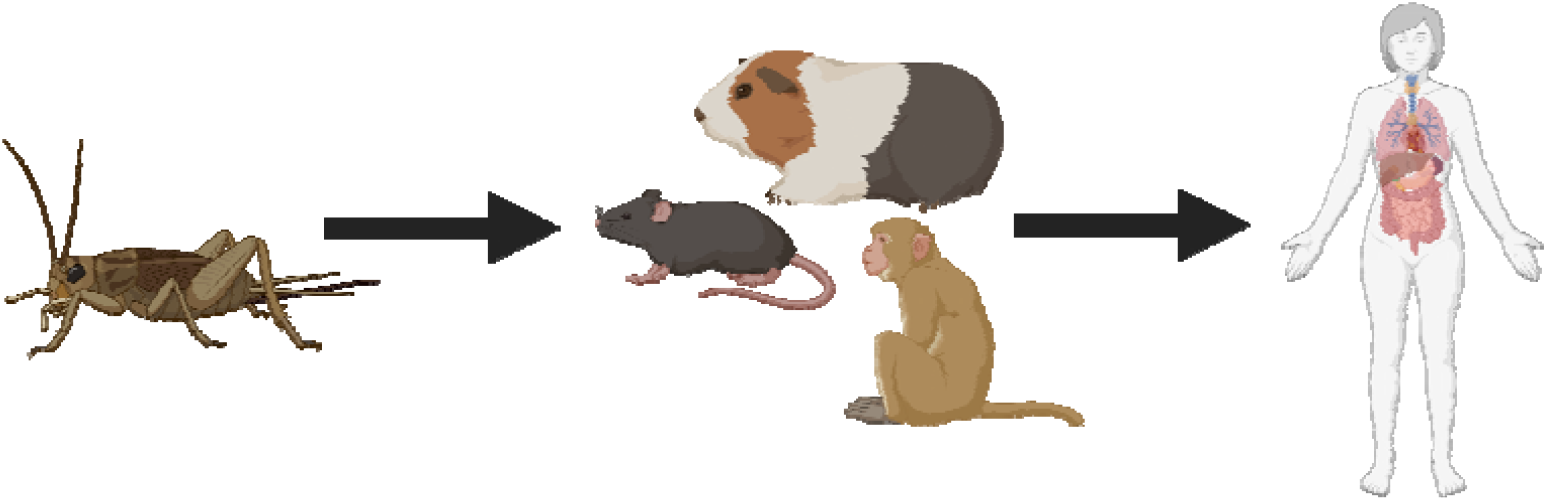
Translational potential of the domestic house cricket as a preclinical platform for gerotherapeutic discovery. Conceptual framework illustrating the use of house crickets as a model for screening and mechanistic evaluation of candidate gerotherapeutics. Illustration created using Biorender.com.

Here, we investigate the effects of three candidate gerotherapeutics (rapamycin, acarbose, and phenylbutyrate) administered individually to aged crickets. These agents target complementary aspects of aging biology: rapamycin inhibits mechanistic target of rapamycin complex 1 (mTORC1) to promote autophagy and suppress senescence [15–16]; acarbose, an α-glucosidase inhibitor, modulates glucose metabolism to mimic caloric restriction and reduce oxidative stress [17]; and phenylbutyrate, a histone deacetylase (HDAC) inhibitor that enhances proteostasis and epigenetic stability while mitigating neuroinflammation [18]. We hypothesize that intermittent treatment with these compounds will extend lifespan, preserve neuromuscular function, and improve cognitive performance in geriatric crickets. By systematically characterizing age-associated functional deterioration and assessing the efficacy of geroprotective interventions, this study aims to establish the house cricket as a scalable and translational model for aging research, bridging mechanistic insights across species.

## Materials and Methods

### Cricket rearing, housing, and experimental design

House crickets were sourced from a commercial supplier (Fluker Farms Inc, Louisiana, USA) and maintained under standardized husbandry in dual-layer Plexiglass enclosures permitting containment and self-regulated light exposure [9]. Environmental conditions were held at 29 ± 1°C and 32 ± 3% relative humidity [7], with shaded refuges for voluntary light avoidance and thermoregulation. Colonies were maintained at 20–30 individuals per cage (10–15 cm^2^ per cricket) to optimize social interaction without overcrowding [9, 13]. Both sexes were co-housed outside specific-pathogen-free conditions to better model wild-type aging trajectories [19].

Crickets arrived at 5 weeks of age, acclimated for 3 weeks on a control cricket diet [9] and water gel packs (Napa Nectar Plus), and were randomly assigned at 8 weeks to control or treatment groups. Treatments were administered for 2 weeks, targeting the mid-life period when moderate functional decline emerges, and behavioral testing was performed at 10 weeks, an age associated with pronounced geriatric impairment [9]. Survival monitoring was conducted in a subset maintained on assigned diets until natural death. Performance was compared with archived younger cohorts to assess rejuvenation effects and with historical geriatric and pooled controls to exclude batch-related artifacts.

### Diet preparation and drug administration

Phenylbutyrate, rapamycin, and acarbose have been shown to extend lifespan in the house cricket [20], and controlled variation in drug-infused food intake exerts negligible effects on survival in this species [21]. All crickets were maintained on a standardized diet composed of Picolab Rodent Diet 20 (5053, irradiated; Purina Mills, USA) bound with gelatin. Gelatin was dissolved in water at 100 °C, cooled to 60 °C, blended with powdered chow, refrigerated overnight at 20 °C, dehydrated to prevent mold growth, and homogenized to a uniform particle size.

Drug-supplemented diets were formulated at dosages previously effective in murine studies [22], with preparation adjusted for compound-specific thermal stability and solubility. Rapamycin (14 ppm; Southwest Research Institute, San Antonio, TX), stable up to 60 °C [23] and insoluble in water [24], was incorporated directly into the gelatin–chow mix. Phenylbutyrate (1000 ppm; Triple Crown America Inc., Perkasie, PA), stable up to 30°C [25] and water-soluble [26], was pre-dissolved in deionized water before incorporation to ensure even distribution. Acarbose (1000 ppm; Spectrum Chemical Mfg Corp, Gardena, CA), stable up to 40 °C [27] and water-soluble [28], was prepared in the same manner as phenylbutyrate. Control groups received the identical base diet without supplementation.

### Olfactory-guided decision-making

We adapted a Y-maze paradigm from Liao et al., 2025 to assess odor discrimination and decision-making. Briefly, one choice arm was infused with an attractive vanilla stimulus (1% extract, 1:1 dilution), and the other with a repellent cinnamon odorant (1% extract, 1:1 dilution) [10]. Individual crickets were placed in the start arm for 10s, then allowed up to 30s to choose an arm; entry was recorded when the entire body crossed the threshold. Odor positions alternated between trials to prevent spatial bias. If a cricket failed to enter a choice arm within the allotted time, the trial was excluded from analysis. Each cricket underwent a total of eight consecutive trials, with an inter-trial interval of 30s to minimize residual olfactory cues from prior choices. To quantify cognitive decision-making via odor preference, percentage of vanilla-scented arm entries served as the primary outcome measure.

### Open field test

Locomotor and exploratory behaviors were assessed in a 5-min open field assay conducted in darkness (*manuscript in progress*). Briefly, crickets were placed individually at the arena center and behavior was video recorded for automated analysis in ezTrack [29]. Central and peripheral zones were manually defined, and positional data were extracted at 30 frames per second (FPS) to quantify distance traveled, average speed, freezing episodes, and zone occupancy. Velocities exceeding the 99th percentile were excluded to remove tracking artifacts. Walking and running thresholds were determined for each treatment group via k-means clustering (*k* = 2) on velocity data, enabling classification of distance, duration, and speed for each gait. All measures were computed separately for central and peripheral zones.

### Induced locomotion assay

Locomotor performance was assessed using a modified rodent treadmill (Exer 3/6, Columbus Instruments) adapted for crickets by disabling the shock grid, adding dividers to prevent lateral escape, and installing paper catch systems for safe recovery (manuscript in preparation). A vanilla-scented filter paper at the terminal end served as a motivational cue [30]. Trials were conducted at 0° incline with an acceleration of 0.1 m/s2. Horizontal jump distances were measured via sidewall rulers, and each individual completed three trials; the best performance was used for analysis. Primary endpoints were running duration (s) and maximum jump distance (cm).

### Euthanasia and morphometrics

Crickets were euthanized using controlled CO_2_ exposure followed by decapitation at the cranio-cervical junction, in accordance with AVMA guidelines [31]. Morphometric measurements were collected immediately post-mortem under standardized conditions. Body length was measured from the frons to the abdominal tip, excluding appendages such as cerci, wings, and ovipositor. Antennal length was determined as the average of both antennae, while hindleg length was recorded with the limb fully extended along its natural axis. Body mass was recorded before and after experimental procedures using an analytical balance (Mettler Toledo XPR205, Switzerland).

Individuals with damaged or missing antennae at euthanasia were omitted from antennal analyses. Crickets that died before the planned endpoint were excluded from all morphological assessments to avoid potential artifacts from post-mortem degradation or cannibalism.

### Statistical Analysis

Group-level characteristics were summarized using standard descriptive statistics. Categorical variables were reported as counts and percentages and compared by χ^2^ tests or Fisher’s exact tests when ≤ 80% of cells contained n ≤ 5. Effect sizes were expressed as relative risk (RR) with 95% confidence intervals (CIs), with a modified Haldane-Anscombe correction applied for zero counts [32]. Continuous variables were expressed as means ± standard deviations (SD) and compared using one-way analysis of variance (ANOVA); effect sizes were calculated as Cohen’s *d* with Hedges’ *g*, with 95% CIs computed using a standardized effect size calculator [33–35].

Analyses were stratified by treatment group and sex. Normality was assessed by the Shapiro-Wilk test, with large samples (N > 30) assumed to follow a normal distribution. Normally distributed data were analyzed with parametric tests; otherwise, non-parametric methods were used. One-way ANOVA or Kruskal-Wallis test evaluated group-level differences, with Dunnett’s post-hoc test applied when comparing to a control or reference group. Two-way ANOVA assessed main and interactive effects of treatment and sex; when interaction terms were not significant, main effects were followed by Bonferroni-adjusted within-sex comparisons and Dunnett’s between-group tests.

Survival analyses used Kaplan-Meier estimates, with group differences tested by log-rank (Mantel-Cox) analysis. Hazard ratios (HRs) and *P*-values were calculated, and multiple comparisons were corrected using the Benjamini-Hochberg method.

Analyses were conducted in GraphPad Prism v10.0.3 (GraphPad Software) or Python v3.13.1 (Python Software Foundation), using SciPy v1.13.1 for inferential statistics and scikit-learn [36] for machine learning. Figures were generated with matplotlib v3.9.2 or GraphPad Prism. Statistical significance was set at α = 0.05.

## Results

### Cohort characteristics and morphometric analyses

Experimental cohorts were sex-balanced and tested between February 27^th^ and April 23^rd^, 2025. Sex distributions did not differ across groups in the scent preference (N = 79), open field (N = 99), or treadmill (N = 96) assays (P’s > 0.05; Appendix 1).

Morphometric analyses of validated body size and proportional traits [9] revealed no treatment effects (*P*’s > 0.05), and these variables were therefore excluded as covariates in behavioral models. Consistent with prior reports [9], within-group sex comparisons showed females and males differed in body weight, body length, and multiple scaling ratios (*P*’s < 0.05). These data are provided in Appendix 2 for completeness but were not incorporated into final behavioral models to maintain focus on treatment effects.

### Olfactory distinction declines with age and is preserved by anti-aging interventions

In the Y-maze scent preference task, all medicated groups exhibited increased vanilla arm entries (*d*’s = −1.82 to −1.28; *P*’s < 0.01) (Fig. 2A). This effect was most pronounced in males, with rapamycin-treated females also outperforming controls (*d*’s = −1.91 to −1.38; *P*’s < 0.05) (Fig. 2B). Relative to adults, controls displayed reduced olfactory preference, whereas treated groups performed at adult-like levels. Decline was observed primarily in males, with no differences detected in females. Compared to mid-aged and historical geriatric cohorts, rapamycin consistently improved olfactory performance, with acarbose and phenylbutyrate showing weaker or variable effects. Similar trends were observed when compared to pooled historical controls.

**Figure 2.**
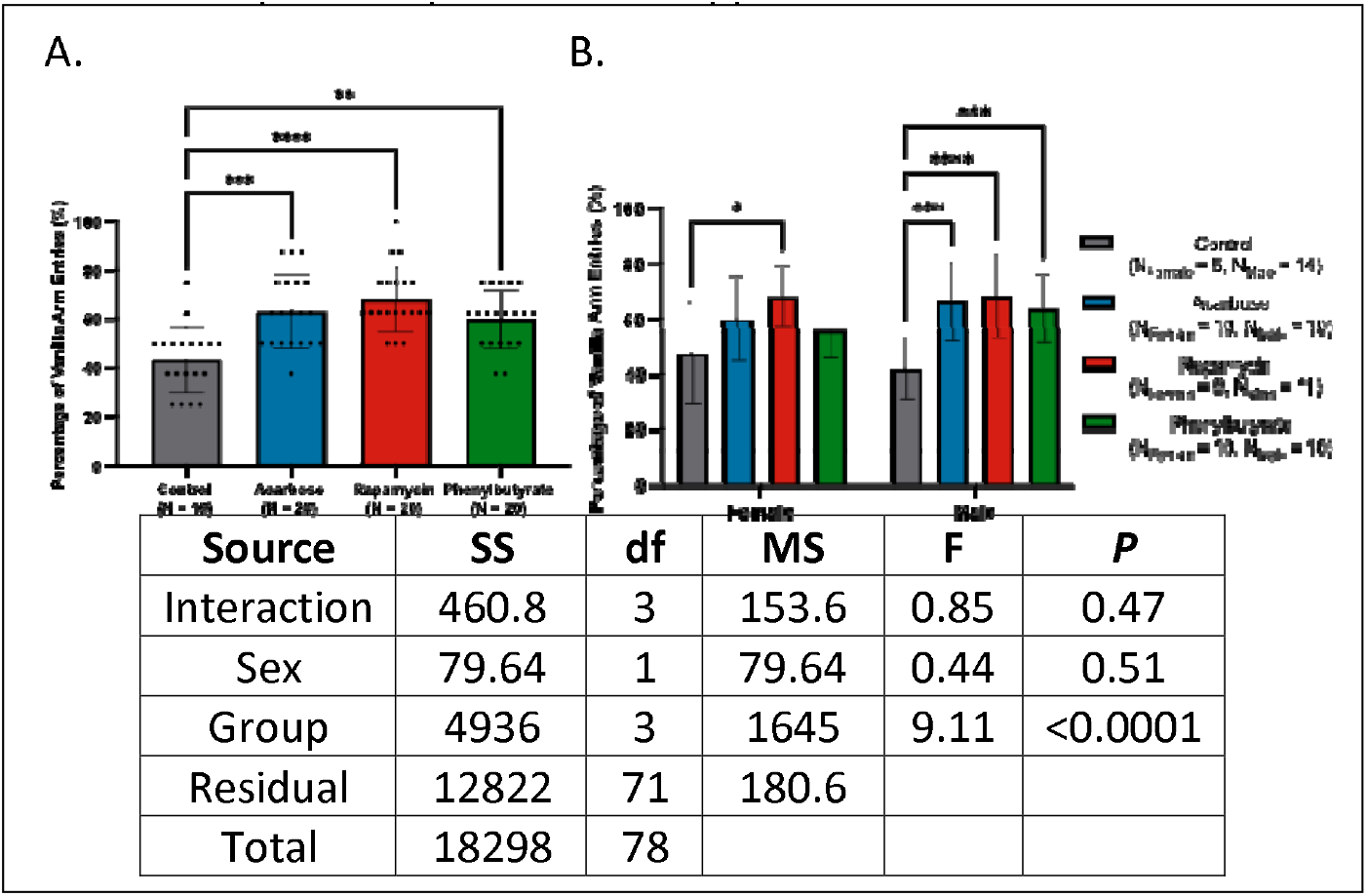
Anti-aging interventions preserve olfactory recognition memory in aged crickets. **(A)** Acarbose, rapamycin, and phenylbutyrate treatments improved vanilla preference relative to controls in geriatric crickets. **(B)** Rapamycin-treated females and all treated males showed enhanced vanilla preference (\**P* < 0.05, \*\**P* < 0.01, \*\*\**P* < 0.001, \*\*\*\**P* < 0.0001).

Full statistical outputs are presented in Appendix 3.

### Velocity-defined clusters revealed well-defined walking and running categorization

Walking and running cutoffs were derived via unsupervised K-means clustering (*k* = 2) of velocity data from ezTrack’s center-of-mass displacement “Location_output” [29], excluding values above the 99^th^ percentile. Clustering consistently yielded two distinct velocity distributions corresponding to walking and running, with rapamycin-treated crickets exhibiting the highest mean velocities across both gaits (Figure 3A-D).

**Figure 3.**
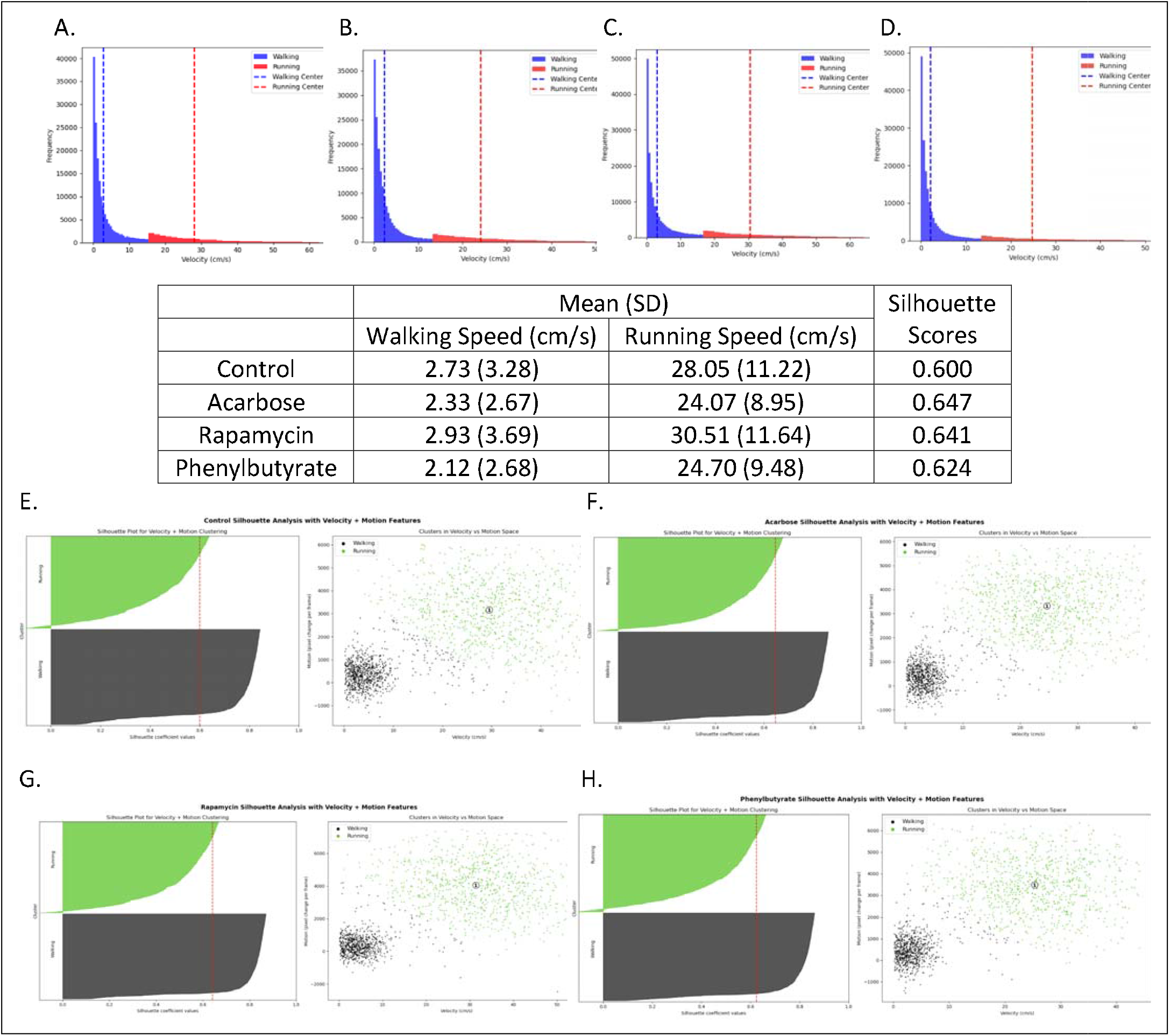
Treatment-dependent velocity distributions and silhouette-based cluster validation. **(A-D)** Velocity distributions for control, acarbose, rapamycin, and phenylbutyrate cohorts (left to right) cohorts reveal treatment-dependent differences in maximal speed profiles. **(E-H)** Two-dimensional K-means clustering using velocity and motion magnitude metrics for **(E)** control, **(F)** acarbose, **(G)** rapamycin, and **(H)** phenylbutyrate. Left: Silhouette plots illustrating the compactness and separation of clusters; Right: Cluster visualizations showing distinct separation between walking and running datapoints. In all conditions, average silhouette scores (vertical red lines) exceeded 0.5, indicating well-defined cluster boundaries based on both center-of-mass velocity and independent motion magnitude metrics.

Cluster validity was confirmed using an orthogonal metric, motion magnitude from frame-to-frame pixel changes in ezTrack’s “Freezing_output.” Two-dimensional clustering of velocity and motion magnitude, followed by silhouette analysis [37], produced scores >0.5 in all groups, confirming well-separated clusters [38]. Representative silhouette plots and cluster visualizations are presented in Figs. 3E-H.

### Rapamycin preserves male-specific restoration of juvenile-like locomotor performance

Total distance traveled was comparable between controls, acarbose-, and rapamycin-treated crickets, with phenylbutyrate-treated individuals trending lower (*d* = 0.85 [95% CI: 0.20, 1.51], *P* > 0.05; Fig. 4A). Sex-stratified analyses revealed reduced activity in phenylbutyrate-treated males (*d* = 1.49 [95% CI: 0.50, 2.48], *P* = 0.049; Fig. 4B). All groups performed below juveniles, except rapamycin-treated males, who matched juvenile levels and exceeded adults. Against historical geriatrics, activity was higher in current control and rapamycin groups, driven by males and rapamycin-treated females; only rapamycin-treated crickets exceeded pooled controls.

**Figure 4.**
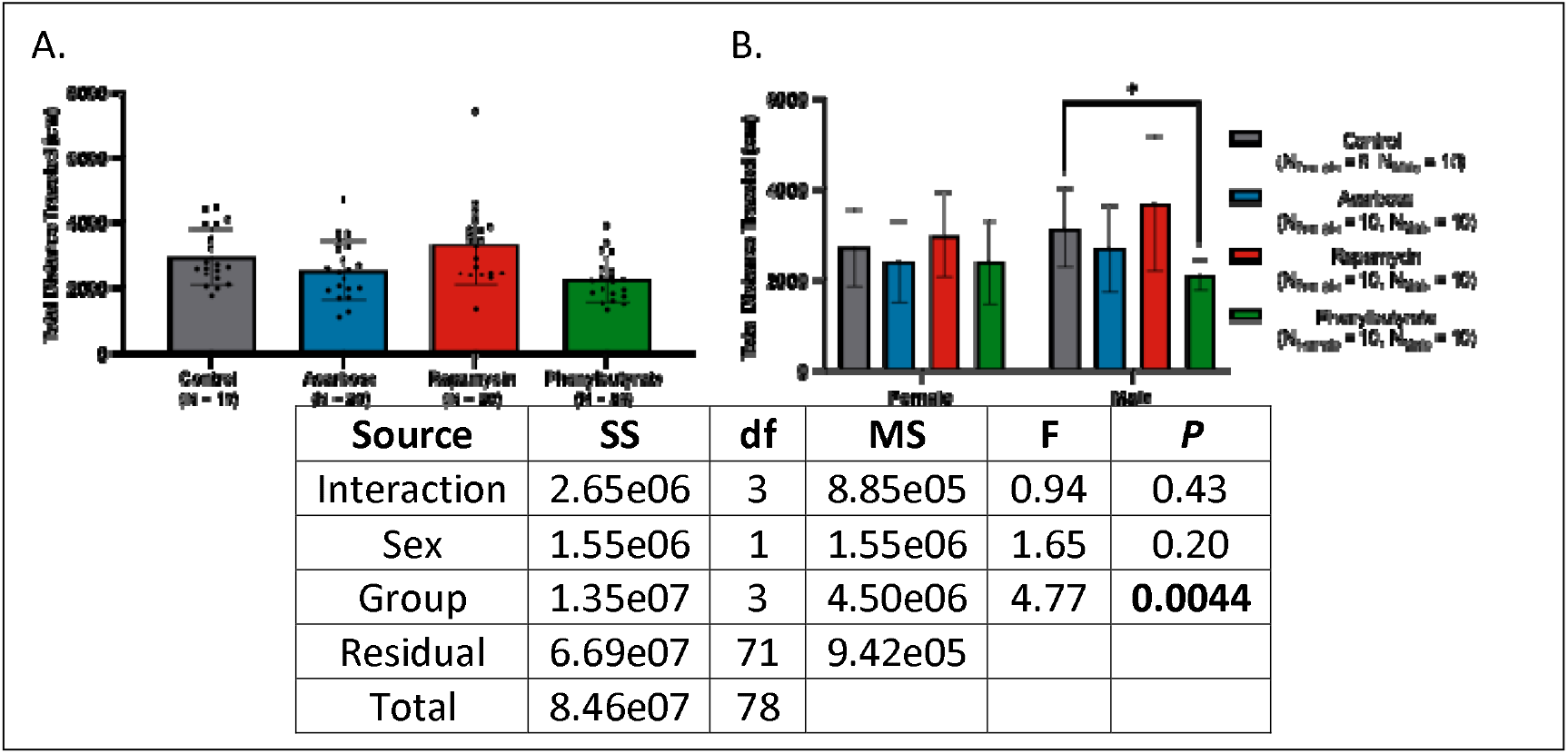
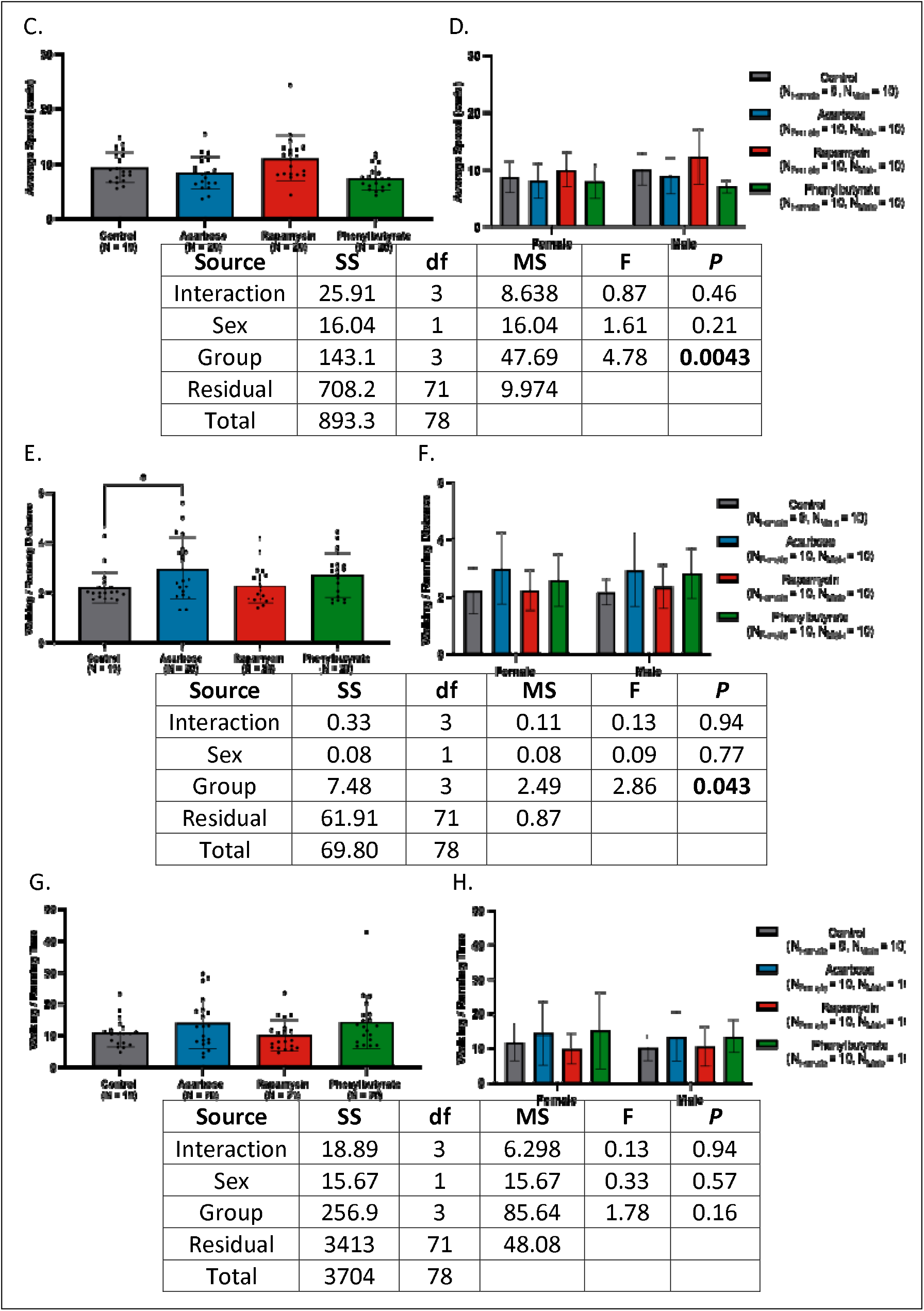
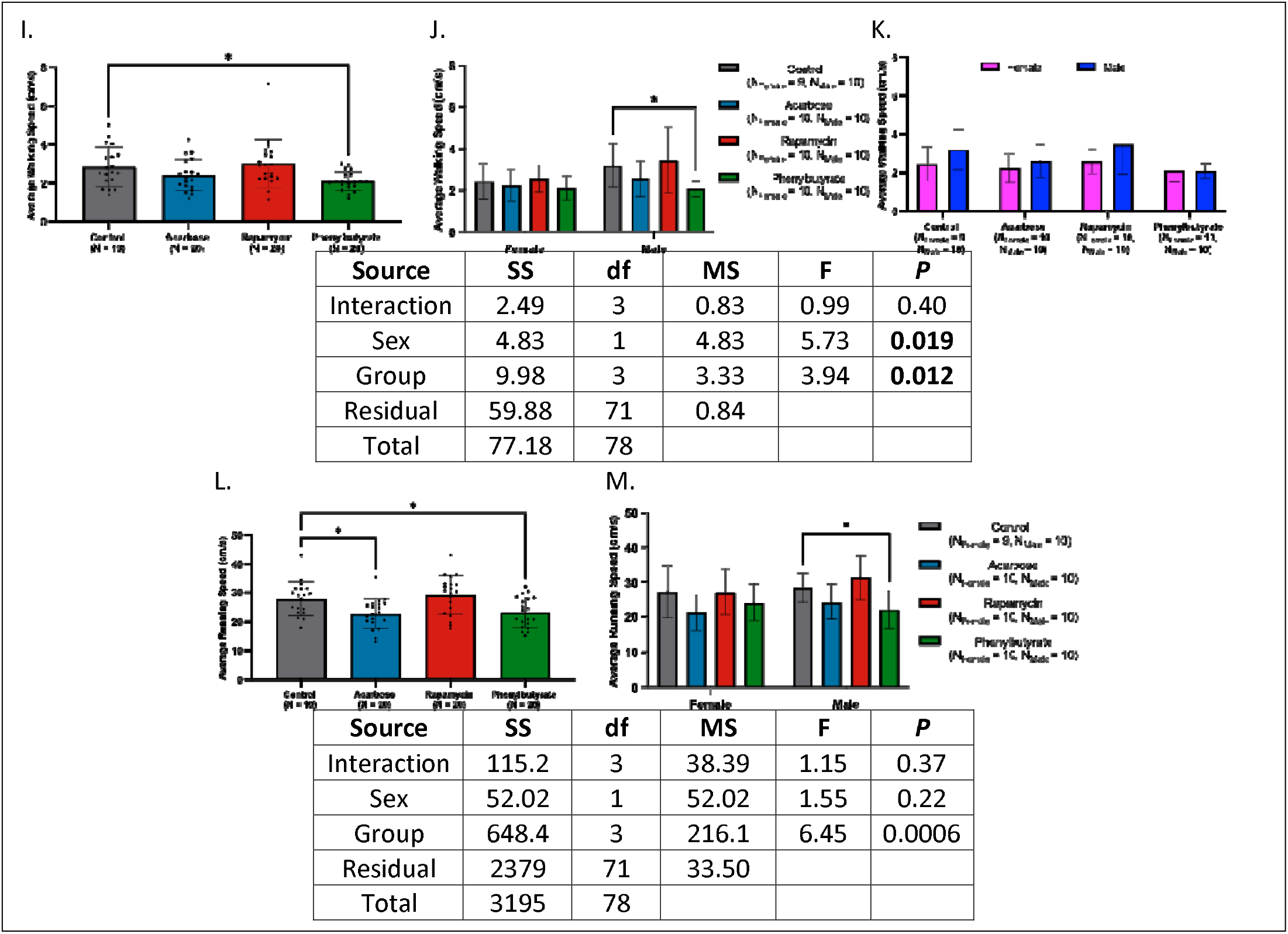
Effects of pharmacological interventions on locomotor performance in geriatric crickets. **(A)** Total distance traveled was comparable across treatment groups. **(B)** Phenylbutyrate-treated males covered less distance than controls. **(C)** Average speed did not differ in the overall cohort, although phenylbutyrate trended lower. **(D)** Sex-stratified analyses revealed no treatment effects, aside from a trend toward reduced speed in phenylbutyrate treated males. **(E)** Acarbose-treated crickets exhibited elevated walking-to-running distance ratios relative to controls. **(F)** This effect was absent when stratified by sex. **(G)** Walking-to-running time ratios were similar across all treatments. **(H)** No sex-specific differences were detected. **(I)** Walking speed was reduced only in phenylbutyrate-treated individuals. **(J)** This reduction persisted in males only. **(K)** No sex differences were observed within treatment groups. **(L)** Running speed was reduced in acarbose- and phenylbutyrate-treated crickets compared to control. **(M)** Sex-stratified analysis identified reduced running speed only in phenylbutyrate-treated females (\**P* < 0.05, \*\**P* < 0.01, \*\*\**P* < 0.001, \*\*\*\**P* < 0.0001).

Average speed was similar across control, acarbose-, and rapamycin-treated groups, but trended lower with phenylbutyrate, particularly in males (*d*’s = 0.75 to 1.32, *P*’s > 0.05); Figs. 4C-D). Only rapamycin-treated males retained juvenile-like speeds and exceeded adults. Versus historical geriatrics, control and rapamycin groups were faster; only rapamycin-treated crickets surpassed pooled controls, driven by males.

Walking-to-running distance ratios were elevated in acarbose-treated crickets (*d* = −0.75 [−1.40, −0.10], *P* < 0.05; Fig. 4E-F). Only rapamycin and control groups were indistinguishable from juveniles and reduced compared to historical geriatrics, particularly in males. Against pooled controls, only rapamycin trended higher without sex-specific effects.

Walking-to running time ratios showed no differences among treatments (Figs. 4G-H). Rapamycin-treated females and control males retained juvenile-like values, while only rapamycin-treated crickets trended lower than pooled controls.

Average walking speed was reduced only in phenylbutyrate-treated crickets, driven by males (*d*’s = 0.90 to 1.32, *P*’s < 0.05) (Figs. 4I-K). All groups walked slower than juveniles; versus adults and historical geriatrics, only control and rapamycin groups were faster, driven by males; versus pooled controls, only rapamycin-treated males were faster.

Running speed was reduced in acarbose-treated crickets and phenylbutyrate-treated males (*d*’s = 0.92 to 1.25, *P*’s < 0.05) (Figs. 4L-M). Only rapamycin-treated males matched juveniles and surpassed pooled controls, while both rapamycin-treated and control males exceeded adult and historical geriatric levels.

Full statistics are provided in Appendix 4.

### Rapamycin offsets spatial bias and alters freezing dynamics

In central-to-peripheral distance ratios, only acarbose-treated females and phenylbutyrate-treated males trended lower than controls (Figs. 5A-B). Versus juveniles, central engagement was reduced in acarbose-treated females and phenylbutyrate-treated males while only acarbose-treated females were lower than adults. Compared to historical geriatric and pooled controls, rapamycin-treated individuals and phenylbutyrate-treated females had elevated ratios.

**Figure 5.**
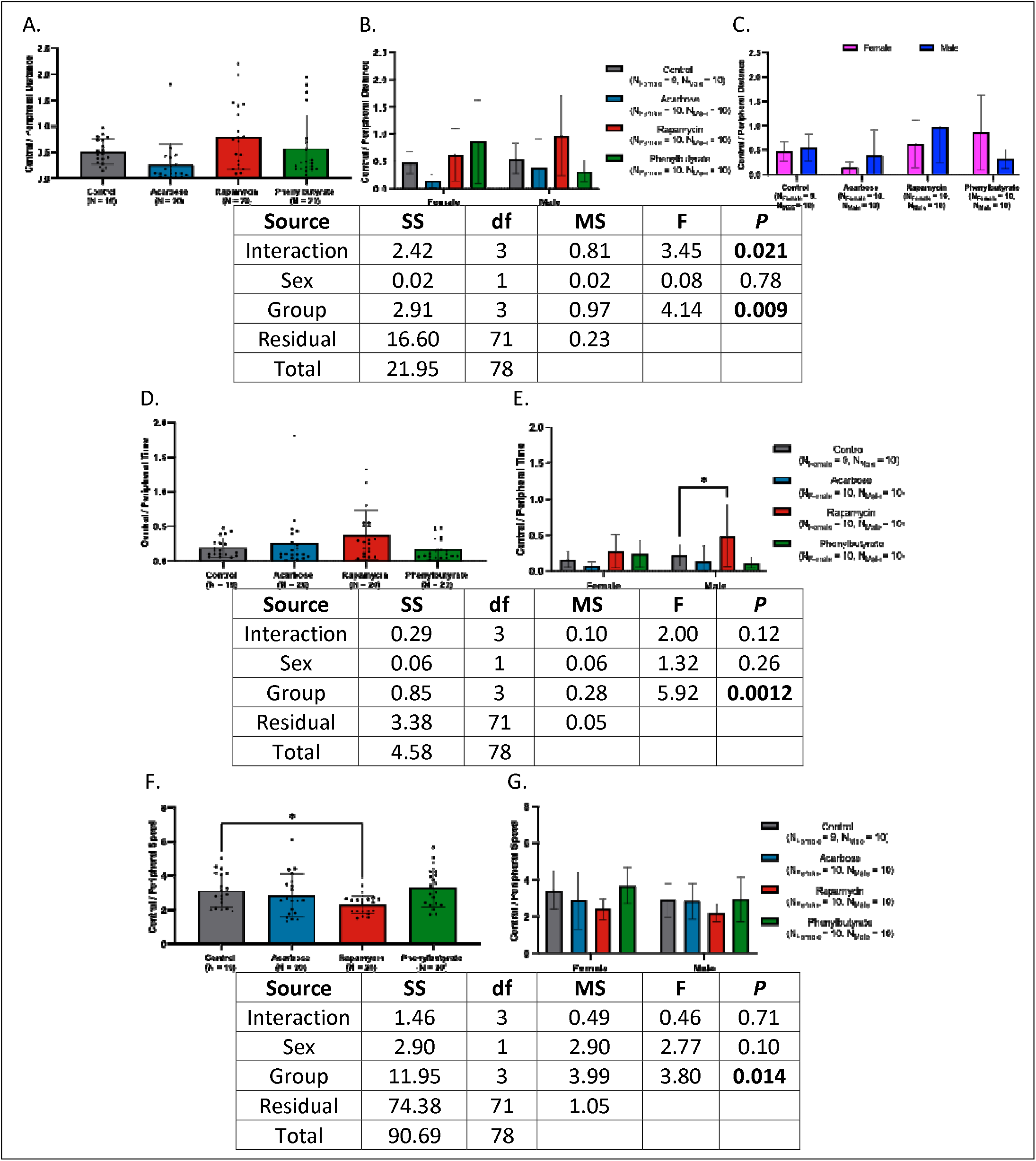
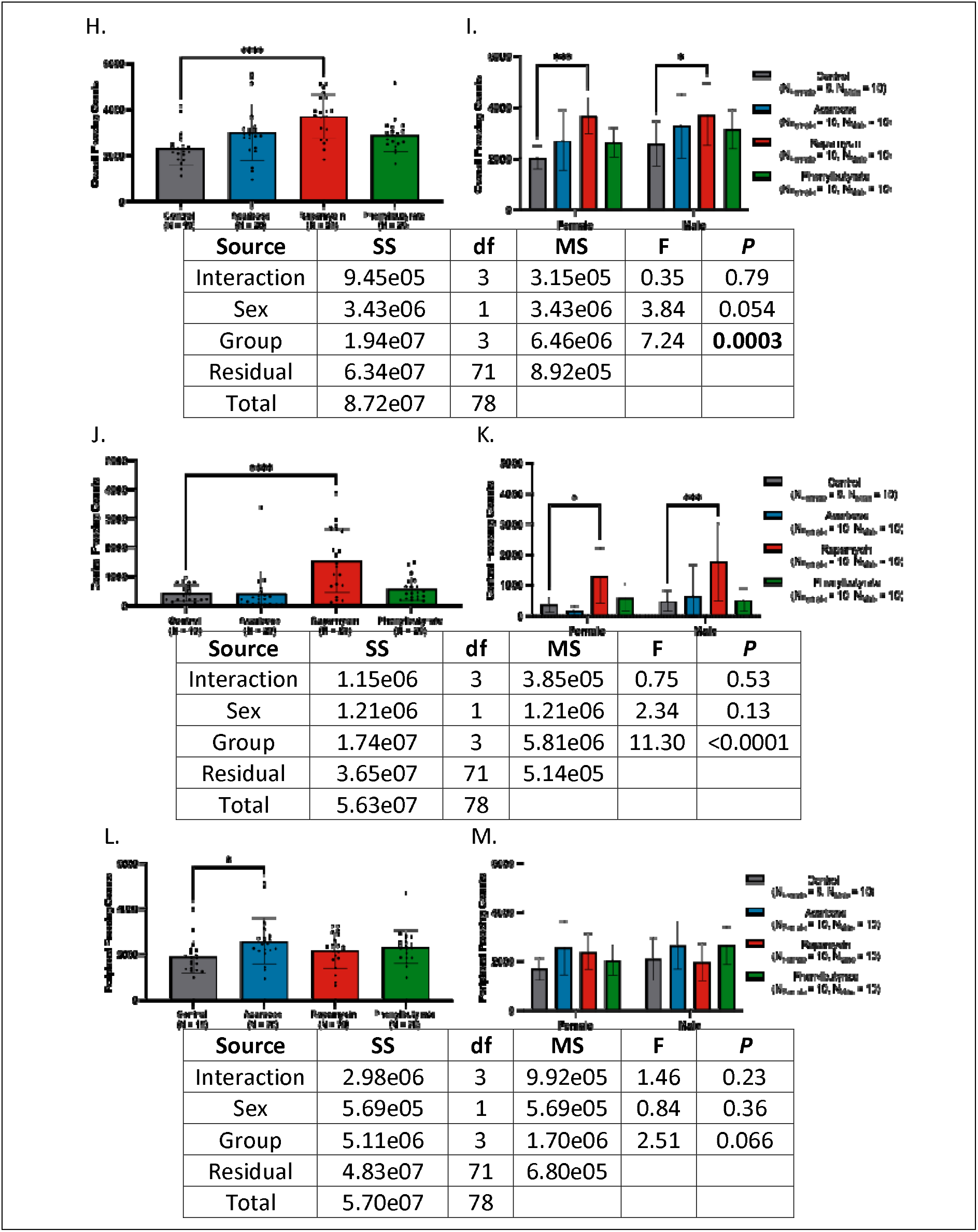
Effects of pharmacological treatments on exploratory behavior and freezing dynamics. **(A)** Central-to-peripheral distance ratios were comparable across treatments, with acarbose-treated crickets trending lower than controls. **(B)** Sex-stratified analysis revealed reduced exploration trends in phenylbutyrate-treated males and acarbose-treated females. **(C)** Phenylbutyrate-treated females exhibited higher ratios than males, a pattern absent in other treatments. **(D)** Rapamycin-treated crickets trended toward greater central zone occupancy relative to controls. **(E)** This effect was restricted to males. **(F)** Central zone speed was lower in rapamycin-treated crickets than in controls. **(G)** No sex-specific differences in central speed were detected. **(H)** Rapamycin-treated crickets froze more than controls. **(I)** Increased freezing with rapamycin was observed in both sexes. **(J)** Central freezing was elevated in rapamycin-treated crickets compared to controls **(K)** This effect was present in both males and females. **(L)** Peripheral freezing was increased in acarbose-treated individuals relative to controls. **(M)** This increase was driven by females; no treatment effects were detected in males (*P < 0.05, \*\**P* < 0.01, \*\*\**P* < 0.001, \*\*\*\**P* < 0.0001).

For central-to-peripheral time ratios, rapamycin-treated males showed higher central occupancy (*d* = −0.80 [95% CI: −1.71, 0.11], *P* < 0.05), while acarbose-treated females trended higher than controls (Figs. 5C-D). All groups spent less time centrally than juveniles, with the sharpest reductions in males; compared to adults, historical geriatrics, and pooled controls, central times were only higher in rapamycin-treated males, while acarbose-treated groups trended higher against historical geriatrics.

Rapamycin altered locomotor dynamics, lowering central speed versus controls (*d* = 1.06 [95% CI: 0.39, 1.73], *P* < 0.05), an effect concentrated in females (Figure 5E-F). Compared to juveniles, central-to-peripheral speed ratios were higher in controls and phenylbutyrate-treated crickets, particularly females. Relative to adults, rapamycin-treated females displayed lower ratios; historical comparisons showed phenylbutyrate-treated females trending higher.

Freezing behavior showed the clearest drug effect. Rapamycin-treated crickets froze more than controls across sexes (*d*’s = −2.57 to −1.03, *P*’s < 0.05), with phenylbutyrate- and acarbose trending higher (Figs. 5G-H). Versus juveniles and adults, rapamycin-treated crickets of both sexes froze more; against historical geriatrics, all medicated groups froze more, with female effects restricted to rapamycin. Pooled analysis confirmed increases for all treatments, with rapamycin females and all treated males elevated.

Central-zone freezing was higher only in rapamycin-treated crickets (*d*’s = −1.33 to −1.31, *P*’s < 0.05) (Figure 5I-J) and preserved at juvenile-like levels in both sexes, while acarbose-treated females froze less than juvenile females, with controls trending lower. Compared to adults, freezing was lower in control and acarbose females. Historical geriatrics and pooled analyses showed higher freezing primarily in rapamycin-treated males.

Peripheral freezing increased only in acarbose-treated crickets (*d* = −0.76 [95% CI: –1.41, –0.11], *P* < 0.05), though rapamycin-treated females trended higher (Figs. 5K-L). Versus juveniles, acarbose- and phenylbutyrate-treated crickets froze more in the periphery, while rapamycin females trended higher. Versus adults, peripheral freezing was elevated in acarbose-, rapamycin-females, and phenylbutyrate-treated males. Historical geriatric comparisons confirmed higher peripheral freezing in acarbose- and phenylbutyrate-treated groups, with rapamycin showing only a trend; effects were largely male-specific. In pooled analyses, acarbose- (both sexes) and phenylbutyrate-treated males froze more than controls.

Complete statistics are provided in Appendix 5.

### Rapamycin consistently preserves induced locomotor performance in a sex-dependent manner

Across all analyses, rapamycin and phenylbutyrate-treated individuals ran longer than controls (*d*’s = −2.64 to −0.87, *P*’s < 0.05) (Figs. 6A-B). Both treatments restored performance to adult/mid-age-like levels and showed longer running durations than historical geriatric and pooled controls, with consistent trends across sexes.

**Figure 6.**
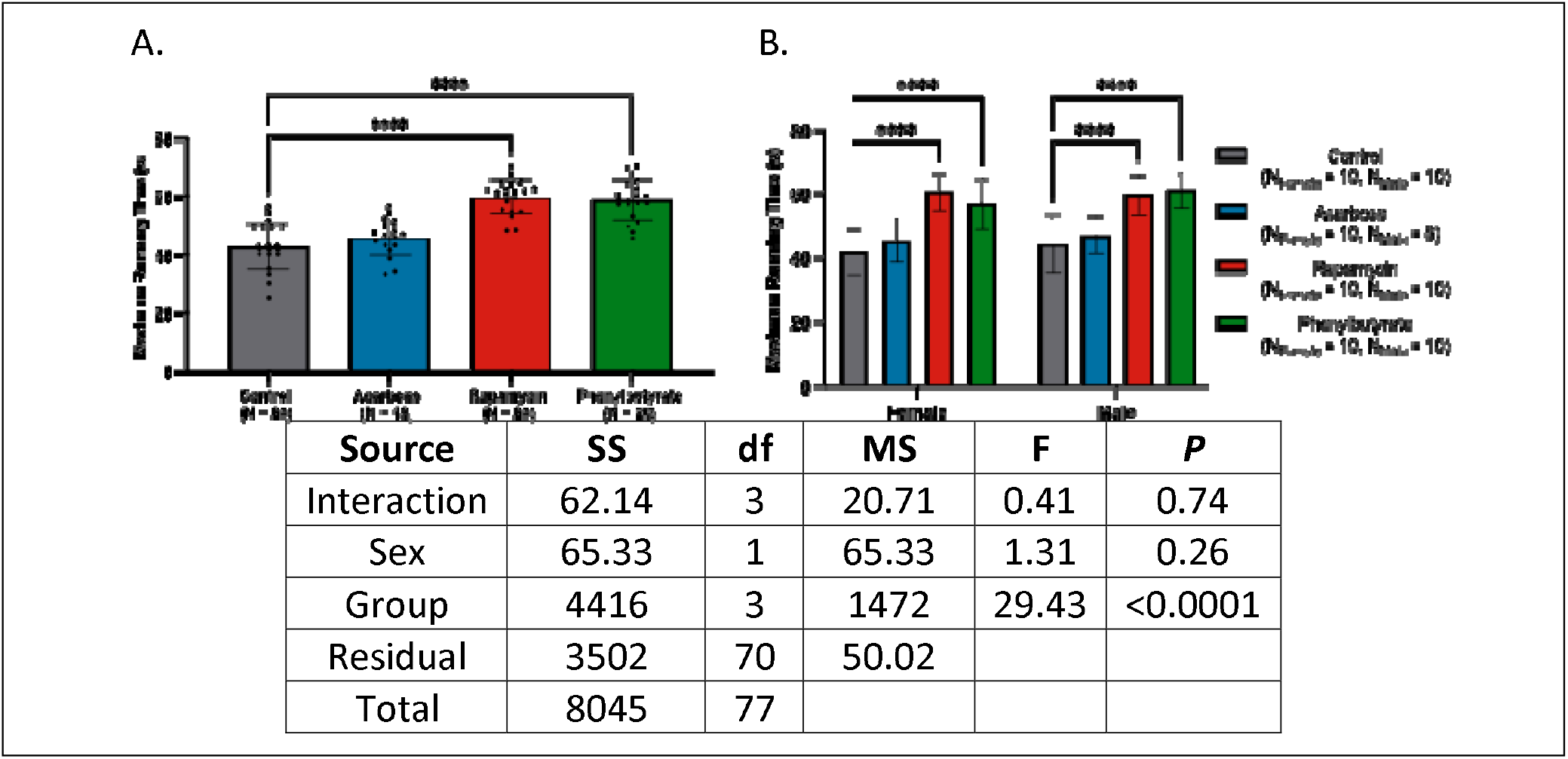
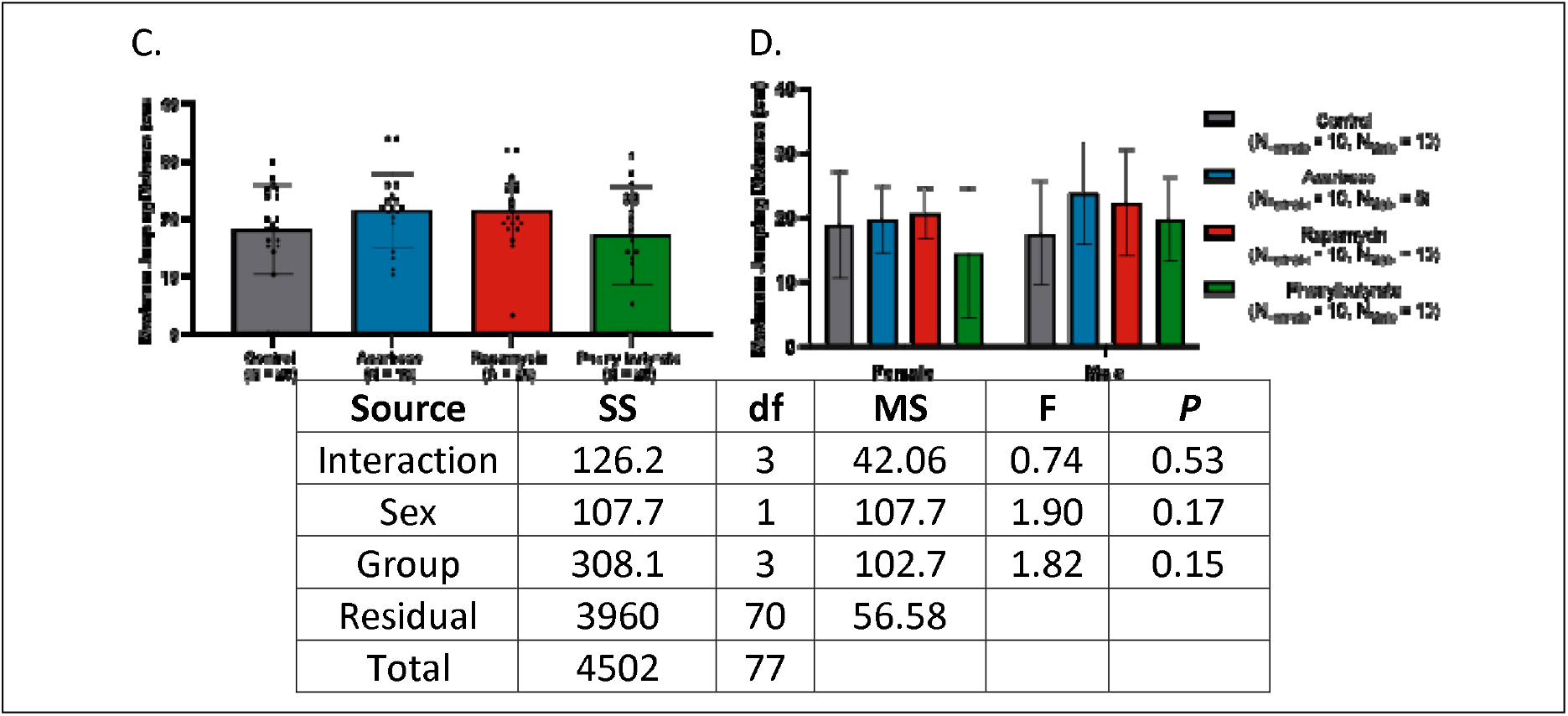
Physical performance across treatment groups. **(A)** Crickets treated with rapamycin or phenylbutyrate exhibited longer running durations than controls. **(B)** This enhancement was evident in both sexes. **(C)** Maximum jumping distance was comparable across treatment groups. **(D)** No sex-specific differences in jumping performance were detected (\*\*\*\**P* < 0.0001).

In contrast, maximum jumping distance did not differ among treatment groups (*d*’s = −0.43 to 0.13, *P*’s > 0.05) (Figs 6C-D). While comparisons with younger cohorts revealed no enhancements, rapamycin-treated females and acarbose-treated males jumped further than historical geriatric and pooled control counterparts, with rapamycin-treated males trending higher. No other treatment-sex combinations differed from controls.

Complete statistics are provided in Appendix 6.

### Intermittent treatment promotes weight gain in females

Percent weight change was estimated as the difference between individual final weight and mean baseline weight of the respective treatment cohort in the absence of longitudinal tracking. While only acarbose-treated females gained more weight than control females (*d* = 0.75 [95% CI: 0.07, 1.42], *P* < 0.05), females gained more weight than males across all medicated groups (*d*’s = −1.05 to −0.80, P’s < 0.05) (Fig. 7). Detailed summary statistics can be found in Appendix 7.

**Figure 7.**
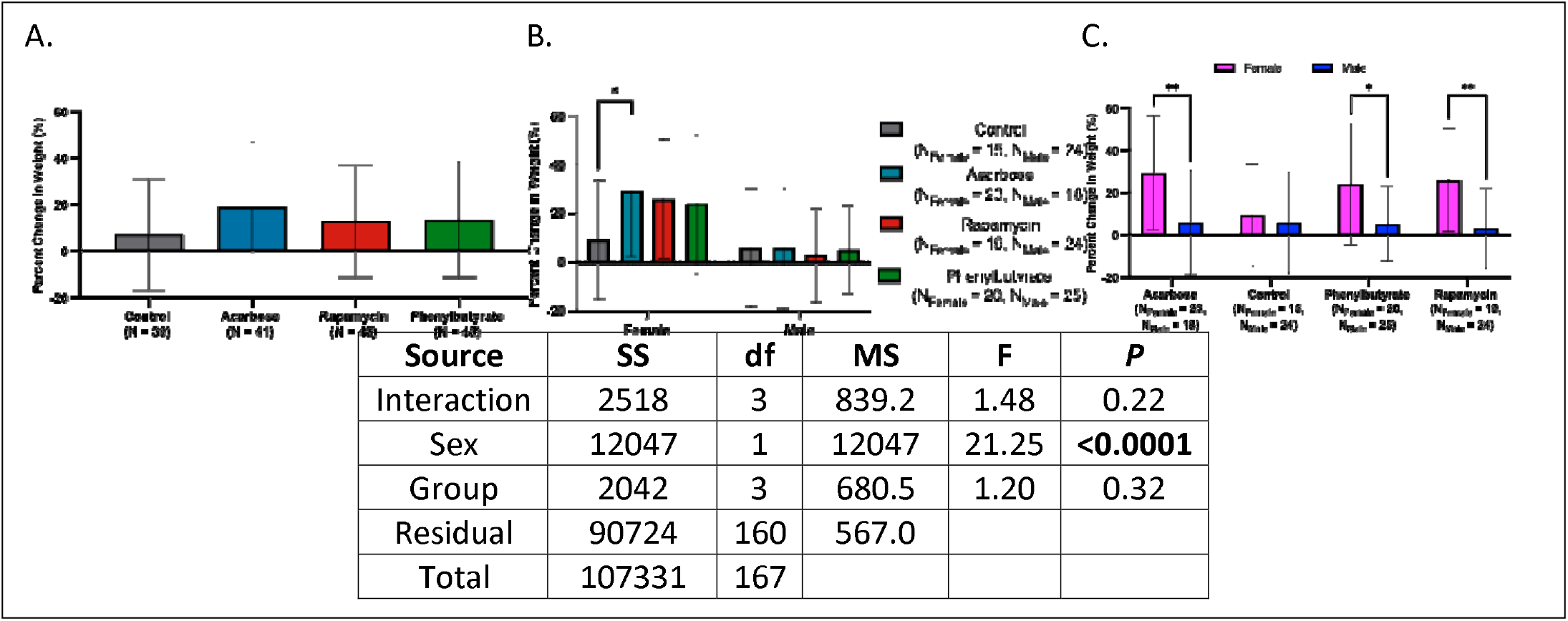
Percent weight change following treatment across groups and sexes. **(A)** Percent weight change was comparable across treatment groups. **(B)** Sex-stratified analysis revealed increased weight change only in acarbose-treated females. **(C)** Within-group sex comparisons showed greater weight change in females than males in the acarbose, rapamycin, and phenylbutyrate groups (*P < 0.05, **P < 0.01).

### Intermittent rapamycin extends, while acarbose shortens post-treatment lifespan in females

Rapamycin-treated individuals survived longer than controls (*HR*: 0.42 [95% CI: 0.25, 0.73], P < 0.001) while acarbose was associated with shortened lifespan (*HR*’s = 2.30 to 2.51, *P*’s = < 0.05), and phenylbutyrate-treated individuals had no survival advantage (Fig 8A). When stratified by sex, rapamycin-treated males survived longer (*HR*: 0.40 [95% CI: 0.20, 0.81], *P* < 0.01) and acarbose-treated females continued to show reduced survival (*HR*: = 3.03 [95% CI: 0.81, 11.4], *P* < 0.01) (Figs. 8B-C). Within-sex comparisons indicated only phenylbutyrate-treated females outlived their male counterparts (*HR*: 3.22 [95% CI: 1.08, 9.65], *P* < 0.01) (Appendix 7).

**Figure 8.**
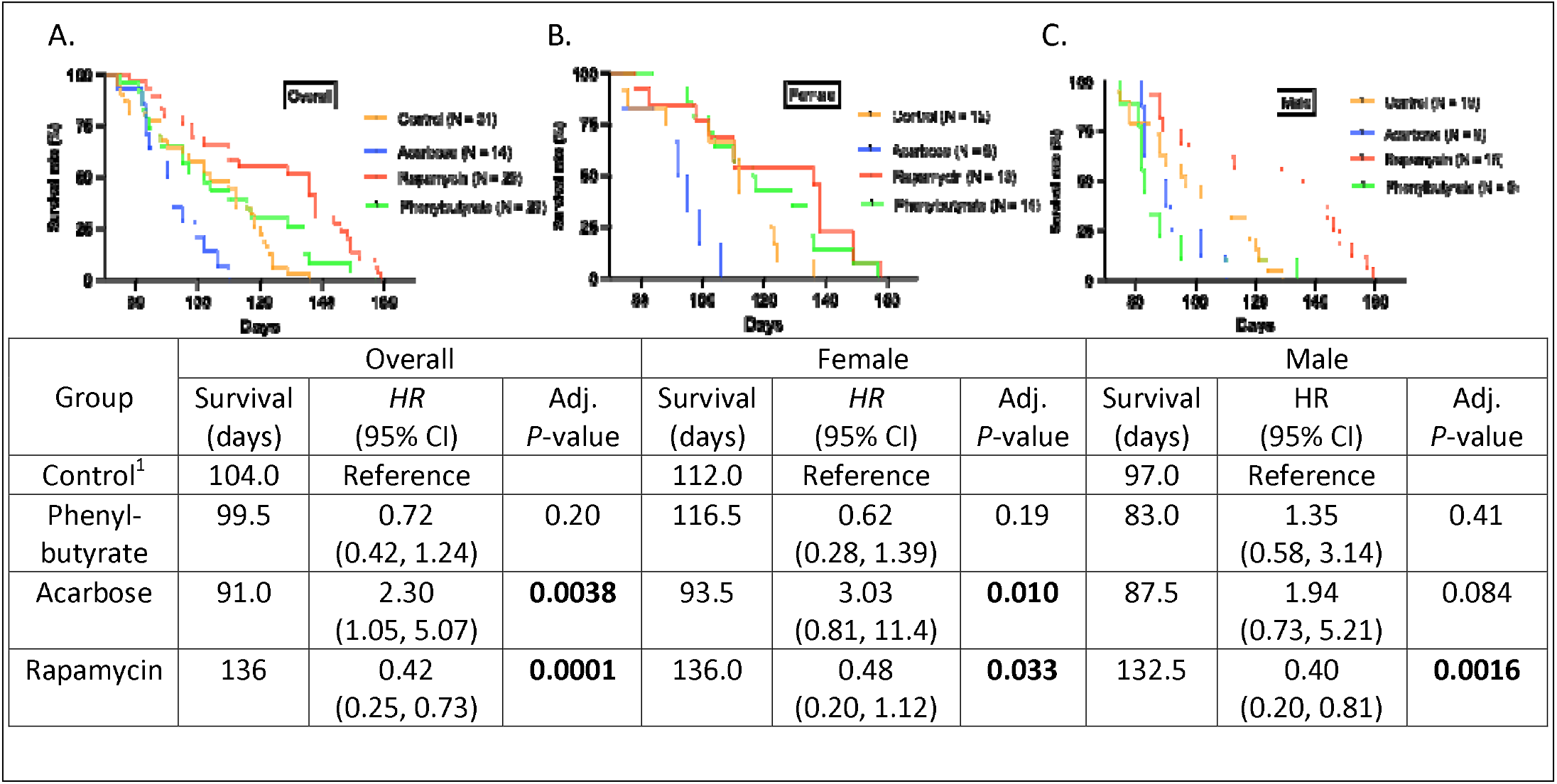
Survival curves of treatment groups. Kaplan-Meier survival curves illustrating the median survival of crickets across control, phenylbutyrate, acarbose, and rapamycin treatment groups in the **(A)** overall, **(B)** female, and **(C)** male populations. All treatment groups showed extended survival compared to the control group in the overall population while phenylbutyrate and rapamycin treated females showed extended survival compared to the control group.

## Discussion

This study establishes the house cricket (*Acheta domesticus*) as a tractable invertebrate model for aging biology, capable of recapitulating hallmark phenotypes including lifespan modulation, cognitive decline, and locomotor impairment. Each geroprotective compound tested (rapamycin, acarbose, and phenylbutyrate) extended lifespan, and select treatments preserved cognitive and physical function in late life. By integrating survival, behavioral, and physiological endpoints, we provide a multidimensional portrait of aging in an invertebrate system with strong translational potential.

### Integrated preservation of cognitive, locomotor, and motivational function through conserved pathways

Across behavioral domains, geroprotective interventions produced discrete yet overlapping benefits, revealing both shared and pathway-specific mechanisms of resilience. In odor-guided decision-making, rapamycin and phenylbutyrate restored food-odor preference to near-juvenile levels, with acarbose producing modest improvement. These effects are consistent with mTOR inhibition reducing neuroinflammation and enhancing autophagy [39–41] and HDAC inhibition stabilizing synaptic architecture [41–42]. The conservation of these effects across phyla suggests that mushroom bodies, central to insect olfactory learning [30, 43], may serve as a key substrate of pharmacologic action.

Exploratory behavior was similarly modulated in a pathway-specific manner. In the open field assay, rapamycin uniquely increased central-zone engagement in aged crickets, an effect largely male-driven and consistent with patterns linking physical robustness to greater environmental engagement in aging humans [44–45]. Phenylbutyrate and acarbose did not alter central occupancy, indicating that only certain pathways, such as those regulated by mTOR, may effectively modulate spatial strategy and motivational drive. Notably, rapamycin-treated crickets also exhibited increased freezing episodes, likely reflecting active pauses for environmental scanning rather than frailty-related immobility, paralleling exploratory “checks” described in invertebrates [46]. These findings reinforce the utility of open field analysis as a multidimensional tool capable of detecting changes in both locomotor vigor and motivational state and highlight the potential to pharmacologically target anxiety-like or avoidant behaviors as modifiable components of late-life frailty.

Locomotor assays revealed further domain-specific plasticity. Rapamycin and phenylbutyrate restored maximal running time to near-youthful levels, consistent with preserved neuromuscular coordination and metabolic efficiency, paralleling rodent findings on mTOR inhibition maintaining muscle integrity into late life [47] and HDAC inhibition enhancing oxidative stress resistance. Rapamycin also preserved walking speed and exploratory vigor, particularly in males. In contrast, jump performance, reflecting maximal power output, remained largely refractory to all treatments, suggesting that structural or biomechanical deficits in late life may be less amenable to systemic metabolic modulation. These distinctions mirror mammalian data showing that endurance- and strength-related traits are often modified via distinct, non-overlapping pathways [48–49]. Together, the behavioral assays (olfactory discrimination, open field exploration, treadmill running, and jump testing) captured separable yet interdependent aspects of functional aging, demonstrating that improvements in one domain do not necessarily generalize to others.

### Durability and sex-specificity of intermittent geroprotective treatment

Intermittent drug administration further revealed compound-specific durability of benefit. Brief rapamycin exposure conferred a persistent survival advantage long after treatment cessation, echoing murine findings where short-term mTORC1 inhibition remodels long-term aging trajectories [50]. In contrast, phenylbutyrate withdrawal yielded no net change, and intermittent acarbose reduced lifespan, particularly in females, suggesting that benefits from these agents may not persist, or may even reverse, post-treatment. These differences were not attributable to altered caloric intake (colony-level food consumption remained stable; unpublished observations), arguing against dietary restriction as the primary mechanism.

Sex-specific trajectories added further complexity. Rapamycin-treated males retained extended survival post-treatment, acarbose-treated females showed reduced longevity relative to controls, and phenylbutyrate-treated females outlived their male counterparts. These patterns partially parallel mammalian data in which females respond more robustly to rapamycin and phenylbutyrate, and males to acarbose [3, 39, 51–52]. Morphometric and physiological differences between sexes, such as heavier body mass in females [9], may influence pharmacokinetics, endocrine signaling, or energetic allocation, shaping drug responsiveness.

Importantly, intermittent treatment outcomes were linked to changes in somatic maintenance. Acarbose-treated females gained more weight than controls, and females across all treatment groups exhibited greater weight gain than males during the dosing period, potentially reflecting improved metabolic resilience or sex-specific allocation of resources. Notably, this somatic gain did not shorten lifespan, suggesting that body mass preservation and longevity were not inversely coupled. Instead, increased somatic investment in females may confer enhanced robustness under pharmacologic intervention, consistent with reports that sex differences in insect lifespan are highly context-dependent [53–54].

### Limitations and future directions

While these findings establish fundamental links between pharmacology and aging phenotypes in crickets, several limitations remain. First, sample sizes for some behavioral assays may have limited power to detect modest drug effects, and so future studies should incorporate simulation-based power calculations, such as Monte Carlo methods [55]. This would allow us to model expected variability and determine optimal sample sizes for detecting moderate treatment effects. Second, locomotor metrics captured gross performance but not fine gait parameters, which could reveal early neuromuscular decline [56]. Emerging methods in insect biomechanics [57] and computer vision-based kinematic tracking [58] would make this feasible, revealing not just how far, but *how* aged crickets move, offering more granular insight into locomotor aging and sarcopenia. Third, tissue-level validation, such as histological scoring of mushroom bodies, musculature, and other organs, will be critical to link functional outcomes with cellular integrity. Development of a standardized geropathology framework for crickets, modeled on murine systems [59–60], would enable such integration and facilitate cross-species comparison. Lastly, future work should test additional interventions (e.g., metformin, NAD^+^ precursors, SGLT2 inhibitors) [61] and combinatorial regimens [22] or explore early-life or “pulse” treatments [62]. Mapping the full trajectory of behavioral, physiological, and histological aging, and determining how each can be modulated, will move the field toward a unified, multidimensional definition of healthspan.

## Conclusion

By demonstrating that interventions known to extend mammalian lifespan can also preserve physiological and cognitive function in crickets, our findings identify conserved mechanisms of aging and intervention response. The house cricket offers a scalable, low-cost, and experimentally flexible platform for geroscience, complementing traditional mammalian systems and enabling rapid preclinical screening of candidate gerotherapeutics. Integrating functional, anatomical, and molecular endpoints will further enhance its translational relevance, bridging fundamental biology with strategies to extend healthy human life.

