## Supplemental Files for "The gerotherapeutic drugs rapamycin, acarbose, and phenylbutyrate extend lifespan and enhance healthy aging in house crickets"

**Appendix 1.**

| Experiment | Duration | Group | N | Sex | | RR (95% CI) | *P*-Value |
| --- | --- | --- | --- | --- | --- | --- | --- |
|  |  |  |  | Female^1^  [N (%)] | Male  [N (%)] |  |  |
| Scent Preference Test | 04/23/25 | Control^1^ | 19 | 5 (26.3) | 14 (73.7) |  |  |
|  |  | Acarbose | 20 | 10 (50.0) | 10 (50.0) | 0.57 (0.25, 1.16) | 0.13 |
|  |  | Rapamycin | 20 | 9 (45.0) | 11 (55.0) | 0.64 (0.28, 1.28) | 0.22 |
|  |  | Phenylbutyrate | 20 | 10 (50.0) | 10 (50.0) | 0.57 (0.25, 1.16) | 0.13 |
| Open Field Test | 02/27/25 | Control^1^ | 19 | 9 (47.4) | 10 (52.6) | 0.95 (0.49, 1.81) | 0.95 |
|  |  | Acarbose | 20 | 10 (50.0) | 10 (50.0) |  |  |
|  |  | Rapamycin | 20 | 10 (50.0) | 10 (50.0) |  |  |
|  |  | Phenylbutyrate | 20 | 10 (50.0) | 10 (50.0) |  |  |
| Treadmill Assay | 02/28/25 | Control^1^ | 20 | 10 (50.0) | 10 (50.0) |  |  |
|  |  | Acarbose | 18 | 10 (55.6) | 8 (44.4) | 0.90 (0.49, 1.67) | 0.73 |
|  |  | Rapamycin | 20 | 10 (50.0) | 10 (50.0) | 1.00 (0.53, 1.88) | >0.99 |
|  |  | Phenylbutyrate | 20 | 10 (50.0) | 10 (50.0) | 1.00 (0.53, 1.88) | >0.99 |

**Table S1. Experimental cohorts demonstrated balanced sex distributions across all lifespan and behavioral assays conducted between June 2024 and April 2025. Sample sizes (N), sex distributions, relative risk (RR) estimates with 95% confidence intervals (CI), and p-values for comparisons of male representation across experimental groups. ^1^Control females served as the reference for effect size estimation and statistical testing. RR = relative risk; CI = confidence interval; *P*-values calculated using Fisher’s exact test or chi-square test, as appropriate.**

| Overall | Control  (N = 44) | Acarbose  (N = 48) | | Rapamycin  (N = 46) | | Phenylbutyrate  (N = 48) | |
| --- | --- | --- | --- | --- | --- | --- | --- |
|  | Mean (SD) | Mean (SD) | *P* | Mean (SD) | *P* | Mean (SD) | *P* |
| Weight (g) | 0.51 (0.12) | 0.55 (0.11) | 0.12 | 0.49 (0.08) | 0.64 | 0.47 (0.07) | 0.12 |
| Body Length (cm) | 2.09 (0.21) | 2.12 (0.19) | 0.78 | 2.07 (0.17) | 0.92 | 2.02 (0.18) | 0.18 |
| Antennal Length (cm) | 2.45 (0.73) | 2.38 (0.53) | 0.90 | 2.61 (0.56) | 0.44 | 2.58 (0.56) | 0.59 |
| Hind Leg Length (cm) | 1.02 (0.08) | 1.00 (0.08) | 0.48 | 1.02 (0.08) | >0.99 | 0.99 (0.08) | 0.18 |
| Femoral Cross-Sectional Area (cm^2^) | 0.03 (0.01) | 0.03 (0.01) | >0.99 | 0.03 (0.01) | >0.99 | 0.03 (0.01) | >0.99 |
| Femoral Volume (cm^3^) | 0.01 (0.00) | 0.01 (0.00) | >0.99 | 0.01 (0.00) | >0.99 | 0.01 (0.00) | >0.99 |
| Femoral Surface Area to Volume | 28.89 (2.31) | 29.49 (1.95) | 0.43 | 28.91 (2.35) | >0.99 | 29.99 (2.38) | 0.05 |
| Hind Leg to Body Length | 0.49 (0.05) | 0.47 (0.05) | 0.14 | 0.50 (0.05) | 0.66 | 0.48 (0.05) | 0.65 |
| Antennal to Body Length | 1.18 (0.28) | 1.30 (0.22) | 0.05 | 1.26 (0.23) | 0.28 | 1.30 (0.25) | 0.05 |
| Hind Leg Length to Weight | 2.25 (0.48) | 2.09 (0.48) | 0.22 | 2.10 (0.45) | 0.27 | 2.17 (0.41) | 0.73 |
| Body Length to Weight | 4.33 (0.92) | 4.16 (0.89) | 0.64 | 4.24 (0.81) | 0.92 | 4.32 (0.74) | >0.99 |
| Hind Leg to Antennal Length | 0.77 (0.52) | 0.67 (0.17) | 0.28 | 0.74 (0.23) | 0.94 | 0.71 (0.21) | 0.67 |
| Antennal Length to Weight | 5.12 (2.05) | 5.42 (1.76) | 0.76 | 5.57 (1.65) | 0.49 | 5.68 (1.69) | 0.31 |
| Female | Control  (N = 17) | Acarbose  (N = 27) | | Rapamycin  (N = 19) | | Phenylbutyrate  (N = 23) | |
|  | Mean (SD) | Mean (SD) | *P* | Mean (SD) | *P* | Mean (SD) | *P* |
| Weight (g) | 0.51 (0.12) | 0.55 (0.11) | 0.48 | 0.54 (0.11) | 0.72 | 0.51 (0.11) | >0.99 |
| Body Length (cm) | 2.09 (0.21) | 2.20 (0.19) | 0.21 | 2.13 (0.17) | 0.89 | 2.06 (0.23) | 0.91 |
| Antennal Length (cm) | 2.45 (0.73) | 2.83 (0.53) | 0.06 | 2.77 (0.33) | 0.18 | 2.63 (0.53) | 0.57 |
| Hind Leg Length (cm) | 1.68 (0.58) | 1.89 (0.52) | 0.46 | 1.74 (0.57) | 0.98 | 1.76 (0.56) | 0.94 |
| Femoral Cross-Sectional Area (cm^2^) | 0.03 (0.00) | 0.03 (0.00) | 0.76 | 0.02 (0.00) | 0.62 | 0.03 (0.01) | 0.92 |
| Femoral Volume (cm^3^) | 0.01 (0.00) | 0.01 (0.00) | >0.99 | 0.01 (0.00) | >0.99 | 0.01 (0.00) | >0.99 |
| Femoral Surface Area to Volume | 27.95 (2.99) | 27.32 (1.99) | 0.78 | 28.14 (2.95) | 0.99 | 27.79 (3.04) | >0.99 |
| Hind Leg to Body Length | 0.80 (0.27) | 0.86 (0.24) | 0.78 | 0.81 (0.25) | >0.99 | 0.86 (0.27) | 0.83 |
| Antennal to Body Length | 1.18 (0.34) | 1.30 (0.26) | 0.29 | 1.30 (0.16) | 0.30 | 1.28 (0.24) | 0.40 |
| Hind Leg Length to Weight | 3.53 (1.65) | 3.57 (1.19) | >0.99 | 3.29 (1.09) | 0.90 | 3.63 (1.45) | 0.99 |
| Body Length to Weight | 4.33 (0.92) | 4.16 (0.89) | 0.81 | 4.07 (0.53) | 0.62 | 4.21 (0.80) | 0.93 |
| Hind Leg to Antennal Length | 0.77 (0.52) | 0.67 (0.17) | 0.49 | 0.62 (0.17) | 0.23 | 0.67 (0.16) | 0.47 |
| Antennal Length to Weight | 5.12 (2.05) | 5.42 (1.76) | 0.86 | 5.29 (0.93) | 0.97 | 5.41 (1.46) | 0.88 |
| Male | Control  (N = 27) | Acarbose  (N = 21) | | Rapamycin  (N = 27) | | Phenylbutyrate  (N = 25) | |
|  | Mean (SD) | Mean (SD) | *P* | Mean (SD) | *P* | Mean (SD) | *P* |
| Weight (g) | 0.37 (0.08) | 0.37 (0.08) | >0.99 | 0.36 (0.06) | 0.92 | 0.37 (0.06) | >0.99 |
| Body Length (cm) | 2.01 (0.20) | 2.01 (0.14) | >0.99 | 1.97 (0.12) | 0.65 | 2.02 (0.13) | 0.99 |
| Antennal Length (cm) | 2.57 (0.63) | 2.70 (0.46) | 0.74 | 2.60 (0.50) | 0.99 | 2.78 (0.50) | 0.36 |
| Hind Leg Length (cm) | 0.95 (0.12) | 0.95 (0.08) | >0.99 | 0.97 (0.08) | 0.76 | 0.97 (0.07) | 0.77 |
| Femoral Cross-Sectional Area (cm^2^) | 0.02 (0.00) | 0.02 (0.00) | >0.99 | 0.02 (0.00) | >0.99 | 0.02 (0.00) | >0.99 |
| Femoral Volume (cm^3^) | 0.01 (0.00) | 0.01 (0.00) | >0.99 | 0.01 (0.00) | >0.99 | 0.01 (0.00) | >0.99 |
| Femoral Surface Area to Volume | 28.08 (2.39) | 28.04 (2.39) | >0.99 | 28.24 (2.55) | 0.99 | 28.00 (1.78) | >0.99 |
| Hind Leg to Body Length | 0.87 (0.24) | 0.86 (0.24) | >0.99 | 0.92 (0.26) | 0.81 | 0.89 (0.25) | 0.98 |
| Antennal to Body Length | 1.28 (0.29) | 1.34 (0.21) | 0.74 | 1.32 (0.24) | 0.88 | 1.37 (0.22) | 0.41 |
| Hind Leg Length to Weight | 4.95 (1.53) | 4.87 (1.44) | >0.99 | 5.20 (1.83) | 0.91 | 4.98 (1.81) | >0.99 |
| Body Length to Weight | 5.78 (1.78) | 5.74 (1.49) | >0.99 | 5.55 (0.78) | 0.86 | 5.56 (1.04) | 0.88 |
| Hind Leg to Antennal Length | 0.73 (0.35) | 0.64 (0.15) | 0.56 | 0.74 (0.35) | >0.99 | 0.65 (0.17) | 0.61 |
| Antennal Length to Weight | 7.21 (1.92) | 7.68 (2.14) | 0.73 | 7.35 (1.82) | 0.99 | 7.58 (1.64) | 0.82 |

**Table S2. Morphological measurements across treatment groups.** Body measurements were assessed in control and treatment groups overall, and separately by sex. Between-group comparisons were conducted using one-way analysis of variance (ANOVA) followed by pairwise comparisons approximating Dunnett’s test (control vs. treatment comparisons only). *P*-values were adjusted for multiple comparisons across treatments within each measurement and Cohen’s *d* with Hedges’ *g* correction were calculated as the effect size with control as the reference group along with 95% confidence intervals obtained. No statistically significant differences were observed at α = 0.05 unless otherwise indicated.

|  | Control | | | | Acarbose | | | |
| --- | --- | --- | --- | --- | --- | --- | --- | --- |
|  | Mean (SD) | | *d*  (95% CI) | *P* | Mean (SD) | | *d*  (95% CI) | *P* |
|  | Female | Male |  |  | Female | Male |  |  |
| Weight (g) | 0.51 (0.12) | 0.37 (0.08) | -1.47  (-2.23, -0.71) | **0.00026** | 0.55 (0.11) | 0.37 (0.08) | 1.80  (1.13, 2.48) | **<0.0001** |
| Body Length (cm) | 2.09 (0.21) | 2.01 (0.20) | -0.13  (-0.87, 0.61) | 0.22 | 2.20 (0.19) | 2.01 (0.14) | 1.10  (0.49, 1.71) | **0.00024** |
| Antennal Length (cm) | 2.45 (0.73) | 2.57 (0.63) | -0.15  (-0.89, 0.59) | 0.58 | 2.83 (0.53) | 2.7 (0.46) | 0.26  (-0.32, 0.83) | 0.37 |
| Hind Leg Length (cm) | 1.68 (0.58) | 0.95 (0.12) | -1.95  (-2.72, -1.17) | **<0.0001** | 1.89 (0.52) | 0.95 (0.08) | 2.34  (1.61, 3.08) | **<0.0001** |
| Femoral Cross-Sectional Area (cm^2^) | 0.03 (0.00) | 0.02 (0.00) | 0.27  (-0.47, 1.01) | >0.99 | 0.03 (0.00) | 0.02 (0.00) | 0.00  (0.00, 0.00) | >0.99 |
| Femoral Volume (cm^3^) | 0.01 (0.00) | 0.01 (0.00) | 0.00  (0.00, 0.00) | >0.99 | 0.01 (0.00) | 0.01 (0.00) | 0.00  (0.00, 0.00) | >0.99 |
| Femoral Surface Area to Volume | 27.95 (2.99) | 28.08 (2.39) | -0.05  (-0.79, 0.69) | 0.88 | 27.32 (1.99) | 28.04 (2.39) | -0.33  (-0.90, 0.25) | 0.27 |
| Hind Leg to Body Length | 0.80 (0.27) | 0.87 (0.24) | -0.27  (-1.01, 0.47) | 0.38 | 0.86 (0.24) | 0.86 (0.24) | 0.00  (-0.57, 0.57) | >0.99 |
| Antennal to Body Length | 1.18 (0.34) | 1.28 (0.29) | -0.27  (-1.01, 0.47) | 0.32 | 1.30 (0.26) | 1.34 (0.21) | -0.16  (-0.74, 0.41) | 0.56 |
| Hind Leg Length to Weight | 3.53 (1.65) | 4.95 (1.53) | -0.79  (-1.54, -0.03) | **0.0074** | 3.57 (1.19) | 4.87 (1.44) | -0.98  (-1.58, -0.38) | **0.0019** |
| Body Length to Weight | 4.33 (0.92) | 5.78 (1.78) | -0.83  (-1.59, -0.08) | **0.0010** | 4.16 (0.89) | 5.74 (1.49) | -1.31  (-1.93, -0.68) | **0.00016** |
| Hind Leg to Antennal Length | 0.77 (0.52) | 0.73 (0.35) | 0.09  (-0.65, 0.83) | 0.78 | 0.67 (0.17) | 0.64 (0.15) | 0.18  (-0.39, 0.75) | 0.52 |
| Antennal Length to Weight | 5.12 (2.05) | 7.21 (1.92) | -0.92  (-1.67, -0.16) | **0.0019** | 5.42 (1.76) | 7.68 (2.14) | -1.15  (-1.76, -0.53) | **0.00036** |
|  | Rapamycin | | | | Phenylbutyrate | | | |
|  | Mean (SD) | | *d*  (95% CI) | *P* | Mean (SD) | | *d*  (95% CI) | *P* |
|  | Female | Male |  |  | Female | Male |  |  |
| Weight (g) | 0.54 (0.11) | 0.36 (0.06) | 2.10  (1.37, 2.84) | **<0.0001** | 0.51 (0.11) | 0.37 (0.06) | 1.57  (0.93, 2.22) | **<0.0001** |
| Body Length (cm) | 2.13 (0.17) | 1.97 (0.12) | 1.10  (0.47, 1.73) | **0.0014** | 2.06 (0.23) | 2.02 (0.13) | 0.21  (-0.35, 0.78) | 0.47 |
| Antennal Length (cm) | 2.77 (0.33) | 2.60 (0.50) | 0.38  (-0.21, 0.97) | 0.17 | 2.63 (0.53) | 2.78 (0.50) | -0.29  (-0.86, 0.28) | 0.32 |
| Hind Leg Length (cm) | 1.74 (0.57) | 0.97 (0.08) | 2.05  (1.32, 2.77) | **<0.0001** | 1.76 (0.56) | 0.97 (0.07) | 1.99  (1.30, 2.68) | **<0.0001** |
| Femoral Cross-Sectional Area (cm^2^) | 0.02 (0.00) | 0.02 (0.00) | 0.00  (0.00, 0.00) | >0.99 | 0.03 (0.01) | 0.02 (0.00) | 1.42  (0.79, 2.06) | **<0.0001** |
| Femoral Volume (cm^3^) | 0.01 (0.00) | 0.01 (0.00) | 0.00  (0.00, 0.00) | >0.99 | 0.01 (0.00) | 0.01 (0.00) | 0.00  (0.00, 0.00) | >0.99 |
| Femoral Surface Area to Volume | 28.14 (2.95) | 28.24 (2.55) | -0.04  (-0.62, 0.55) | 0.91 | 27.79 (3.04) | 28.0 (1.78) | -0.08  (-0.65, 0.48) | 0.77 |
| Hind Leg to Body Length | 0.81 (0.25) | 0.92 (0.26) | -0.42  (-1.02, 0.17) | 0.16 | 0.86 (0.27) | 0.89 (0.25) | -0.11  (-0.68, 0.45) | 0.69 |
| Antennal to Body Length | 1.30 (0.16) | 1.32 (0.24) | -0.09  (-0.68, 0.49) | 0.74 | 1.28 (0.24) | 1.37 (0.22) | -0.39  (-0.96, 0.19) | 0.18 |
| Hind Leg Length to Weight | 3.29 (1.09) | 5.20 (1.83) | -1.20  (-1.83, -0.56) | **<0.0001** | 3.63 (1.45) | 4.98 (1.81) | -0.81  (-1.39, -0.22) | **0.0064** |
| Body Length to Weight | 4.07 (0.53) | 5.55 (0.78) | -2.11  (-2.85, -1.38) | **<0.0001** | 4.21 (0.80) | 5.56 (1.04) | -1.42  (-2.06, -0.79) | **<0.0001** |
| Hind Leg to Antennal Length | 0.62 (0.17) | 0.74 (0.35) | -0.41  (-1.00, 0.19) | 0.13 | 0.67 (0.16) | 0.65 (0.17) | 0.12  (-0.45, 0.69) | 0.68 |
| Antennal Length to Weight | 5.29 (0.93) | 7.35 (1.82) | -1.33  (-1.98, -0.68) | **<0.0001** | 5.41 (1.46) | 7.58 (1.64) | -1.37  (-2.00, -0.74) | **<0.0001** |

**Table S3. Sex-specific differences within treatment groups for morphological measurements.** Female and male crickets were compared within each treatment group (control, acarbose, rapamycin, and phenylbutyrate). Sex-specific comparisons were performed using Welch’s t-tests (allowing unequal variances). Effect sizes were estimated with Cohen’s *d* using bias-corrected Hedges’ *g*, with 95% confidence intervals with females as the reference. *P*-values were Bonferroni-adjusted within each treatment group.

**Appendix 2. Comparative analysis of scent preference across pharmacological treatments.**

| **Percentage of Vanilla Arm Entries (%)** | | | | | | | | | |
| --- | --- | --- | --- | --- | --- | --- | --- | --- | --- |
|  | Overall | | | Female | | | Male | | |
|  | Mean (SD) | *d*  (95% CI) | *P* | Mean (SD) | *d*  (95% CI) | *P* | Mean (SD) | *d*  (95% CI) | *P* |
| Control  (N_Female_ = 5, N_Male_ = 14) | 43.42 (13.42) |  |  | 47.50 (18.54) |  |  | 41.96 (11.61) |  |  |
| Acarbose  (N_Female_ = 10, N_Male_ = 10) | 63.13  (14.89) | -1.36  (-2.06, -0.66) | **0.0007** | 60.00  (15.37) | -0.72  (-1.82, 0.39) | 0.20 | 66.25  (14.49) | -1.82  (-2.78, -0.86) | **0.0001** |
| Rapamycin  (N_Female_ = 9, N_Male_ = 11) | 68.13  (13.13) | -1.82  (-2.57, -1.08) | **<0.0001** | 68.06  (11.02) | -1.38  (-2.58, -0.17) | **0.019** | 68.18  (15.17) | -1.91  (-2.86, -0.96) | **<0.0001** |
| Phenylbutyrate  (N_Female_ = 10, N_Male_ = 10) | 60.00  (11.89) | -1.28  (-1.97, -0.59) | **0.0032** | 56.25  (10.62) | -0.61  (-1.70, 0.49) | 0.45 | 63.75  (12.43) | -1.76  (-2.71, -0.81) | **0.0006** |

**Table S1. Behavioral outcomes for the Y-maze scent preference assay across pharmacological treatment groups.** Mean percentages of entries into the vanilla-scented arm with corresponding standard deviations (SD) are reported for control, acarbose-, rapamycin-, and phenylbutyrate-treated crickets, both overall and stratified by sex (N = number of individuals per group). Pairwise treatment effects relative to controls were quantified using Cohen’s *d* with Hedges’ *g* correction, presented with 95% confidence intervals (CIs) in the format (lower, upper). Adjusted *P*-values were derived from Dunnett’s multiple comparisons post-hoc test.

| A. B. 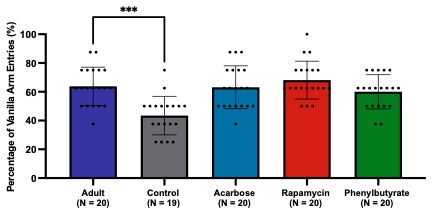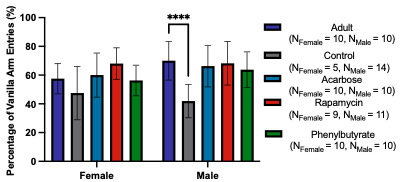 **Source** \| **SS** \| **df** \| **MS** \| **F** \| ***P***; Interaction \| 844.9 \| 4 \| 211.2 \| 1.22 \| 0.31; Sex \| 404.6 \| 1 \| 404.6 \| 2.33 \| 0.13; Group \| 5160 \| 4 \| 1290 \| 7.43 \| <0.0001; Residual \| 15447 \| 89 \| 173.6 \| \|; Total \| 21857 \| 98 \| \| \| | | | | | | | | |
| --- | --- | --- | --- | --- | --- | --- | --- | --- |
|  | Adult-Control | | Adult-Acarbose | | Adult-Rapamycin | | Adult-Phenylbutyrate | |
|  | *d*  (95% CI) | Adj. *P*-Value | *d*  (95% CI) | Adj. *P*-Value | *d*  (95% CI) | Adj. *P*-Value | *d*  (95% CI) | Adj. *P*-Value |
| Overall | 1.49  (0.78, 2.19) | **0.0003** | 0.04  (-0.58, 0.66) | >0.99 | -0.32  (-0.95, 0.30) | >0.99 | 0.29  (-0.33, 0.91) | >0.99 |
| Female | 0.70  (-0.41, 1.80) | 0.46 | -0.18  (-1.06, 0.70) | 0.98 | -0.94  (-1.88, 0.01) | 0.26 | 0.11  (-0.76, 0.99) | >0.99 |
| Male | 2.18  (1.16, 3.20) | **<0.0001** | 0.26  (-0.62, 1.14) | 0.92 | 0.12  (-0.74, 0.98) | 0.99 | 0.46  (-0.43, 1.35) | 0.66 |

**Figure S1. Pairwise comparisons of scent preference between adults and treatment groups.** Adults were compared to control, acarbose-, rapamycin-, and phenylbutyrate-treated crickets across **(A)** overall and **(B)** sex-stratified cohorts. Two-way analysis of variance (ANOVA) results are shown above the comparison table, indicating the effects of treatment group, sex, and their interaction on vanilla arm entry percentage. Pairwise comparisons are quantified using Cohen’s *d* with Hedges’ *g* correction to account for small sample size bias, with effect sizes reported alongside 95% CIs in the format (lower, upper). Adults served as the reference group for all comparisons, and adjusted P-values were calculated using Dunnett’s multiple comparisons post-hoc test (****P* < 0.001, *****P* < 0.0001).

| A. B. 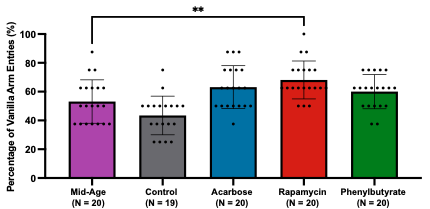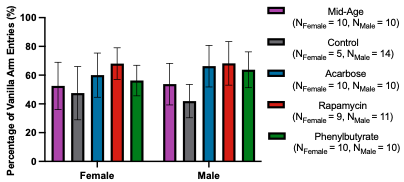 **Source** \| **SS** \| **df** \| **MS** \| **F** \| ***P***; Interaction \| 468.3 \| 4 \| 117.1 \| 0.61 \| 0.66; Sex \| 85.69 \| 1 \| 85.69 \| 0.44 \| 0.51; Group \| 5676 \| 4 \| 1419 \| 7.36 \| <0.0001; Residual \| 17151 \| 89 \| 192.7 \| \|; Total \| 23381 \| 98 \| \| \| | | | | | | | | |
| --- | --- | --- | --- | --- | --- | --- | --- | --- |
|  | Mid-Age-Control | | Mid-Age-Acarbose | | Mid-Age-Rapamycin | | Mid-Age-Phenylbutyrate | |
|  | *d*  (95% CI) | Adj. *P*-Value | *d*  (95% CI) | Adj. *P*-Value | *d*  (95% CI) | Adj. *P*-Value | *d*  (95% CI) | Adj. *P*-Value |
| Overall | 0.66  (0.02, 1.31) | 0.32 | -0.65  (-1.29, -0.02) | 0.18 | -1.04  (-1.70, -0.38) | **0.0087** | -0.50  (-1.12, 0.13) | 0.45 |
| Female | 0.27  (-0.80, 1.35) | 0.92 | -0.45  (-1.34, 0.44) | 0.58 | -1.05  (-2.01, -0.09) | 0.058 | -0.26  (-1.14, 0.62) | 0.94 |
| Male | 0.88  (0.04, 1.73) | 0.13 | -0.83  (-1.74, 0.09) | 0.14 | -0.93  (-1.83, -0.03) | 0.064 | -0.71  (-1.61, 0.19) | 0.31 |

**Figure S2. Comparative analysis of scent preference in mid-aged crickets relative to treatment groups.** The figure presents **(A)** overall and **(B)** sex-stratified comparisons between mid-aged crickets and those receiving control, acarbose, rapamycin, or phenylbutyrate treatments. Results from the two-way ANOVA are provided above the table, summarizing the main effects of treatment, sex, and their interaction on the percentage of vanilla arm entries. Pairwise differences are expressed as Cohen’s *d* with Hedges’ *g* correction to minimize small sample bias, with effect sizes accompanied by 95% CIs in the format (lower, upper). Mid-aged crickets were used as the reference group for all comparisons, and adjusted *P*-values were derived from Dunnett’s post-hoc test (***P* < 0.01).

| A. B. 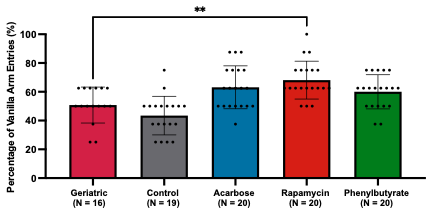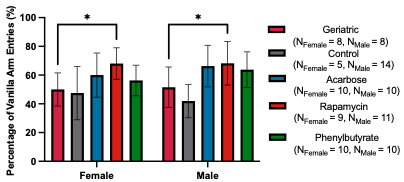 **Source** \| **SS** \| **df** \| **MS** \| **F** \| ***P***; Interaction \| 464.5 \| 4 \| 116.1 \| 0.65 \| 0.63; Sex \| 87.29 \| 1 \| 87.29 \| 0.49 \| 0.49; Group \| 6049 \| 4 \| 1512 \| 8.49 \| <0.0001; Residual \| 15147 \| 85 \| 178.2 \| \|; Total \| 21748 \| 94 \| \| \| | | | | | | | | |
| --- | --- | --- | --- | --- | --- | --- | --- | --- |
|  | Geriatric-Control | | Geriatric -Acarbose | | Geriatric-Rapamycin | | Geriatric-Phenylbutyrate | |
|  | *d*  (95% CI) | Adj. *P*-Value | *d*  (95% CI) | Adj. *P*-Value | *d*  (95% CI) | Adj. *P*-Value | *d*  (95% CI) | Adj. *P*-Value |
| Overall | 0.55  (-0.12, 1.23) | 0.60 | -0.87  (-1.56, -0.18) | 0.12 | -1.32  (-2.05, -0.60) | **0.0046** | -0.74  (-1.42, -0.06) | 0.28 |
| Female | 0.16  (-0.96, 1.28) | 0.99 | -0.69  (-1.64, 0.27) | 0.33 | -1.52  (-2.60, -0.44) | **0.023** | -0.54  (-1.49, 0.41) | 0.72 |
| Male | 0.74  (-0.16, 1.63) | 0.29 | -0.98  (-1.96, 0.01) | 0.071 | -1.08  (-2.05, -0.10) | **0.029** | -0.88  (-1.85, 0.09) | 0.16 |

**Figure S3. Comparison of scent preference between a historical geriatric cohort and treatment groups. (A)** Overall and **(B)** sex-stratified analyses are shown for historical geriatric crickets relative to control, acarbose-, rapamycin-, and phenylbutyrate-treated cohorts. Results from the two-way ANOVA are presented above the table, summarizing the effects of treatment group, sex, and their interaction on the percentage of vanilla arm entries. Pairwise comparisons were calculated using Cohen’s *d* with Hedges’ *g* correction to adjust for small sample size bias, with effect sizes reported alongside 95% CIs in the format (lower, upper). The historical geriatric cohort served as the reference group for all comparisons, and adjusted *P*-values were obtained using Dunnett’s multiple comparisons post-hoc test (**P* < 0.05, ***P* < 0.01).

| A. B. 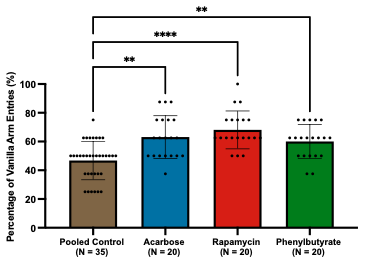 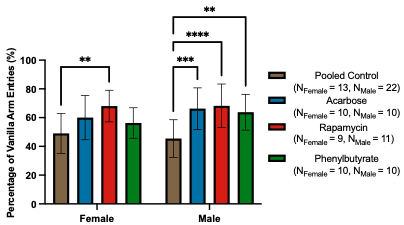 **Source** \| **SS** \| **df** \| **MS** \| **F** \| ***P***; Interaction \| 512.6 \| 3 \| 170.9 \| 0.95 \| 0.42; Sex \| 146.1 \| 1 \| 146.1 \| 0.81 \| 0.37; Group \| 6413 \| 3 \| 2138 \| 11.89 \| <0.0001; Residual \| 15636 \| 87 \| 179.7 \| \|; Total \| 22708 \| 94 \| \| \| | | | | | | |
| --- | --- | --- | --- | --- | --- | --- |
|  | Pooled Control-Acarbose | | Pooled Control-Rapamycin | | Pooled Control-Phenylbutyrate | |
|  | *d*  (95% CI) | Adj. *P*-Value | *d*  (95% CI) | Adj. *P*-Value | *d*  (95% CI) | Adj. *P*-Value |
| Overall | -1.18  (-1.77, -0.58) | **0.0012** | -1.61  (-2.24, -0.99) | **<0.0001** | -1.03  (-1.62, -0.45) | **0.0054** |
| Female | -0.74  (-1.52, 0.03) | 0.14 | -1.45  (-2.31, -0.58) | **0.0044** | -0.54  (-1.30, 0.23) | 0.45 |
| Male | -1.38  (-2.15, -0.60) | **0.0003** | -1.49  (-2.26, -0.73) | **<0.0001** | -1.24  (-2.01, -0.48) | **0.0017** |

**Figure S4. Comparison of scent preference between pooled controls and treatment groups. (A)** Overall and **(B)** sex-stratified analyses are presented for pooled control crickets relative to acarbose-, rapamycin-, and phenylbutyrate-treated cohorts. Results from the two-way ANOVA are shown above the table, summarizing the effects of treatment group, sex, and their interaction on the percentage of vanilla arm entries. Pairwise comparisons were quantified using Cohen’s *d* with Hedges’ *g* correction to account for small sample size bias, with effect sizes reported alongside 95% CIs in the format (lower, upper). Pooled controls served as the reference group for all comparisons, and adjusted P-values were obtained using Dunnett’s multiple comparisons post-hoc test (***P* < 0.01, ****P* < 0.001, *****P* < 0.0001).

**Appendix 3. OFT Measures of Locomotion.**

| **Total Distance (cm)** | | | | | | | | | |
| --- | --- | --- | --- | --- | --- | --- | --- | --- | --- |
|  | Overall | | | Female | | | Male | | |
|  | Mean (SD) | *d*  (95% CI) | *P* | Mean (SD) | *d*  (95% CI) | *P* | Mean (SD) | *d*  (95% CI) | *P* |
| Control  (N_Female_ = 9, N_Male_ = 10) | 2938 (885) |  |  | 2709 (874.9) |  |  | 3144 (887.7) |  |  |
| Acarbose  (N_Female_ = 10, N_Male_ = 10) | 2542 (925) | 0.43  (-0.21, 1.06) | 0.44 | 2400 (908.8) | 0.33  (−0.58, 1.24) | 0.82 | 2685 (966.8) | 0.47  (−0.42, 1.36) | 0.59 |
| Rapamycin  (N_Female_ = 10, N_Male_ = 10) | 3340 (1283) | -0.36  (-0.99, 0.28) | 0.43 | 2990 (948.7) | -0.29  (-1.20, 0.61) | 0.86 | 3689 (1518) | -0.42  (-1.31, 0.47) | 0.45 |
| Phenylbutyrate  (N_Female_ = 10, N_Male_ = 10) | 2245 (705) | 0.85  (0.20, 1.51) | 0.075 | 2393 (940.5) | 0.33  (-0.58, 1.24) | 0.82 | 2097 (338.6) | 1.49  (0.50, 2.48) | **0.049** |

**Table S1. Pairwise comparisons of total distance traveled between control and treatment groups.** Overall and sex-stratified analyses are shown for total distance traveled in control, acarbose-, rapamycin-, and phenylbutyrate-treated crickets. Effect sizes for each comparison were calculated using Cohen’s *d* with Hedges’ *g* correction to minimize small sample size bias, with values reported alongside 95% confidence intervals (CIs) in the format (lower, upper). Controls served as the reference group for all comparisons, and adjusted *P*-values were determined using Dunnett’s multiple comparisons post-hoc test.

| A. B. 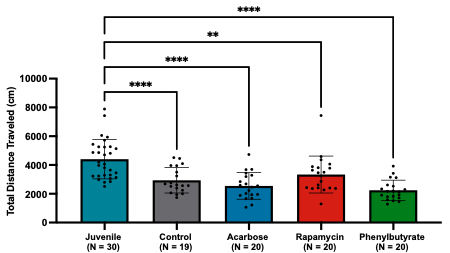 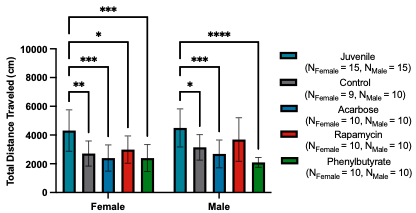 **Source** \| **SS** \| **df** \| **MS** \| **F** \| ***P***; Interaction \| 2.7e06 \| 4 \| 6.8e05 \| 0.55 \| 0.70; Sex \| 1.8e06 \| 1 \| 1.8e06 \| 1.48 \| 0.23; Group \| 7.2e07 \| 4 \| 1.8e07 \| 14.82 \| <0.0001; Residual \| 1.2e08 \| 99 \| 1.2e06 \| \|; Total \| 2.0e08 \| 108 \| \| \| | | | | | | | | |
| --- | --- | --- | --- | --- | --- | --- | --- | --- |
|  | Juvenile-Control | | Juvenile-Acarbose | | Juvenile-Rapamycin | | Juvenile-Phenylbutyrate | |
|  | *d*  (95% CI) | Adj. *P*-Value | *d*  (95% CI) | Adj. *P*-Value | *d*  (95% CI) | Adj. *P*-Value | *d*  (95% CI) | Adj. *P*-Value |
| Overall | 1.20  (0.58, 1.82) | **<0.0001** | 1.52  (0.88, 2.16) | **<0.0001** | 0.79  (0.20, 1.37) | **0.0041** | 1.85  (1.18, 2.52) | **<0.0001** |
| Female | 1.23  (0.33, 2.12) | **0.0032** | 1.47  (0.57, 2.37) | **0.0002** | 1.01  (0.16, 1.85) | **0.016** | 1.46  (0.57, 2.36) | **0.0002** |
| Male | 1.11  (0.25, 1.97) | **0.013** | 1.46  (0.56, 2.36) | **0.0004** | 0.56  (-0.26, 1.37) | 0.24 | 2.19  (1.19, 3.20) | **<0.0001** |

**Figure S1. Comparison of total distance traveled between juvenile crickets and treatment groups. (A)** Overall and **(B)** sex-stratified analyses are presented for juvenile crickets relative to control, acarbose-, rapamycin-, and phenylbutyrate-treated cohorts. Results from the two-way analysis of variance (ANOVA) are shown above the table, summarizing the effects of treatment group, sex, and their interaction on total distance traveled. Pairwise comparisons were quantified using Cohen’s *d* with Hedges’ *g* correction to account for small sample size bias, with effect sizes reported alongside 95% CIs in the format (lower, upper). Juvenile crickets served as the reference group for all comparisons, and adjusted *P*-values were obtained using Dunnett’s multiple comparisons post-hoc test (**P* < 0.05, ***P* < 0.01, ****P* < 0.001, *****P* < 0.0001).

| A. B. 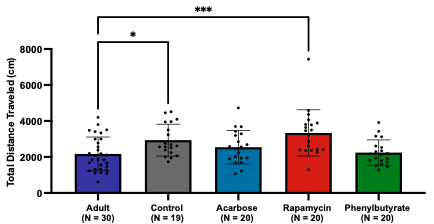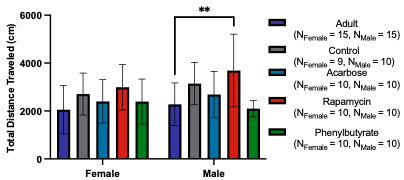 **Source** \| **SS** \| **df** \| **MS** \| **F** \| ***P***; Interaction \| 2.7e06 \| 4 \| 6.7e05 \| 0.72 \| 0.58; Sex \| 1.9e06 \| 1 \| 1.9e06 \| 2.06 \| 0.15; Group \| 2.1e07 \| 4 \| 5.3e06 \| 5.67 \| 0.0004; Residual \| 9.2e07 \| 99 \| 9.3e05 \| \|; Total \| 1.2e08 \| 108 \| \| \| | | | | | | | | |
| --- | --- | --- | --- | --- | --- | --- | --- | --- |
|  | Adult-Control | | Adult-Acarbose | | Adult-Rapamycin | | Adult-Phenylbutyrate | |
|  | *d*  (95% CI) | Adj. *P*-Value | *d*  (95% CI) | Adj. *P*-Value | *d*  (95% CI) | Adj. *P*-Value | *d*  (95% CI) | Adj. *P*-Value |
| Overall | -0.82  (-1.42, -0.23) | **0.028** | -0.39  (-0.96, 0.18) | 0.50 | -1.06  (-1.66, -0.46) | **0.0002** | -0.09  (-0.66, 0.48) | >0.99 |
| Female | -0.65  (-1.50, 0.19) | 0.33 | -0.34  (-1.15, 0.46) | 0.82 | -0.92  (-1.75, -0.08) | 0.069 | -0.33  (-1.14, 0.47) | 0.83 |
| Male | -0.94  (-1.78, -0.10) | 0.11 | -0.43  (-1.24, 0.38) | 0.72 | -1.16  (-2.02, -0.30) | **0.0021** | 0.24  (-0.56, 1.05) | 0.98 |

**Figure S2. Comparison of total distance traveled between adult crickets and treatment groups. (A)** Overall and **(B)** sex-stratified analyses are presented for adult crickets relative to control, acarbose-, rapamycin-, and phenylbutyrate-treated cohorts. Results from the two-way ANOVA are shown above the table, summarizing the effects of treatment group, sex, and their interaction on total distance traveled. Pairwise comparisons were quantified using Cohen’s *d* with Hedges’ *g* correction to account for small sample size bias, with effect sizes reported alongside 95% CIs in the format (lower, upper). Adult crickets served as the reference group for all comparisons, and adjusted *P*-values were obtained using Dunnett’s multiple comparisons post-hoc test (**P* < 0.05, ***P* < 0.01, ****P* < 0.001).

| A. B. 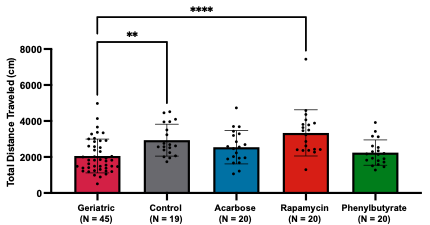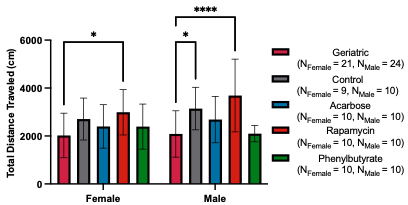 **Source** \| **SS** \| **df** \| **MS** \| **F** \| ***P***; Interaction \| 3.0e06 \| 4 \| 7.5e05 \| 0.80 \| 0.53; Sex \| 1.6e06 \| 1 \| 1.6e06 \| 1.68 \| 0.20; Group \| 2.8e07 \| 4 \| 6.9e06 \| 7.48 \| <0.0001; Residual \| 1.1e08 \| 114 \| 9.3e05 \| \|; Total \| 1.4e08 \| 123 \| \| \| | | | | | | | | |
| --- | --- | --- | --- | --- | --- | --- | --- | --- |
|  | Geriatric-Control | | Geriatric -Acarbose | | Geriatric-Rapamycin | | Geriatric-Phenylbutyrate | |
|  | *d*  (95% CI) | Adj. *P*-Value | *d*  (95% CI) | Adj. *P*-Value | *d*  (95% CI) | Adj. *P*-Value | *d*  (95% CI) | Adj. *P*-Value |
| Overall | -0.94  (-1.50, -0.38) | **0.0042** | -0.51  (-1.05, 0.02) | 0.21 | -1.20  (-1.77, -0.63) | **<0.0001** | -0.21  (-0.74, 0.32) | 0.90 |
| Female | -0.73  (-1.53, 0.07) | 0.25 | -0.40  (-1.16, 0.36) | 0.74 | -1.01  (-1.80, -0.21) | **0.038** | -0.39  (-1.14, 0.37) | 0.75 |
| Male | -1.09  (-1.87, -0.31) | **0.016** | -0.60  (-1.36, 0.15) | 0.32 | -1.36  (-2.17, -0.56) | **<0.0001** | -0.01  (-0.75, 0.73) | >0.99 |

**Figure S3. Comparison of total distance traveled between geriatric crickets and treatment groups. (A)** Overall and **(B)** sex-stratified analyses are presented for geriatric crickets relative to control, acarbose-, rapamycin-, and phenylbutyrate-treated cohorts. Results from the two-way ANOVA are shown above the table, summarizing the effects of treatment group, sex, and their interaction on total distance traveled. Pairwise comparisons were quantified using Cohen’s *d* with Hedges’ *g* correction to account for small sample size bias, with effect sizes reported alongside 95% CIs in the format (lower, upper). Geriatric crickets served as the reference group for all comparisons, and adjusted *P*-values were obtained using Dunnett’s multiple comparisons post-hoc test (**P* < 0.05, ***P* < 0.01, *****P* < 0.0001).

| A. B. 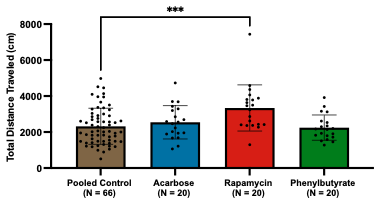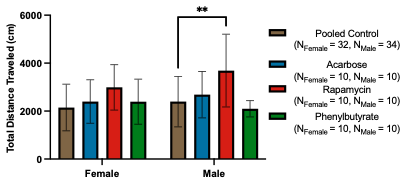 **Source** \| **SS** \| **df** \| **MS** \| **F** \| ***P***; Interaction \| 2.5e06 \| 3 \| 8.4e05 \| 0.82 \| 0.48; Sex \| 1.3e06 \| 1 \| 1.3e06 \| 1.30 \| 0.26; Group \| 1.9e07 \| 3 \| 6.3e06 \| 6.17 \| 0.0006; Residual \| 1.2e08 \| 118 \| 1.0e06 \| \|; Total \| 1.4e08 \| 125 \| \| \| | | | | | | |
| --- | --- | --- | --- | --- | --- | --- |
|  | Pooled Control-Acarbose | | Pooled Control-Rapamycin | | Pooled Control-Phenylbutyrate | |
|  | *d*  (95% CI) | Adj. *P*-Value | *d*  (95% CI) | Adj. *P*-Value | *d*  (95% CI) | Adj. *P*-Value |
| Overall | -0.23 (-0.73, 0.28) | 0.75 | -0.94 (-1.46, -0.42) | **0.0004** | 0.08 (-0.43, 0.58) | 0.99 |
| Female | -0.26 (-0.97, 0.46) | 0.86 | -0.85  (-1.58, -0.12) | 0.066 | -0.25 (-0.96, 0.46) | 0.87 |
| Male | -0.27 (-0.98, 0.43) | 0.80 | -1.09  (-1.83, -0.35) | **0.0016** | 0.31 (-0.40, 1.02) | 0.78 |

**Figure S4. Comparison of total distance traveled between pooled controls and treatment groups. (A)** Overall and **(B)** sex-stratified analyses are presented for pooled control crickets relative to acarbose-, rapamycin-, and phenylbutyrate-treated cohorts. Results from the two-way ANOVA are shown above the table, summarizing the effects of treatment group, sex, and their interaction on total distance traveled. Pairwise comparisons were quantified using Cohen’s *d* with Hedges’ *g* correction to account for small sample size bias, with effect sizes reported alongside 95% CIs in the format (lower, upper). Pooled controls served as the reference group for all comparisons, and adjusted *P*-values were obtained using Dunnett’s multiple comparisons post-hoc test (***P* < 0.01, ****P* < 0.001).

| **Average Speed (cm/s)** | | | | | | | | | |
| --- | --- | --- | --- | --- | --- | --- | --- | --- | --- |
|  | Overall | | | Female | | | Male | | |
|  | Mean (SD) | *d*  (95% CI) | *P* | Mean (SD) | *d*  (95% CI) | *P* | Mean (SD) | *d*  (95% CI) | *P* |
| Control  (N_Female_ = 9, N_Male_ = 10) | 9.34 (2.83) |  |  | 8.62 (2.72) |  |  | 9.98 (2.91) |  |  |
| Acarbose  (N_Female_ = 10, N_Male_ = 10) | 8.38 (3.05) | 0.32  (-0.31, 0.95) | 0.66 | 7.93 (3.05) | 0.23  (−0.68, 1.13) | 0.93 | 8.83 (3.13) | 0.36  (−0.52, 1.25) | 0.75 |
| Rapamycin  (N_Female_ = 10, N_Male_ = 10) | 11.02 (4.19) | -0.46  (-1.09, 0.18) | 0.23 | 9.91 (3.15) | -0.42  (-1.33, 0.49) | 0.69 | 12.1 (4.93) | -0.50  (-1.39, 0.39) | 0.30 |
| Phenylbutyrate  (N_Female_ = 10, N_Male_ = 10) | 7.39 (2.25) | 0.75  (0.10, 1.40) | 0.14 | 7.84 (3.00) | 0.26  (-0.64, 1.16) | 0.91 | 6.95 (1.12) | 1.32  (0.35, 2.28) | 0.089 |

**Table S2. Pairwise comparisons of average speed between control and treatment groups.** Overall and sex-specific analyses are presented for average speed in control, acarbose-, rapamycin-, and phenylbutyrate-treated crickets. Effect sizes for each comparison were computed using Cohen’s *d* with Hedges’ *g* correction to account for small sample size bias and are reported with 95% CIs in the format (lower, upper). Controls were used as the reference group for all comparisons, and adjusted *P*-values were obtained through Dunnett’s multiple comparisons post-hoc test.

| A. B. 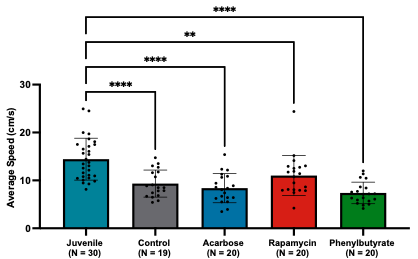 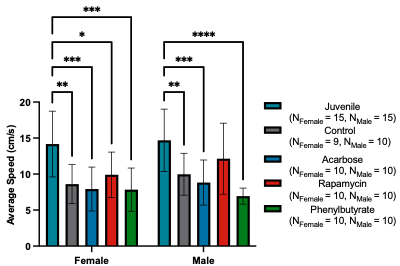 **Source** \| **SS** \| **df** \| **MS** \| **F** \| ***P***; Interaction \| 26.66 \| 4 \| 6.665 \| 0.52 \| 0.72; Sex \| 18.07 \| 1 \| 18.07 \| 1.42 \| 0.24; Group \| 780.7 \| 4 \| 195.2 \| 15.29 \| <0.0001; Residual \| 1263 \| 99 \| 12.76 \| \|; Total \| 2088 \| 108 \| \| \| | | | | | | | | |
| --- | --- | --- | --- | --- | --- | --- | --- | --- |
|  | Juvenile-Control | | Juvenile-Acarbose | | Juvenile-Rapamycin | | Juvenile-Phenylbutyrate | |
|  | *d*  (95% CI) | Adj. *P*-Value | *d*  (95% CI) | Adj. *P*-Value | *d*  (95% CI) | Adj. *P*-Value | *d*  (95% CI) | Adj. *P*-Value |
| Overall | 1.30  (0.67, 1.93) | **<0.0001** | 1.53  (0.89, 2.17) | **<0.0001** | 0.78  (0.20, 1.37) | **0.0046** | 1.88  (1.21, 2.56) | **<0.0001** |
| Female | 1.34  (0.43, 2.25) | **0.0014** | 1.49  (0.59, 2.39) | **0.0002** | 1.01  (0.16, 1.86) | **0.016** | 1.52  (0.62, 2.42) | **0.0001** |
| Male | 1.19  (0.32, 2.05) | **0.0064** | 1.45  (0.56, 2.35) | **0.0004** | 0.55  (-0.27, 1.36) | 0.26 | 2.17  (1.17, 3.17) | **<0.0001** |

**Figure S5. Comparison of average speed between juvenile crickets and treatment groups. (A)** Overall and **(B)** sex-stratified analyses are presented for juvenile crickets relative to control, acarbose-, rapamycin-, and phenylbutyrate-treated cohorts. Results from the two-way ANOVA are shown above the table, summarizing the effects of treatment group, sex, and their interaction on average speed. Pairwise comparisons were quantified using Cohen’s *d* with Hedges’ *g* correction to account for small sample size bias, with effect sizes reported alongside 95% CIs in the format (lower, upper). Juvenile crickets served as the reference group for all comparisons, and adjusted *P*-values were obtained using Dunnett’s multiple comparisons post-hoc test (**P* < 0.05, ***P* < 0.01, ****P* < 0.001, *****P* < 0.0001).

| A. B. 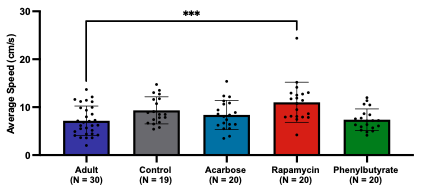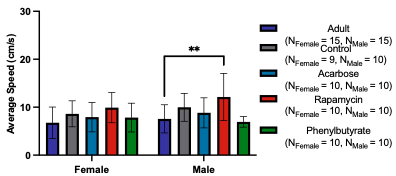 **Source** \| **SS** \| **df** \| **MS** \| **F** \| ***P***; Interaction \| 25.96 \| 4 \| 6.489 \| 0.66 \| 0.62; Sex \| 20.63 \| 1 \| 20.63 \| 2.09 \| 0.15; Group \| 217.8 \| 4 \| 54.45 \| 5.51 \| **0.0005**; Residual \| 977.7 \| 99 \| 9.876 \| \|; Total \| 1242 \| 108 \| \| \| | | | | | | | | |
| --- | --- | --- | --- | --- | --- | --- | --- | --- |
|  | Adult-Control | | Adult-Acarbose | | Adult-Rapamycin | | Adult-Phenylbutyrate | |
|  | *d*  (95% CI) | Adj. *P*-Value | *d*  (95% CI) | Adj. *P*-Value | *d*  (95% CI) | Adj. *P*-Value | *d*  (95% CI) | Adj. *P*-Value |
| Overall | -0.71  (-1.31, -0.12) | 0.071 | -0.39  (-0.96, 0.18) | 0.50 | -1.06  (-1.67, -0.46) | **0.0002** | -0.08  (-0.64, 0.49) | >0.99 |
| Female | -0.58  (-1.43, 0.26) | 0.46 | -0.36  (-1.16, 0.45) | 0.80 | -0.94  (-1.79, -0.10) | 0.057 | -0.33  (-1.14, 0.48) | 0.84 |
| Male | -0.80  (-1.63, 0.03) | 0.20 | -0.40  (-1.21, 0.40) | 0.75 | -1.14  (-2.00, -0.28) | **0.0022** | 0.25  (-0.55, 1.05) | 0.97 |

**Figure S6. Average speed comparisons between adult crickets and treatment groups. (A)** Overall and **(B)** sex-stratified analyses compare adult crickets to control, acarbose-, rapamycin-, and phenylbutyrate-treated cohorts. The two-way ANOVA results are provided above the table, outlining the effects of treatment, sex, and their interaction on average speed. Pairwise differences were calculated using Cohen’s *d* with Hedges’ *g* correction to reduce small sample size bias, with effect sizes shown alongside 95% CIs in the format (lower, upper). Adults were used as the reference group for all comparisons, and adjusted *P*-values were derived from Dunnett’s multiple comparisons post-hoc test (***P* < 0.01, ****P* < 0.001).

| A. B. 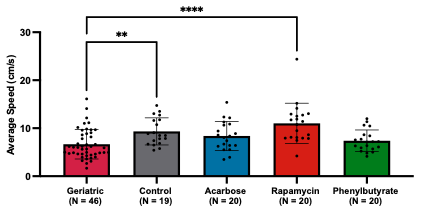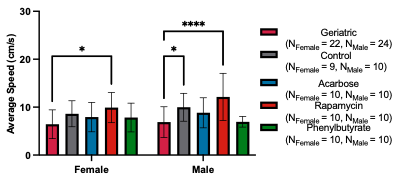 **Source** \| **SS** \| **df** \| **MS** \| **F** \| ***P***; Interaction \| 27.41 \| 4 \| 6.852 \| 0.70 \| 0.60; Sex \| 18.25 \| 1 \| 18.25 \| 1.85 \| 0.18; Group \| 305.3 \| 4 \| 76.32 \| 7.74 \| **<0.0001**; Residual \| 1134 \| 115 \| 9.858 \| \|; Total \| 1485 \| 124 \| \| \| | | | | | | | | |
| --- | --- | --- | --- | --- | --- | --- | --- | --- |
|  | Geriatric-Control | | Geriatric -Acarbose | | Geriatric-Rapamycin | | Geriatric-Phenylbutyrate | |
|  | *d*  (95% CI) | Adj. *P*-Value | *d*  (95% CI) | Adj. *P*-Value | *d*  (95% CI) | Adj. *P*-Value | *d*  (95% CI) | Adj. *P*-Value |
| Overall | -0.88  (-1.43, -0.32) | **0.0087** | -0.55  (-1.09, -0.01) | 0.15 | -1.24  (-1.81, -0.68) | **<0.0001** | -0.25  (-0.78, 0.28) | 0.84 |
| Female | -0.73  (-1.53, 0.07) | 0.27 | -0.49  (-1.25, 0.28) | 0.59 | -1.11  (-1.92, -0.31) | **0.017** | -0.46  (-1.22, 0.30) | 0.64 |
| Male | -0.97  (-1.74, -0.19) | **0.037** | -0.60  (-1.35, 0.15) | 0.32 | -1.35  (-2.16, -0.55) | **<0.0001** | -0.02  (-0.76, 0.71) | >0.99 |

**Figure S7. Average speed comparisons between historical geriatric crickets and treatment groups. (A)** Overall and **(B)** sex-stratified analyses are shown for historical geriatric crickets relative to control, acarbose-, rapamycin-, and phenylbutyrate-treated cohorts. Results from the two-way ANOVA are displayed above the table, summarizing the effects of treatment group, sex, and their interaction on average speed. Pairwise comparisons were determined using Cohen’s *d* with Hedges’ *g* correction to account for small sample size bias, with effect sizes reported alongside 95% CIs in the format (lower, upper). Historical geriatric crickets served as the reference group for all comparisons, and adjusted *P*-values were obtained using Dunnett’s multiple comparisons post-hoc test (**P* < 0.05, ***P* < 0.01, *****P* < 0.0001).

| A. B. 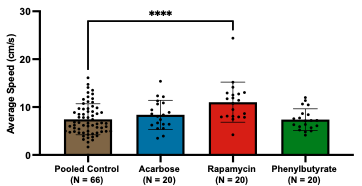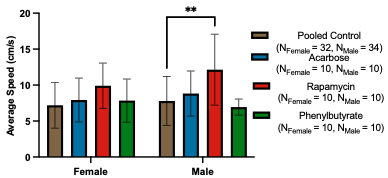 **Source** \| **SS** \| **df** \| **MS** \| **F** \| ***P***; Interaction \| 24.68 \| 3 \| 8.23 \| 0.76 \| 0.52; Sex \| 12.11 \| 1 \| 12.11 \| 1.13 \| 0.29; Group \| 204.1 \| 3 \| 68.05 \| 6.32 \| **0.0005**; Residual \| 1227 \| 114 \| 10.77 \| \|; Total \| 1468 \| 121 \| \| \| | | | | | | |
| --- | --- | --- | --- | --- | --- | --- |
|  | Pooled Control-Acarbose | | Pooled Control-Rapamycin | | Pooled Control-Phenylbutyrate | |
|  | *d*  (95% CI) | Adj. *P*-Value | *d*  (95% CI) | Adj. *P*-Value | *d*  (95% CI) | Adj. *P*-Value |
| Overall | -0.29 (-0.79, 0.21) | 0.59 | -1.02 (-1.54, -0.49) | **<0.0001** | 0.02 (-0.48, 0.52) | >0.99 |
| Female | -0.23 (-0.95, 0.49) | 0.90 | -0.84  (-1.59, -0.09) | 0.074 | -0.20 (-0.93, 0.52) | 0.93 |
| Male | -0.31 (-1.01, 0.40) | 0.75 | -1.13  (-1.87, -0.39) | **0.0011** | 0.27 (-0.44, 0.98) | 0.85 |

**Figure S8. Comparison of average speed between pooled controls and treatment groups. (A)** Overall and **(B)** sex-stratified analyses compare pooled control crickets with acarbose-, rapamycin-, and phenylbutyrate-treated cohorts. The results of the two-way ANOVA are provided above the table, detailing the effects of treatment, sex, and their interaction on average speed. Pairwise comparisons were computed using Cohen’s *d* with Hedges’ *g* correction to address small sample size bias, with effect sizes presented alongside 95% CIs in the format (lower, upper). Pooled controls were used as the reference group for all comparisons, and adjusted *P*-values were calculated using Dunnett’s multiple comparisons post-hoc test (***P* < 0.01, *****P* < 0.0001).

| **Walking/Running Distance** | | | | | | | | | |
| --- | --- | --- | --- | --- | --- | --- | --- | --- | --- |
|  | Overall | | | Female | | | Male | | |
|  | Mean (SD) | *d*  (95% CI) | *P* | Mean (SD) | *d*  (95% CI) | *P* | Mean (SD) | *d*  (95% CI) | *P* |
| Control  (N_Female_ = 9, N_Male_ = 10) | 2.19 (0.64) |  |  | 2.22 (0.82) |  |  | 2.16 (0.47) |  |  |
| Acarbose  (N_Female_ = 10, N_Male_ = 10) | 2.96 (1.25) | -0.75  (-1.40, -0.10) | **0.029** | 2.98 (1.27) | −0.67  (−1.60, 0.25) | 0.19 | 2.93 (1.29) | −0.76  (−1.67, 0.15) | 0.17 |
| Rapamycin  (N_Female_ = 10, N_Male_ = 10) | 2.29 (0.73) | -0.14  (-0.77, 0.49) | 0.98 | 2.23 (0.72) | -0.01  (-0.91, 0.89) | >0.99 | 2.34 (0.78) | -0.27  (-1.15, 0.61) | 0.95 |
| Phenylbutyrate  (N_Female_ = 10, N_Male_ = 10) | 2.69 (0.90) | -0.62  (-1.27, 0.02) | 0.21 | 2.57 (0.93) | -0.38  (-1.29, 0.53) | 0.76 | 2.82 (0.89) | -0.89  (-1.81, 0.03) | 0.28 |

**Table S3. Pairwise comparisons of walking-to-running distance ratio between control and treatment groups.** Overall and sex-stratified analyses are shown for walking-to-running distance ratios in control, acarbose-, rapamycin-, and phenylbutyrate-treated crickets. Effect sizes were calculated using Cohen’s *d* with Hedges’ *g* correction to reduce bias from small sample sizes and are presented with 95% CIs in the format (lower, upper). Controls served as the reference group for all comparisons, and adjusted *P*-values were calculated using Dunnett’s multiple comparisons post-hoc test.

| A. B. 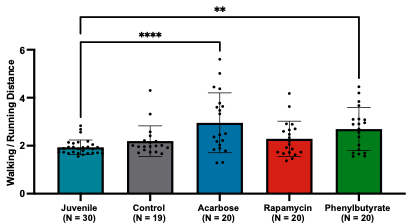 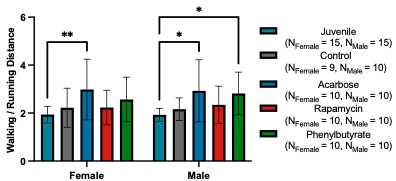 **Source** \| **SS** \| **df** \| **MS** \| **F** \| ***P***; Interaction \| 0.36 \| 4 \| 0.090 \| 0.14 \| 0.87; Sex \| 0.06 \| 1 \| 0.061 \| 0.09 \| 0.76; Group \| 15.41 \| 4 \| 3.85 \| 5.90 \| **0.0003**; Residual \| 64.63 \| 99 \| 0.65 \| \|; Total \| 80.46 \| 108 \| \| \| | | | | | | | | |
| --- | --- | --- | --- | --- | --- | --- | --- | --- |
|  | Juvenile-Control | | Juvenile-Acarbose | | Juvenile-Rapamycin | | Juvenile-Phenylbutyrate | |
|  | *d*  (95% CI) | Adj. *P*-Value | *d*  (95% CI) | Adj. *P*-Value | *d*  (95% CI) | Adj. *P*-Value | *d*  (95% CI) | Adj. *P*-Value |
| Overall | -0.55  (-1.14, 0.03) | 0.66 | -1.23  (-1.85, -0.62) | **<0.0001** | -0.68  (-1.26, -0.10) | 0.36 | -1.22  (-1.83, -0.60) | **0.0046** |
| Female | -0.48  (-1.31, 0.36) | 0.84 | -1.20  (-2.06, -0.33) | **0.0077** | -0.53  (-1.35, 0.28) | 0.81 | -0.95  (-1.79, -0.11) | 0.19 |
| Male | -0.61  (-1.43, 0.20) | 0.90 | -1.16  (-2.02, -0.30) | **0.012** | -0.75  (-1.57, 0.08) | 0.56 | -1.45  (-2.34, -0.55) | **0.030** |

**Figure S9. Walking and running distance comparisons between juvenile crickets and treatment groups. (A)** Overall and **(B)** sex-stratified analyses evaluate juvenile crickets in relation to control, acarbose-, rapamycin-, and phenylbutyrate-treated cohorts. Results from the two-way ANOVA are shown above the table, outlining the contributions of treatment, sex, and their interaction to walking and running distance. Pairwise differences were estimated using Cohen’s *d* with Hedges’ *g* correction to account for small sample size bias, with effect sizes listed alongside 95% CIs in the format (lower, upper). Juvenile crickets served as the reference group for all comparisons, and adjusted *P*-values were derived using Dunnett’s multiple comparisons post-hoc test (**P* < 0.05, ***P* < 0.01, *****P* < 0.0001).

| A. B. 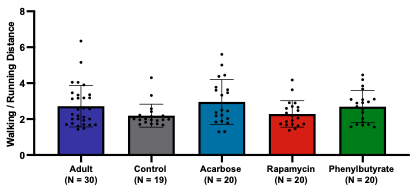 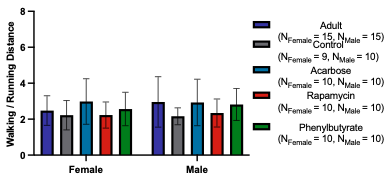 **Source** \| **SS** \| **df** \| **MS** \| **F** \| ***P***; Interaction \| 1.28 \| 4 \| 0.32 \| 0.32 \| 0.86; Sex \| 0.56 \| 1 \| 0.56 \| 0.56 \| 0.46; Group \| 8.20 \| 4 \| 2.05 \| 2.05 \| 0.09; Residual \| 99.05 \| 99 \| 1.00 \| \|; Total \| 109.09 \| 108 \| \| \| | | | | | | | | |
| --- | --- | --- | --- | --- | --- | --- | --- | --- |
|  | Adult-Control | | Adult-Acarbose | | Adult-Rapamycin | | Adult-Phenylbutyrate | |
|  | *d*  (95% CI) | Adj. *P*-Value | *d*  (95% CI) | Adj. *P*-Value | *d*  (95% CI) | Adj. *P*-Value | *d*  (95% CI) | Adj. *P*-Value |
| Overall | 0.52  (-0.06, 1.11) | 0.23 | -0.20  (-0.76, 0.37) | 0.84 | 0.42  (-0.15, 0.99) | 0.39 | 0.03  (-0.54, 0.59) | >0.99 |
| Female | 0.31  (-0.52, 1.14) | 0.94 | -0.47  (-1.28, 0.34) | 0.58 | 0.31  (-0.50, 1.11) | 0.94 | -0.10  (-0.90, 0.70) | >0.99 |
| Male | 0.68  (-0.14, 1.50) | 0.18 | 0.02  (-0.78, 0.82) | >0.99 | 0.50  (-0.31, 1.31) | 0.39 | 0.11  (-0.69, 0.91) | 0.99 |

**Figure S10. Walking and running distance comparisons between adult crickets and treatment groups. (A)** Overall and **(B)** sex-stratified analyses compare adult crickets to control, acarbose-, rapamycin-, and phenylbutyrate-treated cohorts. The results from the two-way ANOVA are provided above the table, summarizing the effects of treatment, sex, and their interaction on walking and running distance. Pairwise comparisons were calculated using Cohen’s *d* with Hedges’ *g* correction to correct for small sample size bias, with effect sizes reported together with 95% CIs in the format (lower, upper). Adults served as the reference group for all comparisons, and adjusted *P*-values were obtained through Dunnett’s multiple comparisons post-hoc test.

| A. B. 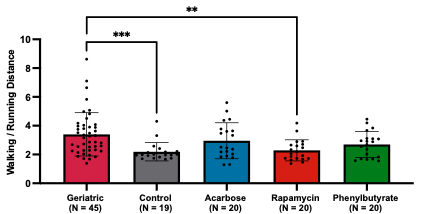 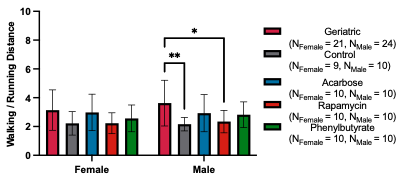 **Source** \| **SS** \| **df** \| **MS** \| **F** \| ***P***; Interaction \| 1.61 \| 4 \| 0.40 \| 0.29 \| 0.89; Sex \| 0.60 \| 1 \| 0.60 \| 0.43 \| 0.51; Group \| 28.01 \| 4 \| 7.00 \| 5.01 \| **0.0009**; Residual \| 159.2 \| 114 \| 1.40 \| \|; Total \| 189.42 \| 123 \| \| \| | | | | | | | | |
| --- | --- | --- | --- | --- | --- | --- | --- | --- |
|  | Geriatric-Control | | Geriatric -Acarbose | | Geriatric-Rapamycin | | Geriatric-Phenylbutyrate | |
|  | *d*  (95% CI) | Adj. *P*-Value | *d*  (95% CI) | Adj. *P*-Value | *d*  (95% CI) | Adj. *P*-Value | *d*  (95% CI) | Adj. *P*-Value |
| Overall | 0.91  (0.35, 1.47) | **0.0010** | 0.30  (-0.23, 0.83) | 0.47 | 0.83  (0.28, 1.37) | **0.0022** | 0.52  (-0.02, 1.05) | 0.094 |
| Female | 0.71  (-0.09, 1.51) | 0.18 | 0.11  (-0.64, 0.87) | 0.99 | 0.72  (-0.05, 1.49) | 0.16 | 0.44  (-0.32, 1.20) | 0.57 |
| Male | 1.05  (0.27, 1.83) | **0.0052** | 0.45  (-0.29, 1.20) | 0.37 | 0.89  (0.13, 1.66) | **0.018** | 0.55  (-0.20, 1.30) | 0.24 |

**Figure S11. Walking and running distance comparisons between geriatric crickets and treatment groups. (A)** Overall and **(B)** sex-stratified analyses are shown for geriatric crickets relative to control, acarbose-, rapamycin-, and phenylbutyrate-treated cohorts. The two-way ANOVA results are presented above the table, highlighting the effects of treatment, sex, and their interaction on walking and running distance. Pairwise comparisons were determined using Cohen’s *d* with Hedges’ *g* correction to adjust for small sample size bias, with effect sizes displayed alongside 95% CIs in the format (lower, upper). Geriatric crickets served as the reference group for all comparisons, and adjusted *P*-values were calculated using Dunnett’s multiple comparisons post-hoc test (**P* < 0.05, ***P* < 0.01, ****P* < 0.001).

| A. B. 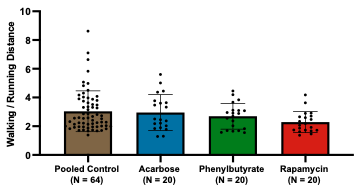 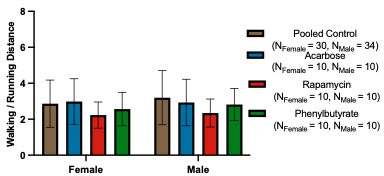 **Source** \| **SS** \| **df** \| **MS** \| **F** \| ***P***; Interaction \| 0.64 \| 3 \| 0.21 \| 0.14 \| 0.94; Sex \| 0.62 \| 1 \| 0.62 \| 0.40 \| 0.53; Group \| 9.14 \| 3 \| 3.05 \| 1.97 \| 0.12; Residual \| 179.7 \| 116 \| 1.55 \| \|; Total \| 190.1 \| 123 \| \| \| | | | | | | |
| --- | --- | --- | --- | --- | --- | --- |
|  | Pooled Control-Acarbose | | Pooled Control-Rapamycin | | Pooled Control-Phenylbutyrate | |
|  | *d*  (95% CI) | Adj. *P*-Value | *d*  (95% CI) | Adj. *P*-Value | *d*  (95% CI) | Adj. *P*-Value |
| Overall | 0.06 (-0.44, 0.57) | 0.99 | 0.57 (0.06, 1.08) | 0.053 | 0.26 (-0.24, 0.77) | 0.60 |
| Female | -0.12 (-0.83, 0.59) | 0.99 | -0.14 (-0.85, 0.57) | 0.40 | 0.20 (-0.51, 0.91) | 0.88 |
| Male | 0.18 (-0.53, 0.89) | 0.90 | 0.69 (-0.03, 1.41) | 0.16 | 0.27 (-0.44, 0.97) | 0.77 |

**Figure S12. Walking and running distance comparisons between pooled control crickets and treatment groups. (A)** Overall and **(B)** sex-stratified analyses present pooled controls in comparison with acarbose-, rapamycin-, and phenylbutyrate-treated cohorts. Results from the two-way ANOVA are shown above the table, summarizing the contributions of treatment group, sex, and their interaction to walking and running distance. Pairwise comparisons were computed using Cohen’s *d* with Hedges’ *g* correction to mitigate small sample size bias, with effect sizes reported together with 95% CIs in the format (lower, upper). Pooled controls were used as the reference group for all comparisons, and adjusted *P*-values were obtained through Dunnett’s multiple comparisons post-hoc test.

| **Walking/Running Time** | | | | | | | | | |
| --- | --- | --- | --- | --- | --- | --- | --- | --- | --- |
|  | Overall | | | Female | | | Male | | |
|  | Mean (SD) | *d*  (95% CI) | *P* | Mean (SD) | *d*  (95% CI) | *P* | Mean (SD) | *d*  (95% CI) | *P* |
| Control  (N_Female_ = 9, N_Male_ = 10) | 10.8 (4.6) |  |  | 11.7 (5.44) |  |  | 10.0 (3.68) |  |  |
| Acarbose  (N_Female_ = 10, N_Male_ = 10) | 13.8 (8.1) | −0.32  (−1.22, 0.59) | 0.38 | 14.28 (9.34) | −0.32  (−1.22, 0.59) | 0.75 | 13.33 (7.09) | −0.56  (−1.46, 0.33) | 0.58 |
| Rapamycin  (N_Female_ = 10, N_Male_ = 10) | 10.1 (5.0) | 0.39  (-0.52, 1.30) | 0.97 | 9.70 (4.37) | 0.39  (-0.52, 1.30) | 0.86 | 10.4 (5.75) | -0.08  (-0.96, 0.80) | >0.99 |
| Phenylbutyrate  (N_Female_ = 10, N_Male_ = 10) | 14.2 (8.5) | -0.35  (-1.26, 0.56) | 0.28 | 15.0 (11.23) | -0.35  (-1.26, 0.56) | 0.59 | 13.4 (4.85) | -0.76  (-1.66, 0.15) | 0.57 |

**Table S4. Pairwise comparisons of walking-to-running time ratio between control and treatment groups.** Overall and sex-stratified analyses are presented for walking-to-running time ratios in control, acarbose-, rapamycin-, and phenylbutyrate-treated crickets. Effect sizes for each comparison were computed using Cohen’s *d* with Hedges’ *g* correction to account for small sample size bias and are reported with 95% CIs in the format (lower, upper). Controls served as the reference group for all comparisons, and adjusted *P*-values were derived using Dunnett’s multiple comparisons post-hoc test.

| A. B. 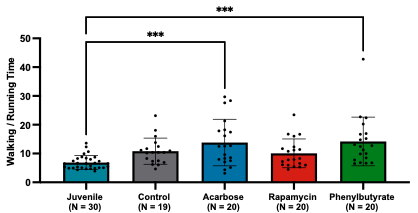 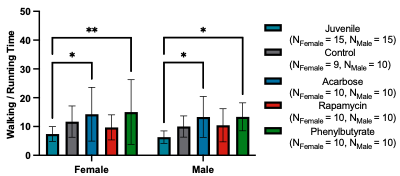 **Source** \| **SS** \| **df** \| **MS** \| **F** \| ***P***; Interaction \| 19.10 \| 4 \| 4.775 \| 0.13 \| 0.97; Sex \| 22.81 \| 1 \| 22.81 \| 0.63 \| 0.43; Group \| 888.4 \| 4 \| 222.1 \| 6.16 \| **0.0002**; Residual \| 3570 \| 99 \| 36.06 \| \|; Total \| 4500 \| 108 \| \| \| | | | | | | | | |
| --- | --- | --- | --- | --- | --- | --- | --- | --- |
|  | Juvenile-Control | | Juvenile-Acarbose | | Juvenile-Rapamycin | | Juvenile-Phenylbutyrate | |
|  | *d*  (95% CI) | Adj. *P*-Value | *d*  (95% CI) | Adj. *P*-Value | *d*  (95% CI) | Adj. *P*-Value | *d*  (95% CI) | Adj. *P*-Value |
| Overall | -1.14  (-1.75, -0.52) | 0.084 | -1.26  (-1.88, -0.64) | **0.0003** | -0.87  (-1.46, -0.28) | 0.20 | -1.28  (-1.89, -0.66) | **0.0001** |
| Female | -1.07  (-1.95, -0.19) | 0.29 | -1.08  (-1.93, -0.22) | **0.022** | -0.66  (-1.48, 0.16) | 0.78 | -1.01  (-1.85, -0.16) | **0.0092** |
| Male | -1.25  (-2.12, -0.38) | 0.39 | -1.43  (-2.32, -0.54) | **0.019** | -0.99  (-1.84, -0.15) | 0.30 | -1.97  (-2.94, -1.00) | **0.018** |

**Figure S13. Walking and running time comparisons between juvenile crickets and treatment groups. (A)** Overall and **(B)** sex-stratified analyses assess juvenile crickets relative to control, acarbose-, rapamycin-, and phenylbutyrate-treated cohorts. The two-way ANOVA results are displayed above the table, indicating the effects of treatment, sex, and their interaction on walking and running time. Pairwise differences were derived using Cohen’s *d* with Hedges’ *g* correction to minimize bias from small sample sizes, with effect sizes reported alongside 95% CIs in the format (lower, upper). Juvenile crickets served as the reference group for all comparisons, and adjusted *P*-values were calculated using Dunnett’s multiple comparisons post-hoc test (**P* < 0.05, ***P* < 0.01, ****P* < 0.001).

| A. B. 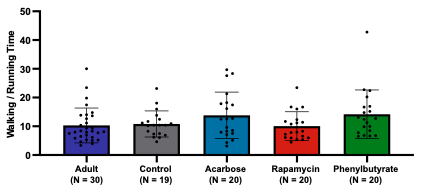 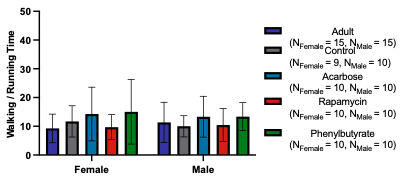 **Source** \| **SS** \| **df** \| **MS** \| **F** \| ***P***; Interaction \| 66.43 \| 4 \| 16.61 \| 0.37 \| 0.83; Sex \| 2.35 \| 1 \| 2.347 \| 0.05 \| 0.82; Group \| 340.0 \| 4 \| 84.99 \| 1.89 \| 0.12; Residual \| 4450 \| 99 \| 44.95 \| \|; Total \| 4859 \| 108 \| \| \| | | | | | | | | |
| --- | --- | --- | --- | --- | --- | --- | --- | --- |
|  | Adult-Control | | Adult-Acarbose | | Adult-Rapamycin | | Adult-Phenylbutyrate | |
|  | *d*  (95% CI) | Adj. *P*-Value | *d*  (95% CI) | Adj. *P*-Value | *d*  (95% CI) | Adj. *P*-Value | *d*  (95% CI) | Adj. *P*-Value |
| Overall | -0.09  (-0.67, 0.48) | >0.99 | -0.50  (-1.07, 0.08) | 0.22 | 0.03  (-0.53, 0.60) | >0.99 | -0.54  (-1.12, 0.04) | 0.14 |
| Female | -0.46  (-1.29, 0.38) | 0.82 | -0.69  (-1.52, 0.13) | 0.22 | -0.09  (-0.89, 0.71) | >0.99 | -0.69  (-1.51, 0.13) | 0.13 |
| Male | 0.22  (-0.59, 1.02) | 0.97 | -0.27  (-1.08, 0.53) | 0.89 | 0.14  (-0.66, 0.94) | 0.99 | -0.32  (-1.12, 0.49) | 0.89 |

**Figure S14. Walking and running time comparisons between adult crickets and treatment groups. (A)** Overall and **(B)** sex-stratified analyses compare adult crickets to control, acarbose-, rapamycin-, and phenylbutyrate-treated cohorts. Results from the two-way ANOVA are provided above the table, detailing the effects of treatment, sex, and their interaction on walking and running time. Pairwise comparisons were calculated using Cohen’s *d* with Hedges’ *g* correction to reduce bias associated with small sample sizes, with effect sizes presented alongside 95% CIs in the format (lower, upper). Adults served as the reference group for all comparisons, and adjusted *P*-values were determined using Dunnett’s multiple comparisons post-hoc test.

| A. B. 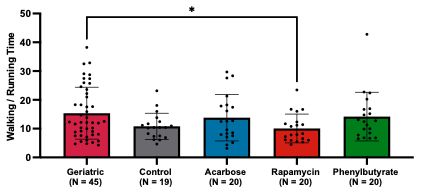 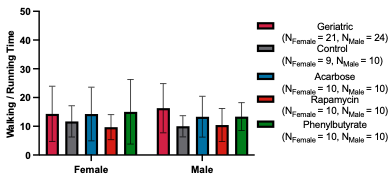 **Source** \| **SS** \| **df** \| **MS** \| **F** \| ***P***; Interaction \| 77.01 \| 4 \| 19.25 \| 0.32 \| 0.87; Sex \| 2.82 \| 1 \| 2.823 \| 0.05 \| 0.83; Group \| 524.4 \| 4 \| 131.1 \| 2.15 \| 0.08; Residual \| 6938 \| 114 \| 60.86 \| \|; Total \| 7542 \| 123 \| \| \| | | | | | | | | |
| --- | --- | --- | --- | --- | --- | --- | --- | --- |
|  | Geriatric-Control | | Geriatric -Acarbose | | Geriatric-Rapamycin | | Geriatric-Phenylbutyrate | |
|  | *d*  (95% CI) | Adj. *P*-Value | *d*  (95% CI) | Adj. *P*-Value | *d*  (95% CI) | Adj. *P*-Value | *d*  (95% CI) | Adj. *P*-Value |
| Overall | 0.57  (0.02, 1.11) | 0.11 | 0.18  (-0.35, 0.70) | 0.89 | 0.65  (0.11, 1.19) | **0.042** | 0.13  (-0.40, 0.66) | 0.96 |
| Female | 0.30  (-0.49, 1.08) | 0.85 | 0.00  (-0.75, 0.76) | >0.99 | 0.54  (-0.23, 1.30) | 0.39 | -0.07  (-0.82, 0.69) | >0.99 |
| Male | 0.82  (0.06, 1.58) | 0.12 | 0.35  (-0.39, 1.10) | 0.75 | 0.73  (-0.03, 1.49) | 0.17 | 0.37  (-0.38, 1.11) | 0.76 |

**Figure S15. Walking and running time comparisons between geriatric crickets and treatment groups. (A)** Overall and **(B)** sex-stratified analyses are shown for geriatric crickets relative to control, acarbose-, rapamycin-, and phenylbutyrate-treated cohorts. The two-way ANOVA results are displayed above the table, summarizing the effects of treatment, sex, and their interaction on walking and running time. Pairwise comparisons were computed using Cohen’s *d* with Hedges’ *g* correction to account for small sample size bias, with effect sizes reported together with 95% CIs in the format (lower, upper). Geriatric crickets were used as the reference group for all comparisons, and adjusted *P*-values were obtained via Dunnett’s multiple comparisons post-hoc test (**P* < 0.05).

| A. B. 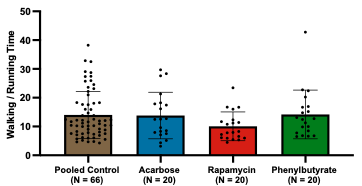 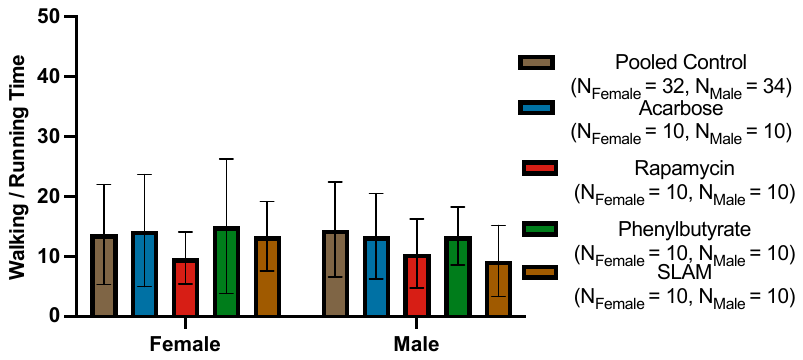 **Source** \| **SS** \| **df** \| **MS** \| **F** \| ***P***; Interaction \| 31.32 \| 3 \| 10.44 \| 0.17 \| 0.92; Sex \| 1.706 \| 1 \| 1.706 \| 0.03 \| 0.87; Group \| 265.9 \| 3 \| 88.62 \| 1.44 \| 0.24; Residual \| 7275 \| 118 \| 61.66 \| \|; Total \| 7574 \| 125 \| \| \| | | | | | | |
| --- | --- | --- | --- | --- | --- | --- |
|  | Pooled Control-Acarbose | | Pooled Control-Rapamycin | | Pooled Control-Phenylbutyrate | |
|  | *d*  (95% CI) | Adj. *P*-Value | *d*  (95% CI) | Adj. *P*-Value | *d*  (95% CI) | Adj. *P*-Value |
| Overall | 0.03 (-0.48, 0.53) | >0.99 | 0.52 (0.01, 1.03) | 0.14 | -0.02 (-0.52, 0.48) | >0.99 |
| Female | -0.07 (-0.78, 0.64) | 0.99 | 0.04 (-0.67, 0.75) | 0.41 | -0.15 (-0.86, 0.56) | 0.94 |
| Male | 0.14 (-0.56, 0.85) | 0.97 | 0.68 (-0.04, 1.40) | 0.39 | 0.14 (-0.56, 0.85) | 0.97 |

**Figure S16. Walking and running time comparisons between pooled control crickets and treatment groups. (A)** Overall and **(B)** sex-stratified analyses present pooled controls in comparison with acarbose-, rapamycin-, and phenylbutyrate-treated cohorts. Results from the two-way ANOVA are shown above the table, outlining the contributions of treatment group, sex, and their interaction to walking and running time. Pairwise comparisons were determined using Cohen’s *d* with Hedges’ *g* correction to address small sample size bias, with effect sizes reported alongside 95% CIs in the format (lower, upper). Pooled controls served as the reference group for all comparisons, and adjusted *P*-values were calculated using Dunnett’s multiple comparisons post-hoc test.

| **Average Walking Speed (cm/s)** | | | | | | | | | |
| --- | --- | --- | --- | --- | --- | --- | --- | --- | --- |
|  | Overall | | | Female | | | Male | | |
|  | Mean (SD) | *d*  (95% CI) | *P* | Mean (SD) | *d*  (95% CI) | *P* | Mean (SD) | *d*  (95% CI) | *P* |
| Control  (N_Female_ = 9, N_Male_ = 10) | 2.83 (1.04) |  |  | 2.43 (0.89) |  |  | 3.20 (1.07) |  |  |
| Acarbose  (N_Female_ = 10, N_Male_ = 10) | 2.40 (0.82) | 0.45  (-0.18, 1.09) | 0.35 | 2.23 (0.77) | 0.23  (−0.67, 1.13) | 0.93 | 2.57 (0.86) | 0.62  (−0.28, 1.52) | 0.31 |
| Rapamycin  (N_Female_ = 10, N_Male_ = 10) | 3.00 (1.26) | -0.14  (-0.77, 0.48) | 0.90 | 2.56 (0.66) | -0.16  (-1.06, 0.74) | 0.98 | 3.44 (1.58) | -0.17  (-1.05, 0.71) | 0.88 |
| Phenylbutyrate  (N_Female_ = 10, N_Male_ = 10) | 2.09 (0.50) | 0.90  (0.24, 1.55) | **0.045** | 2.10 (0.61) | 0.42  (-0.49, 1.33) | 0.77 | 2.09 (0.40) | 1.32  (0.35, 2.28) | **0.023** |

**Table S5. Pairwise comparisons of average walking speed between control and treatment groups.** Overall and sex-stratified analyses are provided for average walking speed in control, acarbose-, rapamycin-, and phenylbutyrate-treated crickets. Effect sizes were calculated using Cohen’s *d* with Hedges’ *g* correction to minimize bias from small sample sizes, with values presented alongside 95% CIs in the format (lower, upper). Controls were used as the reference group for all comparisons, and adjusted *P*-values were obtained through Dunnett’s multiple comparisons post-hoc test.

|  | Mean (SD) | | *d* (95% CI) | *P* |
| --- | --- | --- | --- | --- |
|  | Female | Male |  |  |
| Control | 2.43 (0.89) | 3.20 (1.07) | -0.74 (-1.67, 0.19) | 0.29 |
| Acarbose | 2.23 (0.77) | 2.57 (0.86) | -0.40 (-1.28, 0.49) | >0.99 |
| Rapamycin | 2.56 (0.66) | 3.44 (1.58) | -0.70 (-1.60, 0.21) | 0.14 |
| Phenylbutyrate | 2.10 (0.61) | 2.09 (0.40) | 0.02 (-0.86, 0.90) | >0.99 |

**Table S6. Group means and effect sizes for average walking speed comparing sexes within each treatment group.** Mean values with standard deviations (SD) are reported for average walking speed in control, acarbose-, rapamycin-, and phenylbutyrate-treated crickets. Pairwise comparisons are quantified using Cohen’s *d* with Hedges’ *g* correction to account for small sample size bias. Effect sizes are reported alongside 95% CIs in the format (lower, upper). Females served as the reference group for all comparisons. Adjusted *P*-values were calculated using Bonferroni’s multiple comparisons post-hoc test.

| A. B. 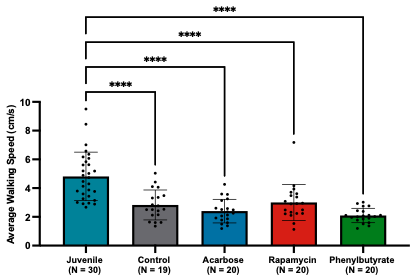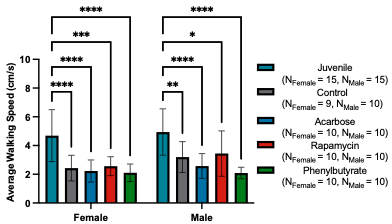 **Source** \| **SS** \| **df** \| **MS** \| **F** \| ***P***; Interaction \| 2.80 \| 4 \| 0.70 \| 0.49 \| 0.74; Sex \| 5.29 \| 1 \| 5.29 \| 3.68 \| 0.06; Group \| 119.1 \| 4 \| 29.77 \| 20.74 \| **<0.0001**; Residual \| 142.1 \| 99 \| 1.44 \| \|; Total \| 269.29 \| 108 \| \| \| | | | | | | | | |
| --- | --- | --- | --- | --- | --- | --- | --- | --- |
|  | Juvenile-Control | | Juvenile-Acarbose | | Juvenile-Rapamycin | | Juvenile-Phenylbutyrate | |
|  | *d*  (95% CI) | Adj. *P*-Value | *d*  (95% CI) | Adj. *P*-Value | *d*  (95% CI) | Adj. *P*-Value | *d*  (95% CI) | Adj. *P*-Value |
| Overall | 1.32  (0.69, 1.95) | **<0.0001** | 1.68  (1.03, 2.34) | **<0.0001** | 1.16  (0.55, 1.77) | **<0.0001** | 1.98  (1.30, 2.67) | **<0.0001** |
| Female | 1.42  (0.50, 2.33) | **<0.0001** | 1.59  (0.68, 2.51) | **<0.0001** | 1.40  (0.51, 2.29) | **0.0001** | 1.71  (0.78, 2.64) | **<0.0001** |
| Male | 1.18  (0.32, 2.05) | **0.0022** | 1.68  (0.75, 2.60) | **<0.0001** | 0.91  (0.07, 1.75) | **0.011** | 2.15  (1.15, 3.15) | **<0.0001** |

**Figure S17. Average walking speed comparisons between juvenile crickets and treatment groups. (A)** Overall and **(B)** sex-stratified analyses evaluate juvenile crickets in relation to control, acarbose-, rapamycin-, and phenylbutyrate-treated cohorts. Results from the two-way ANOVA are provided above the table, highlighting the effects of treatment group, sex, and their interaction on average walking speed. Pairwise contrasts were estimated using Cohen’s *d* with Hedges’ *g* correction to limit small sample size bias, with effect sizes presented alongside 95% CIs in the format (lower, upper). Juvenile crickets served as the reference group for all comparisons, and adjusted *P*-values were derived using Dunnett’s multiple comparisons post-hoc test (**P* < 0.05, ***P* < 0.01, ****P* < 0.001, *****P* < 0.0001).

| A. B. 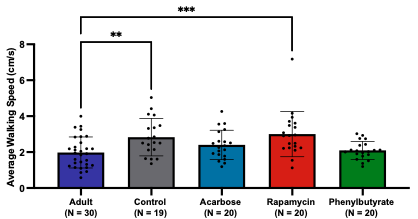 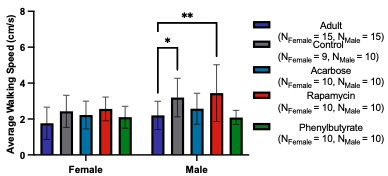 **Source** \| **SS** \| **df** \| **MS** \| **F** \| ***P***; Interaction \| 2.51 \| 4 \| 0.6268 \| 0.78 \| 0.54; Sex \| 6.20 \| 1 \| 6.20 \| 7.68 \| **0.0067**; Group \| 17.66 \| 4 \| 4.41 \| 5.47 \| **0.0005**; Residual \| 79.96 \| 99 \| 0.81 \| \|; Total \| 106.33 \| 108 \| \| \| | | | | | | | | |
| --- | --- | --- | --- | --- | --- | --- | --- | --- |
|  | Adult-Control | | Adult-Acarbose | | Adult-Rapamycin | | Adult-Phenylbutyrate | |
|  | *d*  (95% CI) | Adj. *P*-Value | *d*  (95% CI) | Adj. *P*-Value | *d*  (95% CI) | Adj. *P*-Value | *d*  (95% CI) | Adj. *P*-Value |
| Overall | -0.89  (-1.49, -0.29) | **0.0082** | -0.49  (-1.06, 0.09) | 0.35 | -0.96  (-1.56, -0.37) | **0.0008** | -0.15  (-0.71, 0.42) | 0.98 |
| Female | -0.72  (-1.57, 0.13) | 0.25 | -0.53  (-1.35, 0.28) | 0.55 | -0.95  (-1.79, -0.11) | 0.11 | -0.41  (-1.22, 0.40) | 0.79 |
| Male | -1.06  (-1.92, -0.21) | **0.029** | -0.44  (-1.25, 0.37) | 0.72 | -1.03  (-1.88, -0.18) | **0.0039** | 0.16  (-0.64, 0.96) | 0.99 |

**Figure S18. Average walking speed comparisons between adult crickets and treatment groups. (A)** Overall and **(B)** sex-stratified analyses compare adult crickets with control, acarbose-, rapamycin-, and phenylbutyrate-treated cohorts. Results from the two-way ANOVA are displayed above the table, summarizing the effects of treatment, sex, and their interaction on average walking speed. Pairwise comparisons were calculated using Cohen’s *d* with Hedges’ *g* correction to control for small sample size bias, with effect sizes reported alongside 95% CIs in the format (lower, upper). Adults served as the reference group for all comparisons, and adjusted *P*-values were obtained through Dunnett’s multiple comparisons post-hoc test (**P* < 0.05, ***P* < 0.01, ****P* < 0.001).

| A. B. 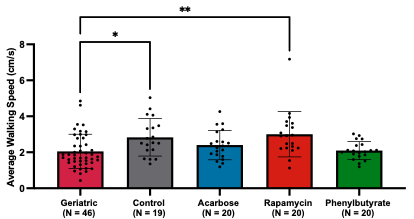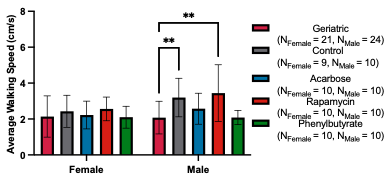 **Source** \| **SS** \| **df** \| **MS** \| **F** \| ***P***; Interaction \| 4.67 \| 4 \| 1.17 \| 1.26 \| 0.29; Sex \| 4.09 \| 1 \| 4.09 \| 4.42 \| **0.038**; Group \| 16.16 \| 4 \| 4.04 \| 4.36 \| **0.003**; Residual \| 105.6 \| 114 \| 0.93 \| \|; Total \| 130.5 \| 123 \| \| \| | | | | | | | | |
| --- | --- | --- | --- | --- | --- | --- | --- | --- |
|  | Geriatric-Control | | Geriatric -Acarbose | | Geriatric-Rapamycin | | Geriatric-Phenylbutyrate | |
|  | *d*  (95% CI) | Adj. *P*-Value | *d*  (95% CI) | Adj. *P*-Value | *d*  (95% CI) | Adj. *P*-Value | *d*  (95% CI) | Adj. *P*-Value |
| Overall | -0.78  (-1.34, -0.23) | **0.012** | -0.38  (-0.91, 0.15) | 0.50 | -0.89  (-1.43, -0.34) | **0.0011** | -0.05  (-0.57, 0.48) | >0.99 |
| Female | -0.26  (-1.04, 0.52) | 0.89 | -0.08  (-0.84, 0.67) | >0.99 | -0.40  (-1.16, 0.36) | 0.65 | 0.04  (-0.71, 0.79) | >0.99 |
| Male | -1.14  (-1.93, -0.36) | **0.01** | -0.53  (-1.28, 0.21) | 0.50 | -1.17  (-1.95, -0.38) | **0.0011** | -0.01  (-0.75, 0.73) | >0.99 |

**Figure S19. Average walking speed comparisons between historical geriatric crickets and treatment groups. (A)** Overall and **(B)** sex-stratified analyses present geriatric crickets relative to control, acarbose-, rapamycin-, and phenylbutyrate-treated cohorts. The results from the two-way ANOVA are shown above the table, describing the effects of treatment group, sex, and their interaction on average walking speed. Pairwise comparisons were determined using Cohen’s *d* with Hedges’ *g* correction to account for small sample size bias, with effect sizes provided alongside 95% CIs in the format (lower, upper). Geriatric crickets were used as the reference group for all comparisons, and adjusted *P*-values were calculated using Dunnett’s multiple comparisons post-hoc test (**P* < 0.05, ***P* < 0.01).

| A. B. 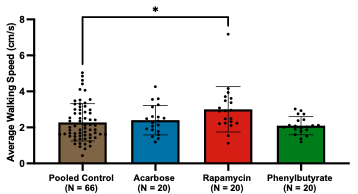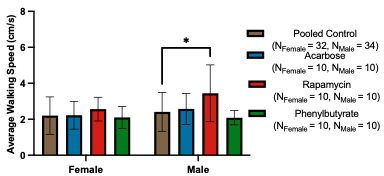 **Source** \| **SS** \| **df** \| **MS** \| **F** \| ***P***; Interaction \| 2.32 \| 3 \| 0.77 \| 0.79 \| 0.50; Sex \| 3.06 \| 1 \| 3.06 \| 3.14 \| 0.079; Group \| 9.75 \| 3 \| 3.25 \| 3.33 \| **0.022**; Residual \| 115.2 \| 118 \| 0.98 \| \|; Total \| 130.3 \| 125 \| \| \| | | | | | | |
| --- | --- | --- | --- | --- | --- | --- |
|  | Pooled Control-Acarbose | | Pooled Control-Rapamycin | | Pooled Control-Phenylbutyrate | |
|  | *d*  (95% CI) | Adj. *P*-Value | *d*  (95% CI) | Adj. *P*-Value | *d*  (95% CI) | Adj. *P*-Value |
| Overall | -0.12 (-0.62, 0.38) | 0.95 | -0.65 (-1.16, -0.14) | **0.014** | 0.20 (-0.30, 0.70) | 0.83 |
| Female | -0.03  (-0.74, 0.68) | >0.99 | -0.17  (-0.88, 0.54) | 0.66 | 0.10  (-0.61, 0.81) | 0.99 |
| Male | -0.15  (-0.86, 0.55) | 0.95 | -0.84  (-1.57, -0.12) | **0.013** | 0.32  (-0.39, 1.03) | 0.73 |

**Figure S20. Average walking speed comparisons between pooled control crickets and treatment groups. (A)** Overall and **(B)** sex-stratified analyses compare pooled controls with acarbose-, rapamycin-, and phenylbutyrate-treated cohorts. Results from the two-way ANOVA are presented above the table, outlining the effects of treatment, sex, and their interaction on average walking speed. Pairwise differences were computed using Cohen’s *d* with Hedges’ *g* correction to minimize small sample size bias, with effect sizes reported together with 95% CIs in the format (lower, upper). Pooled controls served as the reference group for all comparisons, and adjusted *P*-values were obtained using Dunnett’s multiple comparisons post-hoc test (**P* < 0.05).

| **Average Running Speed (cm/s)** | | | | | | | | | |
| --- | --- | --- | --- | --- | --- | --- | --- | --- | --- |
|  | Overall | | | Female | | | Male | | |
|  | Mean (SD) | *d*  (95% CI) | *P* | Mean (SD) | *d*  (95% CI) | *P* | Mean (SD) | *d*  (95% CI) | *P* |
| Control  (N_Female_ = 9, N_Male_ = 10) | 27.9 (5.9) |  |  | 27.3 (7.65) |  |  | 28.4 (4.12) |  |  |
| Acarbose  (N_Female_ = 10, N_Male_ = 10) | 22.7 (5.2) | 0.92  (0.26, 1.58) | **0.02** | 21.18 (5.20) | 0.90  (−0.04, 1.85) | 0.06 | 24.29 (5.07) | 0.85  (−0.06, 1.77) | 0.27 |
| Rapamycin  (N_Female_ = 10, N_Male_ = 10) | 29.2 (6.7) | -0.20  (-0.83, 0.43) | 0.81 | 27.1 (6.61) | 0.03  (-0.87, 0.93) | >0.99 | 31.4 (6.41) | -0.53  (-1.43, 0.36) | 0.52 |
| Phenylbutyrate  (N_Female_ = 10, N_Male_ = 10) | 23.1 (5.3) | 0.84  (0.18, 1.49) | 0.033 | 24.1 (5.30) | 0.47  (-0.44, 1.38) | 0.48 | 22.1 (5.45) | 1.25  (0.29, 2.21) | **0.045** |

**Table S7. Pairwise comparisons of average running speed between control and treatment groups.** Overall and sex-stratified analyses are shown for average running speed in control, acarbose-, rapamycin-, and phenylbutyrate-treated crickets. Effect sizes for each comparison were determined using Cohen’s *d* with Hedges’ *g* correction to account for small sample size bias, with values reported together with 95% CIs in the format (lower, upper). Controls served as the reference group for all comparisons, and adjusted *P*-values were calculated using Dunnett’s multiple comparisons post-hoc test.

| A. B. 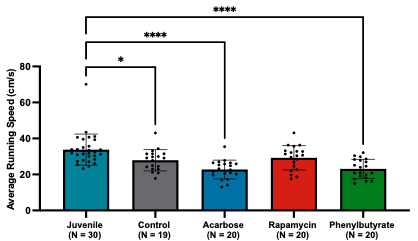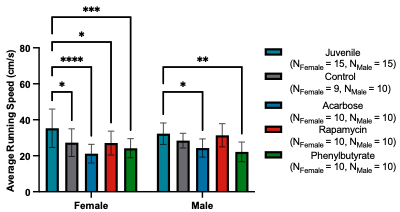 **Source** \| **SS** \| **df** \| **MS** \| **F** \| ***P***; Interaction \| 232.3 \| 4 \| 58.07 \| 1.29 \| 0.28; Sex \| 12.85 \| 1 \| 12.85 \| 0.28 \| 0.59; Group \| 2063 \| 4 \| 515.7 \| 11.42 \| **<0.0001**; Residual \| 4473 \| 99 \| 45.18 \| \|; Total \| 6781 \| 108 \| \| \| | | | | | | | | |
| --- | --- | --- | --- | --- | --- | --- | --- | --- |
|  | Juvenile-Control | | Juvenile-Acarbose | | Juvenile-Rapamycin | | Juvenile-Phenylbutyrate | |
|  | *d*  (95% CI) | Adj. *P*-Value | *d*  (95% CI) | Adj. *P*-Value | *d*  (95% CI) | Adj. *P*-Value | *d*  (95% CI) | Adj. *P*-Value |
| Overall | 0.75  (0.16, 1.34) | **0.013** | 1.46  (0.83, 2.09) | **<0.0001** | 0.57  (-0.01, 1.14) | 0.075 | 1.40  (0.77, 2.03) | **<0.0001** |
| Female | 0.80  (-0.06, 1.65) | **0.02** | 1.53  (0.62, 2.43) | **<0.0001** | 0.85  (0.02, 1.69) | **0.013** | 1.21  (0.34, 2.07) | **0.0004** |
| Male | 0.70  (-0.12, 1.52) | 0.46 | 1.37  (0.48, 2.25) | **0.017** | 0.14  (-0.66, 0.94) | 0.99 | 1.70  (0.77, 2.63) | **0.0013** |

**Figure S21. Average running speed comparisons between juvenile crickets and treatment groups. (A)** Overall and **(B)** sex-stratified analyses show juvenile crickets evaluated against control, acarbose-, rapamycin-, and phenylbutyrate-treated cohorts. The two-way ANOVA results are provided above the table, summarizing the roles of treatment, sex, and their interaction in influencing average running speed. Pairwise comparisons were derived using Cohen’s *d* with Hedges’ *g* correction to reduce bias from small sample sizes, with effect sizes displayed alongside 95% CIs in the format (lower, upper). Juvenile crickets were designated as the reference group for all comparisons, and adjusted *P*-values were calculated using Dunnett’s multiple comparisons post-hoc test (**P* < 0.05, ***P* < 0.01, ****P* < 0.001, *****P* < 0.0001).

| A. B. 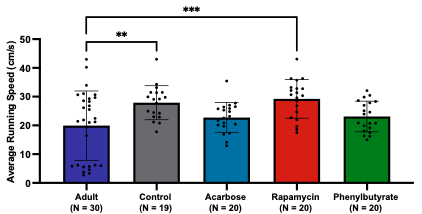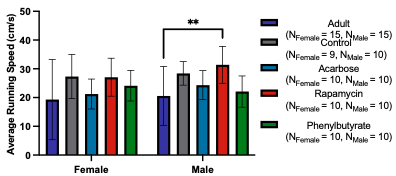 **Source** \| **SS** \| **df** \| **MS** \| **F** \| ***P***; Interaction \| 115.9 \| 4 \| 28.99 \| 0.44 \| 0.78; Sex \| 63.51 \| 1 \| 63.51 \| 0.96 \| 0.33; Group \| 1377 \| 4 \| 344.3 \| 5.18 \| **0.0008**; Residual \| 6584 \| 99 \| 66.50 \| \|; Total \| 8140 \| 108 \| \| \| | | | | | | | | |
| --- | --- | --- | --- | --- | --- | --- | --- | --- |
|  | Adult-Control | | Adult-Acarbose | | Adult-Rapamycin | | Adult-Phenylbutyrate | |
|  | *d*  (95% CI) | Adj. *P*-Value | *d*  (95% CI) | Adj. *P*-Value | *d*  (95% CI) | Adj. *P*-Value | *d*  (95% CI) | Adj. *P*-Value |
| Overall | -0.77  (-1.37, -0.18) | **0.0040** | -0.28  (-0.84, 0.29) | 0.59 | -0.89  (-1.48, -0.30) | **0.0005** | -0.31  (-0.88, 0.25) | 0.48 |
| Female | -0.64  (-1.49, 0.20) | 0.08 | -0.16  (-0.96, 0.64) | 0.95 | -0.65  (-1.47, 0.17) | 0.08 | -0.41  (-1.22, 0.40) | 0.43 |
| Male | -0.90  (-1.74, -0.06) | 0.07 | -0.42  (-1.23, 0.39) | 0.65 | -1.17  (-2.03, -0.31) | **0.006** | -0.17  (-0.97, 0.63) | 0.98 |

**Figure S22. Average running speed comparisons between adult crickets and treatment groups. (A)** Overall and **(B)** sex-stratified analyses compare adult crickets to control, acarbose-, rapamycin-, and phenylbutyrate-treated cohorts. Results from the two-way ANOVA are shown above the table, detailing the effects of treatment group, sex, and their interaction on average running speed. Pairwise comparisons were calculated using Cohen’s *d* with Hedges’ *g* correction to account for small sample size bias, with effect sizes reported alongside 95% CIs in the format (lower, upper). Adult crickets served as the reference group for all comparisons, and adjusted *P*-values were determined using Dunnett’s multiple comparisons post-hoc test (***P* < 0.01, ****P* < 0.001).

| A. B. 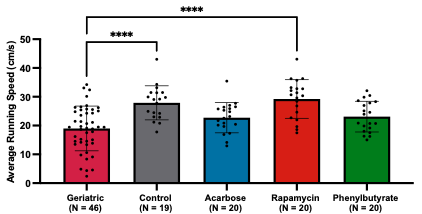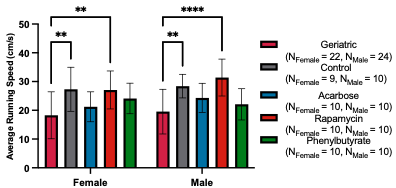 **Source** \| **SS** \| **df** \| **MS** \| **F** \| ***P***; Interaction \| 116.0 \| 4 \| 29.00 \| 0.64 \| 0.63; Sex \| 67.42 \| 1 \| 67.42 \| 1.50 \| 0.22; Group \| 1991 \| 4 \| 497.8 \| 11.07 \| **<0.0001**; Residual \| 5173 \| 115 \| 44.99 \| \|; Total \| 7347 \| 124 \| \| \| | | | | | | | | |
| --- | --- | --- | --- | --- | --- | --- | --- | --- |
|  | Geriatric-Control | | Geriatric -Acarbose | | Geriatric-Rapamycin | | Geriatric-Phenylbutyrate | |
|  | *d*  (95% CI) | Adj. *P*-Value | *d*  (95% CI) | Adj. *P*-Value | *d*  (95% CI) | Adj. *P*-Value | *d*  (95% CI) | Adj. *P*-Value |
| Overall | -1.21  (-1.79, -0.63) | **<0.0001** | -0.52  (-1.05, 0.02) | 0.13 | -1.35  (-1.93, -0.78) | **<0.0001** | -0.57  (-1.11, -0.03) | 0.08 |
| Female | -1.10  (-1.92, -0.27) | **0.0035** | -0.39  (-1.14, 0.37) | 0.66 | -1.12  (-1.92, -0.31) | **0.0031** | -0.77  (-1.55, 0.01) | 0.09 |
| Male | -1.24  (-2.04, -0.45) | **0.0025** | -0.65  (-1.40, 0.10) | 0.21 | -1.56  (-2.38, -0.73) | **<0.0001** | -0.35  (-1.09, 0.40) | 0.75 |

**Figure S23. Average running speed comparisons between geriatric crickets and treatment groups. (A)** Overall and **(B)** sex-stratified analyses present geriatric crickets in comparison with control, acarbose-, rapamycin-, and phenylbutyrate-treated cohorts. The results from the two-way ANOVA are provided above the table, summarizing the contributions of treatment group, sex, and their interaction to average running speed. Pairwise comparisons were computed using Cohen’s *d* with Hedges’ *g* correction to adjust for small sample size bias, with effect sizes shown alongside 95% CIs in the format (lower, upper). Geriatric crickets served as the reference group for all comparisons, and adjusted *P*-values were obtained using Dunnett’s multiple comparisons post-hoc test (***P* < 0.01, *****P* < 0.0001).

| A. B. 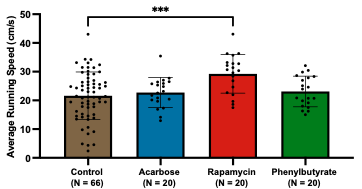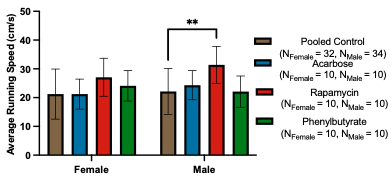 **Source** \| **SS** \| **df** \| **MS** \| **F** \| ***P***; Interaction \| 119.5 \| 3 \| 39.82 \| 0.76 \| 0.52; Sex \| 60.36 \| 1 \| 60.36 \| 1.15 \| 0.29; Group \| 880.1 \| 3 \| 293.4 \| 5.57 \| 0.0013; Residual \| 6211 \| 118 \| 52.63 \| \|; Total \| 7271 \| 125 \| \| \| | | | | | | |
| --- | --- | --- | --- | --- | --- | --- |
|  | Pooled Control-Acarbose | | Pooled Control-Rapamycin | | Pooled Control-Phenylbutyrate | |
|  | *d*  (95% CI) | Adj. *P*-Value | *d*  (95% CI) | Adj. *P*-Value | *d*  (95% CI) | Adj. *P*-Value |
| Overall | -0.15 (-0.65, 0.35) | 0.89 | -0.95 (-1.47, -0.43) | **0.0002** | -0.19 (-0.70, 0.31) | 0.79 |
| Female | 0.01 (-0.70, 0.72) | >0.99 | -0.69  (-1.42, 0.03) | 0.08 | -0.35 (-1.06, 0.36) | 0.60 |
| Male | -0.28 (-0.99, 0.43) | 0.79 | -1.18  (-1.93, -0.43) | **0.0017** | 0.01 (-0.70, 0.71) | >0.99 |

**Figure S24. Average running speed comparisons between pooled control crickets and treatment groups. (A)** Overall and **(B)** sex-stratified analyses show pooled controls evaluated against acarbose-, rapamycin-, and phenylbutyrate-treated cohorts. Results from the two-way ANOVA are presented above the table, outlining the effects of treatment group, sex, and their interaction on average running speed. Pairwise comparisons were calculated using Cohen’s *d* with Hedges’ *g* correction to minimize bias associated with small sample sizes, with effect sizes reported alongside 95% CIs in the format (lower, upper). Pooled controls served as the reference group for all comparisons, and adjusted *P*-values were derived using Dunnett’s multiple comparisons post-hoc test (***P* < 0.01, ****P* < 0.001).

**Appendix 4. OFT Measures of Exploration.**

| **Central/Peripheral Distance** | | | | | | | | | |
| --- | --- | --- | --- | --- | --- | --- | --- | --- | --- |
|  | Overall | | | Female | | | Male | | |
|  | Mean (SD) | *d*  (95% CI) | *P* | Mean (SD) | *d*  (95% CI) | *P* | Mean (SD) | *d*  (95% CI) | *P* |
| Control  (N_Female_ = 9, N_Male_ = 10) | 0.51 (0.25) |  |  | 0.47 (0.21) |  |  | 0.55 (0.28) |  |  |
| Acarbose  (N_Female_ = 10, N_Male_ = 10) | 0.26 (0.40) | 0.73  (0.08, 1.38) | 0.28 | 0.14 (0.13) | 1.83  (0.76, 2.90) | 0.30 | 0.38 (0.54) | 0.38  (−0.51, 1.26) | 0.79 |
| Rapamycin  (N_Female_ = 10, N_Male_ = 10) | 0.79 (0.64) | -0.56  (-1.20, 0.08) | 0.20 | 0.62 (0.49) | -0.37  (-1.28, 0.54) | 0.84 | 0.97 (0.74) | -0.72  (-1.62, 0.19) | 0.14 |
| Phenylbutyrate  (N_Female_ = 10, N_Male_ = 10) | 0.58 (0.62) | -0.14  (-0.77, 0.49) | 0.94 | 0.86 (0.77) | -0.64  (-1.57, 0.28) | 0.20 | 0.31 (0.20) | 0.94  (0.02, 1.87) | 0.57 |

**Table S1. Pairwise comparisons of central-to-peripheral distance ratio between control and treatment groups.** Overall and sex-stratified analyses are presented for the central/peripheral distance ratio in control, acarbose-, rapamycin-, and phenylbutyrate-treated crickets. Effect sizes were computed using Cohen’s *d* with Hedges’ *g* correction to reduce small sample size bias, with values reported alongside 95% confidence intervals (CIs) in the format (lower, upper). Controls were used as the reference group for all comparisons, and adjusted *P*-values were derived using Dunnett’s multiple comparisons post-hoc test.

|  | Mean (SD) | |  |  |
| --- | --- | --- | --- | --- |
|  | Female | Male | *d* (95% CI) | *P* |
| Control | 0.47 (0.21) | 0.55 (0.28) | -0.31 (-1.21, 0.60) | >0.99 |
| Acarbose | 0.14 (0.13) | 0.38 (0.54) | -0.59 (-1.48, 0.31) | >0.99 |
| Rapamycin | 0.62 (0.49) | 0.97 (0.74) | -0.53 (-1.43, 0.36) | 0.44 |
| Phenylbutyrate | 0.86 (0.77) | 0.31 (0.20) | 0.94 (-0.09, 1.86) | 0.055 |

**Table S2. Group means and effect sizes for central to peripheral distance ratios comparing sexes within each treatment group.** Mean values with standard deviations (SD) are reported for each locomotor and exploratory measure in control, acarbose-, rapamycin-, and phenylbutyrate-treated crickets. Pairwise comparisons are quantified using Cohen’s *d* with Hedges’ *g* correction to account for small sample size bias. Effect sizes are reported alongside 95% CIs in the format (lower, upper). Females served as the reference group for all comparisons. Adjusted *P*-values were calculated using Bonferroni’s multiple comparisons post-hoc test.

| A. B. 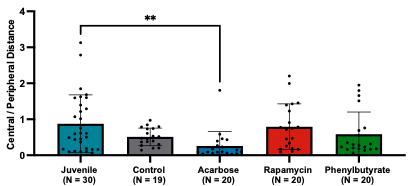 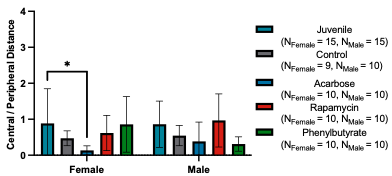 **Source** \| **SS** \| **df** \| **MS** \| **F** \| ***P***; Interaction \| 2.44 \| 4 \| 0.61 \| 1.71 \| 0.15; Sex \| 0.01 \| 1 \| 0.01 \| 0.03 \| 0.86; Group \| 5.37 \| 4 \| 1.34 \| 3.76 \| **0.0068**; Residual \| 35.34 \| 99 \| 0.36 \| \|; Total \| 43.16 \| 108 \| \| \| | | | | | | | | |
| --- | --- | --- | --- | --- | --- | --- | --- | --- |
|  | Juvenile-Control | | Juvenile-Acarbose | | Juvenile-Rapamycin | | Juvenile-Phenylbutyrate | |
|  | *d*  (95% CI) | Adj. *P*-Value | *d*  (95% CI) | Adj. *P*-Value | *d*  (95% CI) | Adj. *P*-Value | *d*  (95% CI) | Adj. *P*-Value |
| Overall | 0.55  (-0.04, 1.13) | 0.14 | 0.90  (0.30, 1.49) | **0.0024** | 0.11  (-0.46, 0.67) | 0.98 | 0.39  (-0.18, 0.96) | 0.30 |
| Female | 0.52  (-0.32, 1.36) | 0.31 | 0.96  (0.12, 1.81) | **0.010** | 0.32  (-0.48, 1.13) | 0.67 | 0.03  (-0.77, 0.83) | >0.99 |
| Male | 0.57  (-0.25, 1.38) | 0.53 | 0.77  (-0.06, 1.60) | 0.17 | -0.16  (-0.96, 0.65) | 0.98 | 1.03  (0.18, 1.88) | 0.091 |

**Figure S1. Central-to-peripheral distance ratio comparisons between juvenile crickets and treatment groups. (A)** Overall and **(B)** sex-stratified analyses present juvenile crickets relative to control, acarbose-, rapamycin-, and phenylbutyrate-treated cohorts. Results from the two-way ANOVA are shown above the table, summarizing the effects of treatment, sex, and their interaction on the central/peripheral distance ratio. Pairwise comparisons were quantified using Cohen’s *d* with Hedges’ *g* correction to account for small sample size bias, with effect sizes provided alongside 95% CIs in the format (lower, upper). Juvenile crickets were designated as the reference group for all comparisons, and adjusted *P*-values were obtained using Dunnett’s multiple comparisons post-hoc test (**P* < 0.05, ***P* < 0.01).

| A. B. 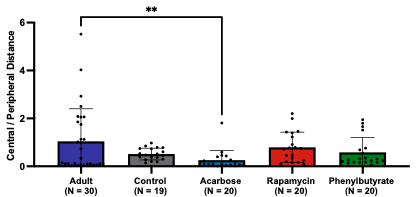 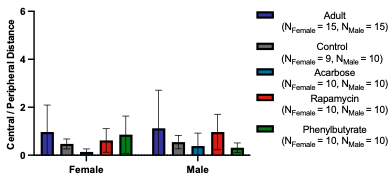 **Source** \| **SS** \| **df** \| **MS** \| **F** \| ***P***; Interaction \| 2.51 \| 4 \| 0.63 \| 0.89 \| 0.47; Sex \| 0.08 \| 1 \| 0.08 \| 0.12 \| 0.73; Group \| 8.49 \| 4 \| 2.12 \| 3.01 \| **0.022**; Residual \| 69.84 \| 99 \| 0.71 \| \|; Total \| 80.92 \| 108 \| \| \| | | | | | | | | |
| --- | --- | --- | --- | --- | --- | --- | --- | --- |
|  | Adult-Control | | Adult-Acarbose | | Adult-Rapamycin | | Adult-Phenylbutyrate | |
|  | *d*  (95% CI) | Adj. *P*-Value | *d*  (95% CI) | Adj. *P*-Value | *d*  (95% CI) | Adj. *P*-Value | *d*  (95% CI) | Adj. *P*-Value |
| Overall | 0.48  (-0.10, 1.07) | 0.11 | 0.71  (0.12, 1.29) | **0.0059** | 0.22  (-0.35, 0.78) | 0.71 | 0.40  (-0.17, 0.97) | 0.19 |
| Female | 0.53  (-0.31, 1.37) | 0.46 | 0.91  (0.07, 1.75) | 0.062 | 0.36  (-0.44, 1.17) | 0.73 | 0.11  (-0.69, 0.91) | 0.99 |
| Male | 0.44  (-0.37, 1.25) | 0.30 | 0.56  (-0.26, 1.37) | 0.12 | 0.11  (-0.69, 0.91) | 0.98 | 0.63  (-0.19, 1.45) | 0.071 |

**Figure S2. Central-to-peripheral distance ratio comparisons between adult crickets and treatment groups. (A)** Overall and **(B)** sex-stratified analyses compare adult crickets to control, acarbose-, rapamycin-, and phenylbutyrate-treated cohorts. Results from the two-way ANOVA are presented above the table, detailing the effects of treatment group, sex, and their interaction on the central/peripheral distance ratio. Pairwise comparisons were calculated using Cohen’s *d* with Hedges’ *g* correction to adjust for small sample size bias, with effect sizes reported alongside 95% CIs in the format (lower, upper). Adult crickets served as the reference group for all comparisons, and adjusted *P*-values were derived using Dunnett’s multiple comparisons post-hoc test (***P* < 0.01).

| A. B. 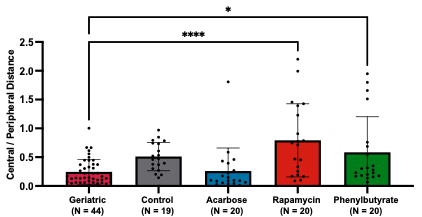 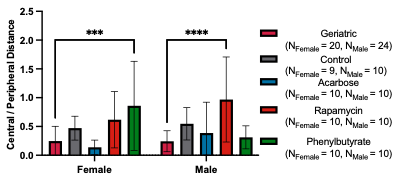 **Source** \| **SS** \| **df** \| **MS** \| **F** \| ***P***; Interaction \| 2.43 \| 4 \| 0.61 \| 3.69 \| **0.0073**; Sex \| 0.02 \| 1 \| 0.02 \| 0.10 \| 0.75; Group \| 5.31 \| 4 \| 1.33 \| 8.05 \| **<0.0001**; Residual \| 18.61 \| 113 \| 0.16 \| \|; Total \| 26.37 \| 122 \| \| \| | | | | | | | | |
| --- | --- | --- | --- | --- | --- | --- | --- | --- |
|  | Geriatric-Control | | Geriatric -Acarbose | | Geriatric-Rapamycin | | Geriatric-Phenylbutyrate | |
|  | *d*  (95% CI) | Adj. *P*-Value | *d*  (95% CI) | Adj. *P*-Value | *d*  (95% CI) | Adj. *P*-Value | *d*  (95% CI) | Adj. *P*-Value |
| Overall | -1.16  (-1.74, -0.59) | 0.085 | -0.07  (-0.60, 0.46) | >0.99 | -1.37  (-1.95, -0.79) | **<0.0001** | -0.87  (-1.42, -0.32) | **0.013** |
| Female | -0.87  (-1.68, -0.06) | 0.49 | 0.47  (-0.29, 1.23) | 0.91 | -1.04  (-1.83, -0.24) | 0.072 | -1.24  (-2.05, -0.42) | **0.0007** |
| Male | -1.42  (-2.23, -0.61) | 0.17 | -0.42  (-1.17, 0.32) | 0.81 | -1.69  (-2.53, -0.85) | **<0.0001** | -0.37  (-1.11, 0.38) | 0.98 |

**Figure S3. Central-to-peripheral distance ratio comparisons between historical geriatric crickets and treatment groups. (A)** Overall and **(B)** sex-stratified analyses show historical geriatric crickets compared with control, acarbose-, rapamycin-, and phenylbutyrate-treated cohorts. Results from the two-way ANOVA are provided above the table, summarizing the effects of treatment, sex, and their interaction on the central/peripheral distance ratio. Pairwise comparisons were determined using Cohen’s *d* with Hedges’ *g* correction to minimize small sample size bias, with effect sizes presented alongside 95% CIs in the format (lower, upper). Historical geriatric crickets served as the reference group for all comparisons, and adjusted *P*-values were calculated using Dunnett’s multiple comparisons post-hoc test (**P* < 0.05, ****P* < 0.001, *****P* < 0.0001).

| A. B. 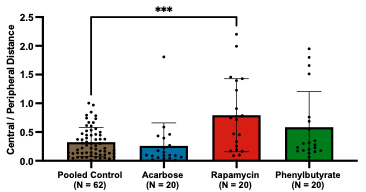 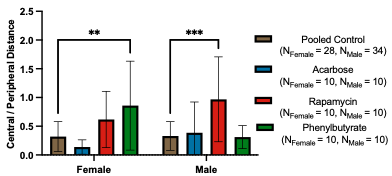 **Source** \| **SS** \| **df** \| **MS** \| **F** \| ***P***; Interaction \| 2.41 \| 3 \| 0.80 \| 4.71 \| **0.0039**; Sex \| 0.01 \| 1 \| 0.01 \| 0.03 \| 0.86; Group \| 4.37 \| 3 \| 1.46 \| 8.54 \| **<0.0001**; Residual \| 19.44 \| 114 \| 0.17 \| \|; Total \| 26.23 \| 121 \| \| \| | | | | | | |
| --- | --- | --- | --- | --- | --- | --- |
|  | Pooled Control-Acarbose | | Pooled Control-Rapamycin | | Pooled Control-Phenylbutyrate | |
|  | *d*  (95% CI) | Adj. *P*-Value | *d*  (95% CI) | Adj. *P*-Value | *d*  (95% CI) | Adj. *P*-Value |
| Overall | 0.22 (-0.29, 0.72) | 0.91 | -1.22 (-1.76, -0.68) | **0.0001** | -0.69 (-1.20, -0.18) | 0.058 |
| Female | 0.75 (0.01, 1.49) | 0.53 | -0.88  (-1.63, -0.13) | 0.14 | -1.19 (-1.95, -0.42) | **0.0017** |
| Male | -0.15 (-0.85, 0.56) | 0.98 | -1.54  (-2.31, -0.76) | **0.0001** | 0.08 (-0.62, 0.79) | >0.99 |

**Figure S4. Central-to-peripheral distance ratio comparisons between pooled control crickets and treatment groups. (A)** Overall and **(B)** sex-stratified analyses present pooled controls in comparison with acarbose-, rapamycin-, and phenylbutyrate-treated cohorts. Results from the two-way ANOVA are displayed above the table, outlining the effects of treatment, sex, and their interaction on the central/peripheral distance ratio. Pairwise comparisons were computed using Cohen’s *d* with Hedges’ *g* correction to account for small sample size bias, with effect sizes shown alongside 95% CIs in the format (lower, upper). Pooled controls were used as the reference group for all comparisons, and adjusted *P*-values were obtained using Dunnett’s multiple comparisons post-hoc test (***P* < 0.01, ****P* < 0.001).

| **Central/Peripheral Time** | | | | | | | | | |
| --- | --- | --- | --- | --- | --- | --- | --- | --- | --- |
|  | Overall | | | Female | | | Male | | |
|  | Mean (SD) | *d*  (95% CI) | *P* | Mean (SD) | *d*  (95% CI) | *P* | Mean (SD) | *d*  (95% CI) | *P* |
| Control  (N_Female_ = 9, N_Male_ = 10) | 0.19 (0.14) |  |  | 0.16 (0.12) |  |  | 0.22 (0.15) |  |  |
| Acarbose  (N_Female_ = 10, N_Male_ = 10) | 0.26 (0.40) | -0.23  (-0.86, 0.40) | 0.81 | 0.06 (0.07) | 0.99  (0.03, 1.94) | 0.63 | 0.14 (0.21) | 0.42  (−0.47, 1.31) | 0.74 |
| Rapamycin  (N_Female_ = 10, N_Male_ = 10) | 0.38 (0.36) | -0.67  (-1.32, -0.03) | 0.12 | 0.27 (0.24) | -0.54  (-1.46, 0.37) | 0.54 | 0.49 (0.43) | -0.80  (-1.71, 0.11) | **0.022** |
| Phenylbutyrate  (N_Female_ = 10, N_Male_ = 10) | 0.18 (0.16) | 0.07  (-0.56, 0.69) | >0.99 | 0.24 (0.20) | -0.46  (-1.37, 0.46) | 0.78 | 0.11 (0.08) | 0.88  (-0.04, 1.79) | 0.57 |

**Table S3. Pairwise comparisons of central-to-peripheral time ratio between control and treatment groups.** Overall and sex-stratified analyses are shown for the central/peripheral time ratio in control, acarbose-, rapamycin-, and phenylbutyrate-treated crickets. Effect sizes for each comparison were calculated using Cohen’s *d* with Hedges’ *g* correction to minimize small sample size bias, with values reported together with 95% CIs in the format (lower, upper). Controls served as the reference group for all comparisons, and adjusted *P*-values were obtained through Dunnett’s multiple comparisons post-hoc test.

| A. B. 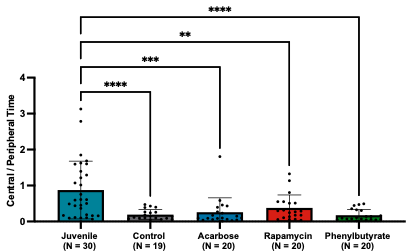 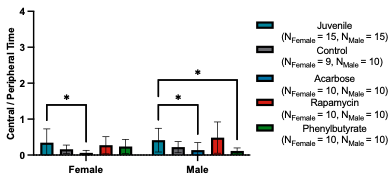 **Source** \| **SS** \| **df** \| **MS** \| **F** \| ***P***; Interaction \| 0.29 \| 4 \| 0.07 \| 1.02 \| 0.40; Sex \| 0.09 \| 1 \| 0.09 \| 1.32 \| 0.25; Group \| 1.46 \| 4 \| 0.36 \| 5.16 \| **0.0008**; Residual \| 6.98 \| 99 \| 0.07 \| \|; Total \| 8.82 \| 108 \| \| \| | | | | | | | | |
| --- | --- | --- | --- | --- | --- | --- | --- | --- |
|  | Juvenile-Control | | Juvenile-Acarbose | | Juvenile-Rapamycin | | Juvenile-Phenylbutyrate | |
|  | *d*  (95% CI) | Adj. *P*-Value | *d*  (95% CI) | Adj. *P*-Value | *d*  (95% CI) | Adj. *P*-Value | *d*  (95% CI) | Adj. *P*-Value |
| Overall | 0.65  (0.06, 1.24) | **<0.0001** | 0.32  (-0.25, 0.89) | **0.0001** | 0.00  (-0.57, 0.57) | **0.0028** | 0.68  (0.10, 1.26) | **<0.0001** |
| Female | 0.56  (-0.28, 1.40) | 0.32 | 0.90  (0.07, 1.74) | **0.039** | 0.20  (-0.60, 1.01) | 0.93 | 0.30  (-0.50, 1.10) | 0.75 |
| Male | 0.71  (-0.12, 1.53) | 0.24 | 0.94  (0.10, 1.78) | **0.044** | -0.18  (-0.98, 0.62) | 0.92 | 1.14  (0.28, 2.00) | **0.025** |

**Figure S5. Central-to-peripheral time ratio comparisons between juvenile crickets and treatment groups. (A)** Overall and **(B)** sex-stratified analyses evaluate juvenile crickets relative to control, acarbose-, rapamycin-, and phenylbutyrate-treated cohorts. Results from the two-way ANOVA are presented above the table, summarizing the effects of treatment group, sex, and their interaction on the central/peripheral time ratio. Pairwise comparisons were determined using Cohen’s *d* with Hedges’ *g* correction to correct for small sample size bias, with effect sizes reported alongside 95% CIs in the format (lower, upper). Juvenile crickets served as the reference group for all comparisons, and adjusted *P*-values were calculated using Dunnett’s multiple comparisons post-hoc test (**P* < 0.05, ***P* < 0.01, ****P* < 0.001, *****P* < 0.0001).

| A. B. 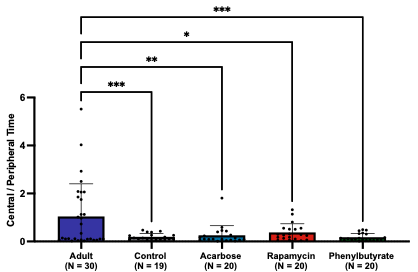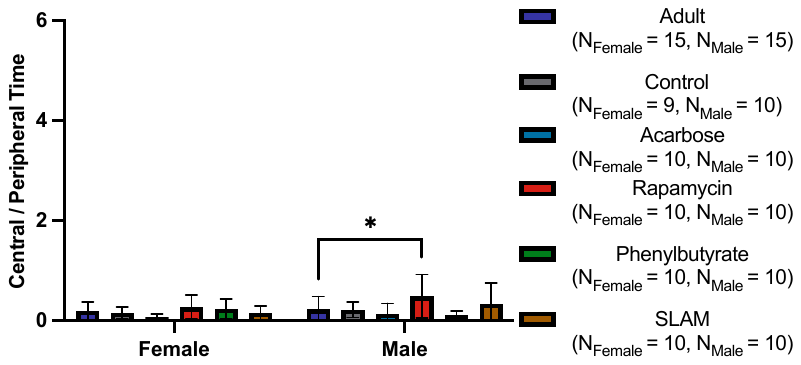 **Source** \| **SS** \| **df** \| **MS** \| **F** \| ***P***; Interaction \| 0.29 \| 4 \| 0.07 \| 1.51 \| 0.20; Sex \| 0.07 \| 1 \| 0.07 \| 1.48 \| 0.23; Group \| 0.85 \| 4 \| 0.21 \| 4.43 \| **0.0024**; Residual \| 4.73 \| 99 \| 0.05 \| \|; Total \| 5.94 \| 108 \| \| \| | | | | | | | | |
| --- | --- | --- | --- | --- | --- | --- | --- | --- |
|  | Adult-Control | | Adult-Acarbose | | Adult-Rapamycin | | Adult-Phenylbutyrate | |
|  | *d*  (95% CI) | Adj. *P*-Value | *d*  (95% CI) | Adj. *P*-Value | *d*  (95% CI) | Adj. *P*-Value | *d*  (95% CI) | Adj. *P*-Value |
| Overall | 0.15  (-0.42, 0.73) | **0.0009** | -0.13  (-0.70, 0.44) | **0.0020** | -0.55  (-1.13, 0.02) | **0.012** | 0.20  (-0.37, 0.77) | **0.0005** |
| Female | 0.24  (-0.59, 1.07) | 0.99 | 0.92  (0.08, 1.76) | 0.38 | -0.33  (-1.13, 0.48) | 0.84 | -0.21  (-1.01, 0.60) | 0.98 |
| Male | 0.04  (-0.76, 0.84) | >0.99 | 0.37  (-0.44, 1.18) | 0.71 | -0.76  (-1.58, 0.07) | **0.019** | 0.58  (-0.24, 1.39) | 0.52 |

**Figure S6. Central-to-peripheral time ratio comparisons between adult crickets and treatment groups. (A)** Overall and **(B)** sex-stratified analyses compare adult crickets to control, acarbose-, rapamycin-, and phenylbutyrate-treated cohorts. Results from the two-way ANOVA are shown above the table, detailing the effects of treatment, sex, and their interaction on the central/peripheral time ratio. Pairwise comparisons were calculated using Cohen’s *d* with Hedges’ *g* correction to address small sample size bias, with effect sizes presented alongside 95% CIs in the format (lower, upper). Adult crickets served as the reference group for all comparisons, and adjusted *P*-values were obtained using Dunnett’s multiple comparisons post-hoc test (**P* < 0.05, ***P* < 0.01, ****P* < 0.001).

| A. B. 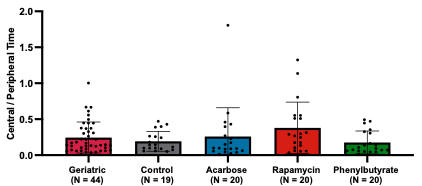 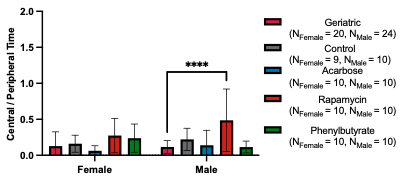 **Source** \| **SS** \| **df** \| **MS** \| **F** \| ***P***; Interaction \| 0.32 \| 4 \| 0.08 \| 2.09 \| 0.087; Sex \| 0.05 \| 1 \| 0.05 \| 1.33 \| 0.25; Group \| 1.08 \| 4 \| 0.27 \| 7.06 \| **<0.0001**; Residual \| 4.31 \| 113 \| 0.04 \| \|; Total \| 5.76 \| 122 \| \| \| | | | | | | | | |
| --- | --- | --- | --- | --- | --- | --- | --- | --- |
|  | Geriatric-Control | | Geriatric -Acarbose | | Geriatric-Rapamycin | | Geriatric-Phenylbutyrate | |
|  | *d*  (95% CI) | Adj. *P*-Value | *d*  (95% CI) | Adj. *P*-Value | *d*  (95% CI) | Adj. *P*-Value | *d*  (95% CI) | Adj. *P*-Value |
| Overall | -0.47  (-1.01, 0.07) | 0.91 | -0.55  (-1.08, -0.01) | >0.99 | -1.10  (-1.66, -0.54) | 0.21 | -0.39  (-0.92, 0.14) | 0.78 |
| Female | -0.16  (-0.94, 0.62) | 0.98 | 0.40  (-0.36, 1.16) | 0.84 | -0.64  (-1.41, 0.13) | 0.43 | -0.54  (-1.30, 0.23) | 0.19 |
| Male | -0.89  (-1.65, -0.12) | 0.47 | -0.14  (-0.88, 0.59) | >0.99 | -1.50  (-2.32, -0.68) | **<0.0001** | 0.11  (-0.63, 0.85) | >0.99 |

**Figure S7. Central-to-peripheral time ratio comparisons between historical geriatric crickets and treatment groups. (A)** Overall and **(B)** sex-stratified analyses present historical geriatric crickets relative to control, acarbose-, rapamycin-, and phenylbutyrate-treated cohorts. Results from the two-way ANOVA are provided above the table, summarizing the effects of treatment, sex, and their interaction on the central/peripheral time ratio. Pairwise comparisons were computed using Cohen’s *d* with Hedges’ *g* correction to reduce small sample size bias, with effect sizes reported alongside 95% CIs in the format (lower, upper). Historical geriatric crickets served as the reference group for all comparisons, and adjusted *P*-values were obtained using Dunnett’s multiple comparisons post-hoc test (*****P* < 0.0001).

| A. B. 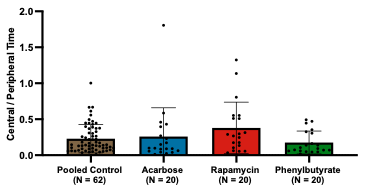 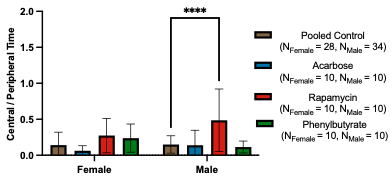 **Source** \| **SS** \| **df** \| **MS** \| **F** \| ***P***; Interaction \| 0.30 \| 3 \| 0.10 \| 2.60 \| 0.056; Sex \| 0.05 \| 1 \| 0.05 \| 1.21 \| 0.27; Group \| 1.00 \| 3 \| 0.33 \| 8.64 \| **<0.0001**; Residual \| 4.41 \| 114 \| 0.04 \| \|; Total \| 5.76 \| 121 \| \| \| | | | | | | |
| --- | --- | --- | --- | --- | --- | --- |
|  | Pooled Control-Acarbose | | Pooled Control-Rapamycin | | Pooled Control-Phenylbutyrate | |
|  | *d*  (95% CI) | Adj. *P*-Value | *d*  (95% CI) | Adj. *P*-Value | *d*  (95% CI) | Adj. *P*-Value |
| Overall | -0.12 (-0.62, 0.38) | 0.95 | -0.61 (-1.13, -0.10) | 0.08 | 0.28 (-0.23, 0.78) | 0.81 |
| Female | 0.49 (-0.24, 1.22) | 0.62 | -0.66  (-1.40, 0.07) | 0.18 | -0.53 (-1.26, 0.20) | 0.43 |
| Male | 0.07 (-0.64, 0.77) | >0.99 | -1.45  (-2.22, -0.68) | **<0.0001** | 0.35 (-0.36, 1.06) | 0.94 |

**Figure S8. Central-to-peripheral time ratio comparisons between pooled control crickets and treatment groups. (A)** Overall and **(B)** sex-stratified analyses show pooled controls compared with acarbose-, rapamycin-, and phenylbutyrate-treated cohorts. Results from the two-way ANOVA are presented above the table, outlining the effects of treatment group, sex, and their interaction on the central/peripheral time ratio. Pairwise comparisons were determined using Cohen’s *d* with Hedges’ *g* correction to account for small sample size bias, with effect sizes reported alongside 95% CIs in the format (lower, upper). Pooled controls served as the reference group for all comparisons, and adjusted *P*-values were calculated using Dunnett’s multiple comparisons post-hoc test (*****P* < 0.0001).

| **Central/Peripheral Speed** | | | | | | | | | |
| --- | --- | --- | --- | --- | --- | --- | --- | --- | --- |
|  | Overall | | | Female | | | Male | | |
|  | Mean (SD) | *d*  (95% CI) | *P* | Mean (SD) | *d*  (95% CI) | *P* | Mean (SD) | *d*  (95% CI) | *P* |
| Control  (N_Female_ = 9, N_Male_ = 10) | 3.13 (0.99) |  |  | 3.42 (1.03) |  |  | 2.87 (0.93) |  |  |
| Acarbose  (N_Female_ = 10, N_Male_ = 10) | 2.84 (1.26) | 0.25  (-0.38, 0.88) | 0.70 | 2.85 (1.55) | 0.41  (−0.50, 1.32) | 0.48 | 2.82 (0.98) | 0.05  (−0.83, 0.93) | >0.99 |
| Rapamycin  (N_Female_ = 10, N_Male_ = 10) | 2.28 (0.53) | 1.06  (0.39, 1.73) | **0.031** | 2.40 (0.60) | 1.17  (0.20, 2.15) | 0.086 | 2.17 (0.46) | 0.91  (-0.01, 1.83) | 0.29 |
| Phenylbutyrate  (N_Female_ = 10, N_Male_ = 10) | 3.30 (1.15) | -0.15  (-0.78, 0.47) | 0.92 | 3.66 (1.00) | -0.23  (-1.13, 0.68) | 0.91 | 2.93 (1.23) | -0.05  (-0.93, 0.82) | >0.99 |

**Table S4. Pairwise comparisons of central-to-peripheral speed ratio between control and treatment groups.** Overall and sex-stratified analyses are presented for the central/peripheral speed ratio in control, acarbose-, rapamycin-, and phenylbutyrate-treated crickets. Effect sizes were determined using Cohen’s *d* with Hedges’ *g* g correction to account for small sample size bias, with values reported alongside 95% CIs in the format (lower, upper). Controls were used as the reference group for all comparisons, and adjusted *P*-values were calculated using Dunnett’s multiple comparisons post-hoc test.

| A. B. 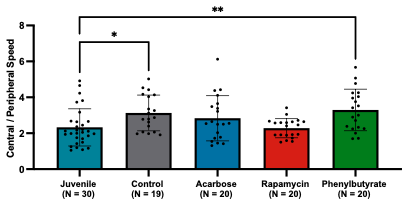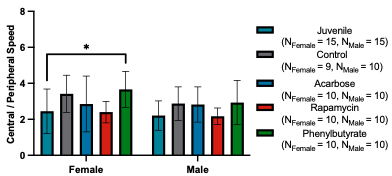 **Source** \| **SS** \| **df** \| **MS** \| **F** \| ***P***; Interaction \| 1.57 \| 4 \| 0.39 \| 0.37 \| 0.83; Sex \| 3.33 \| 1 \| 3.33 \| 3.14 \| 0.08; Group \| 18.77 \| 4 \| 4.69 \| 4.42 \| **0.0025**; Residual \| 105.1 \| 99 \| 1.06 \| \|; Total \| 128.8 \| 108 \| \| \| | | | | | | | | |
| --- | --- | --- | --- | --- | --- | --- | --- | --- |
|  | Juvenile-Control | | Juvenile-Acarbose | | Juvenile-Rapamycin | | Juvenile-Phenylbutyrate | |
|  | *d*  (95% CI) | Adj. *P*-Value | *d*  (95% CI) | Adj. *P*-Value | *d*  (95% CI) | Adj. *P*-Value | *d*  (95% CI) | Adj. *P*-Value |
| Overall | -0.77  (-1.37, -0.18) | **0.032** | -0.44  (-1.02, 0.13) | 0.27 | 0.06  (-0.51, 0.62) | >0.99 | -0.88  (-1.47, -0.29) | **0.0056** |
| Female | -0.81  (-1.66, 0.05) | 0.096 | -0.28  (-1.09, 0.52) | 0.76 | 0.05  (-0.75, 0.85) | >0.99 | -1.02  (-1.87, -0.17) | **0.018** |
| Male | -0.74  (-1.56, 0.09) | 0.35 | -0.67  (-1.49, 0.16) | 0.42 | 0.06  (-0.75, 0.86) | >0.99 | -0.70  (-1.52, 0.13) | 0.27 |

**Figure S9. Central-to-peripheral speed ratio comparisons between juvenile crickets and treatment groups. (A)** Overall and **(B)** sex-stratified analyses examine juvenile crickets in relation to control, acarbose-, rapamycin-, and phenylbutyrate-treated cohorts. The results from the two-way ANOVA are summarized above the table, describing the influence of treatment group, sex, and their interaction on the central/peripheral speed ratio. Pairwise contrasts were derived using Cohen’s *d* with Hedges’ *g* correction to limit bias from small sample sizes, with effect sizes shown alongside 95% CIs in the format (lower, upper). Juvenile crickets were designated as the reference group for all comparisons, and adjusted *P*-values were obtained through Dunnett’s multiple comparisons post-hoc test (**P* < 0.05, ***P* < 0.01).

| A. B. 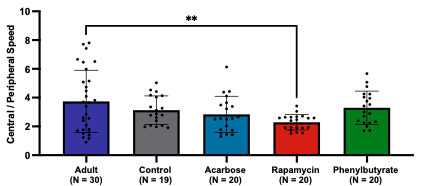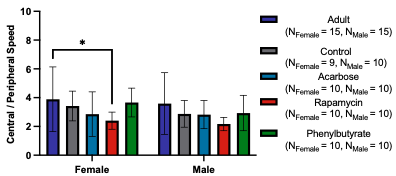 **Source** \| **SS** \| **df** \| **MS** \| **F** \| ***P***; Interaction \| 1.49 \| 4 \| 0.37 \| 0.18 \| 0.95; Sex \| 3.57 \| 1 \| 3.57 \| 1.69 \| 0.20; Group \| 27.78 \| 4 \| 6.94 \| 3.28 \| **0.014**; Residual \| 209.5 \| 99 \| 2.12 \| \|; Total \| 242.3 \| 108 \| \| \| | | | | | | | | |
| --- | --- | --- | --- | --- | --- | --- | --- | --- |
|  | Adult-Control | | Adult-Acarbose | | Adult-Rapamycin | | Adult-Phenylbutyrate | |
|  | *d*  (95% CI) | Adj. *P*-Value | *d*  (95% CI) | Adj. *P*-Value | *d*  (95% CI) | Adj. *P*-Value | *d*  (95% CI) | Adj. *P*-Value |
| Overall | 0.33  (-0.25, 0.91) | 0.43 | 0.48  (-0.10, 1.05) | 0.11 | 0.84  (0.25, 1.43) | **0.0025** | 0.24  (-0.33, 0.80) | 0.69 |
| Female | 0.24  (-0.59, 1.07) | 0.87 | 0.50  (-0.31, 1.32) | 0.26 | 0.81  (-0.02, 1.64) | **0.049** | 0.12  (-0.68, 0.92) | 0.99 |
| Male | 0.39  (-0.42, 1.20) | 0.59 | 0.42  (-0.39, 1.23) | 0.53 | 0.81  (-0.02, 1.64) | 0.065 | 0.35  (-0.46, 1.15) | 0.66 |

**Figure S10. Central-to-peripheral speed ratio comparisons between adult crickets and treatment groups. (A)** Overall and **(B)** sex-stratified analyses compare adult crickets against control, acarbose-, rapamycin-, and phenylbutyrate-treated cohorts. Results from the two-way ANOVA are presented above the table, outlining the effects of treatment, sex, and their interaction on the central/peripheral speed ratio. Pairwise comparisons were computed using Cohen’s *d* with Hedges’ *g* correction to address small sample size bias, with effect sizes reported together with 95% CIs in the format (lower, upper). Adult crickets served as the reference group for all comparisons, and adjusted *P*-values were calculated using Dunnett’s multiple comparisons post-hoc test (**P* < 0.05, ***P* < 0.01).

| A. B. 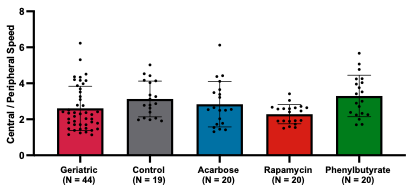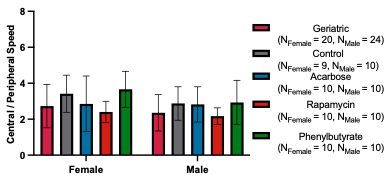 **Source** \| **SS** \| **df** \| **MS** \| **F** \| ***P***; Interaction \| 1.46 \| 4 \| 0.36 \| 0.33 \| 0.86; Sex \| 4.02 \| 1 \| 4.02 \| 3.62 \| 0.060; Group \| 15.30 \| 4 \| 3.83 \| 3.44 \| **0.011**; Residual \| 125.7 \| 113 \| 1.11 \| \|; Total \| 146.5 \| 122 \| \| \| | | | | | | | | |
| --- | --- | --- | --- | --- | --- | --- | --- | --- |
|  | Geriatric-Control | | Geriatric -Acarbose | | Geriatric-Rapamycin | | Geriatric-Phenylbutyrate | |
|  | *d*  (95% CI) | Adj. *P*-Value | *d*  (95% CI) | Adj. *P*-Value | *d*  (95% CI) | Adj. *P*-Value | *d*  (95% CI) | Adj. *P*-Value |
| Overall | -0.44  (-0.98, 0.10) | 0.28 | -0.18  (-0.71, 0.34) | 0.88 | 0.31  (-0.22, 0.83) | 0.69 | -0.56  (-1.10, -0.03) | 0.080 |
| Female | -0.58  (-1.37, 0.22) | 0.33 | -0.09  (-0.84, 0.67) | >0.99 | 0.30  (-0.45, 1.06) | 0.87 | -0.79  (-1.57, -0.01) | 0.087 |
| Male | -0.50  (-1.25, 0.24) | 0.55 | -0.45  (-1.19, 0.30) | 0.64 | 0.21  (-0.53, 0.95) | 0.98 | -0.52  (-1.26, 0.23) | 0.45 |

**Figure S11. Central-to-peripheral speed ratio comparisons between historical geriatric crickets and treatment groups. (A)** Overall and **(B)** sex-stratified analyses present historical geriatric crickets relative to control, acarbose-, rapamycin-, and phenylbutyrate-treated cohorts. Results from the two-way ANOVA are shown above the table, summarizing the effects of treatment, sex, and their interaction on the central/peripheral speed ratio. Pairwise differences were calculated using Cohen’s *d* with Hedges’ *g* correction to minimize small sample size bias, with effect sizes reported alongside 95% CIs in the format (lower, upper). Historical geriatric crickets served as the reference group for all comparisons, and adjusted *P*-values were determined using Dunnett’s multiple comparisons post-hoc test.

| A. B. 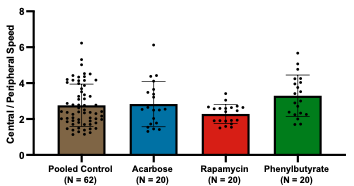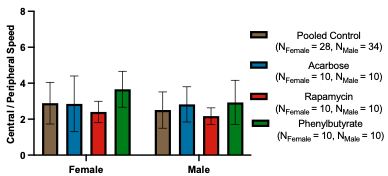 **Source** \| **SS** \| **df** \| **MS** \| **F** \| ***P***; Interaction \| 1.31 \| 3 \| 0.44 \| 0.39 \| 0.76; Sex \| 2.83 \| 1 \| 2.83 \| 2.52 \| 0.12; Group \| 10.64 \| 3 \| 3.55 \| 3.16 \| **0.028**; Residual \| 128.0 \| 114 \| 1.12 \| \|; Total \| 142.8 \| 121 \| \| \| | | | | | | |
| --- | --- | --- | --- | --- | --- | --- |
|  | Pooled Control-Acarbose | | Pooled Control-Rapamycin | | Pooled Control-Phenylbutyrate | |
|  | *d*  (95% CI) | Adj. *P*-Value | *d*  (95% CI) | Adj. *P*-Value | *d*  (95% CI) | Adj. *P*-Value |
| Overall | -0.07 (-0.57, 0.44) | 0.99 | 0.45 (-0.06, 0.95) | 0.25 | -0.46 (-0.96, 0.05) | 0.17 |
| Female | 0.03 (-0.69, 0.75) | >0.99 | 0.44 (-0.29, 1.17) | 0.49 | -0.67 (-1.41, 0.07) | 0.14 |
| Male | -0.30 (-1.01, 0.40) | 0.79 | 0.38 (-0.33, 1.09) | 0.74 | -0.39 (-1.10, 0.32) | 0.60 |

**Figure S12. Central-to-peripheral speed ratio comparisons between pooled control crickets and treatment groups. (A)** Overall and **(B)** sex-stratified analyses display pooled controls compared with acarbose-, rapamycin-, and phenylbutyrate-treated cohorts. The two-way ANOVA results are provided above the table, summarizing the contributions of treatment group, sex, and their interaction on the central/peripheral speed ratio. Pairwise comparisons were assessed using Cohen’s *d* with Hedges’ *g* correction to control for small sample size bias, with effect sizes listed alongside 95% CIs in the format (lower, upper). Pooled controls were used as the reference group for all comparisons, and adjusted *P*-values were derived using Dunnett’s multiple comparisons post-hoc test.

| **Overall Freezing Counts** | | | | | | | | | |
| --- | --- | --- | --- | --- | --- | --- | --- | --- | --- |
|  | Overall | | | Female | | | Male | | |
|  | Mean (SD) | *d*  (95% CI) | *P* | Mean (SD) | *d*  (95% CI) | *P* | Mean (SD) | *d*  (95% CI) | *P* |
| Control  (N_Female_ = 9, N_Male_ = 10) | 2302 (755) |  |  | 2020 (467.6) |  |  | 2555 (891.3) |  |  |
| Acarbose  (N_Female_ = 10, N_Male_ = 10) | 2985 (1239) | -0.65  (-1.29, 0.00) | 0.071 | 2699 (1199) | −0.70  (−1.63, 0.23) | 0.28 | 3271 (1274) | −0.62  (−1.52, 0.27) | 0.22 |
| Rapamycin  (N_Female_ = 10, N_Male_ = 10) | 3691 (984) | -1.55  (-2.26, -0.83) | **<0.0001** | 3670 (719.7) | -2.57  (-3.78, -1.35) | **0.001** | 3711 (1235) | -1.03  (-1.96, -0.10) | **0.021** |
| Phenylbutyrate  (N_Female_ = 10, N_Male_ = 10) | 2875 (721) | -0.76  (-1.41, -0.11) | 0.15 | 2615 (576.9) | -1.08  (-2.04, -0.11) | 0.38 | 3136 (782.5) | -0.66  (-1.56, 0.24) | 0.38 |

**Table S5. Pairwise comparisons of overall freezing counts between control and treatment groups.** Overall and sex-stratified analyses are provided for overall freezing counts in control, acarbose-, rapamycin-, and phenylbutyrate-treated crickets. Effect sizes for each comparison were calculated using Cohen’s *d* with Hedges’ *g* correction to minimize bias from small sample sizes, with values reported alongside 95% CIs in the format (lower, upper). Controls served as the reference group for all comparisons, and adjusted *P*-values were obtained using Dunnett’s multiple comparisons post-hoc test.

| A. B. 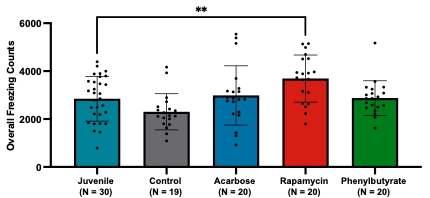 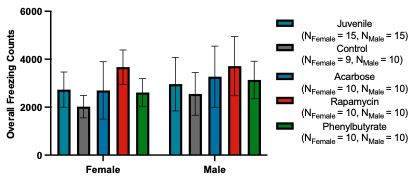 **Source** \| **SS** \| **df** \| **MS** \| **F** \| ***P***; Interaction \| 1.13e06 \| 4 \| 2.83e05 \| 0.32 \| 0.87; Sex \| 3.81e06 \| 1 \| 3.81e06 \| 4.27 \| 0.042; Group \| 1.97e07 \| 4 \| 4.93e06 \| 5.52 \| **0.0005**; Residual \| 8.83e07 \| 99 \| 8.92e05 \| \|; Total \| 1.13e08 \| 108 \| \| \| | | | | | | | | |
| --- | --- | --- | --- | --- | --- | --- | --- | --- |
|  | Juvenile-Control | | Juvenile-Acarbose | | Juvenile-Rapamycin | | Juvenile-Phenylbutyrate | |
|  | *d*  (95% CI) | Adj. *P*-Value | *d*  (95% CI) | Adj. *P*-Value | *d*  (95% CI) | Adj. *P*-Value | *d*  (95% CI) | Adj. *P*-Value |
| Overall | 0.61  (0.03, 1.20) | 0.17 | -0.13  (-0.69, 0.44) | 0.97 | -0.87  (-1.46, -0.28) | **0.0097** | -0.03  (-0.60, 0.53) | >0.99 |
| Female | 1.06  (0.18, 1.93) | 0.24 | 0.03  (-0.77, 0.83) | >0.99 | -1.25  (-2.12, -0.37) | 0.060 | 0.17  (-0.64, 0.97) | >0.99 |
| Male | 0.38  (-0.43, 1.19) | 0.70 | -0.25  (-1.06, 0.55) | 0.86 | -0.62  (-1.44, 0.19) | 0.18 | -0.17  (-0.97, 0.63) | 0.98 |

**Figure S13. Overall freezing count comparisons between juvenile crickets and treatment groups. (A)** Overall and **(B)** sex-stratified analyses evaluate juvenile crickets relative to control, acarbose-, rapamycin-, and phenylbutyrate-treated cohorts. Results from the two-way ANOVA are displayed above the table, outlining the effects of treatment group, sex, and their interaction on overall freezing counts. Pairwise comparisons were computed using Cohen’s *d* with Hedges’ *g* correction to account for small sample size bias, with effect sizes presented alongside 95% CIs in the format (lower, upper). Juvenile crickets served as the reference group for all comparisons, and adjusted *P*-values were obtained using Dunnett’s multiple comparisons post-hoc test (***P* < 0.01).

| A. B. 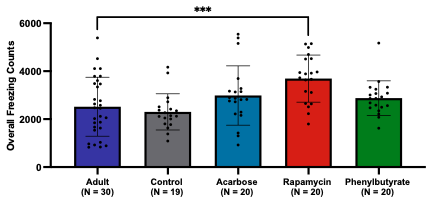 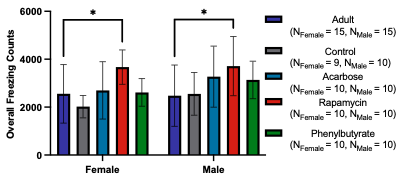 **Source** \| **SS** \| **df** \| **MS** \| **F** \| ***P***; Interaction \| 2.27e06 \| 4 \| 5.67e05 \| 0.52 \| 0.72; Sex \| 2.68e06 \| 1 \| 2.68e06 \| 2.47 \| 0.12; Group \| 2.38e07 \| 4 \| 5.95e06 \| 5.50 \| **0.0005**; Residual \| 1.07e08 \| 99 \| 1.08e06 \| \|; Total \| 1.36e08 \| 108 \| \| \| | | | | | | | | |
| --- | --- | --- | --- | --- | --- | --- | --- | --- |
|  | Adult-Control | | Adult-Acarbose | | Adult-Rapamycin | | Adult-Phenylbutyrate | |
|  | *d*  (95% CI) | Adj. *P*-Value | *d*  (95% CI) | Adj. *P*-Value | *d*  (95% CI) | Adj. *P*-Value | *d*  (95% CI) | Adj. *P*-Value |
| Overall | 0.20  (-0.38, 0.77) | 0.90 | -0.37  (-0.94, 0.20) | 0.36 | -1.01  (-1.61, -0.41) | **0.0006** | -0.33  (-0.90, 0.24) | 0.60 |
| Female | 0.51  (-0.33, 1.35) | 0.58 | -0.11  (-0.91, 0.69) | 0.99 | -1.02  (-1.87, -0.17) | **0.037** | -0.05  (-0.86, 0.75) | >0.99 |
| Male | -0.06  (-0.86, 0.74) | >0.99 | -0.60  (-1.42, 0.22) | 0.21 | -0.94  (-1.79, -0.10) | **0.017** | -0.57  (-1.39, 0.24) | 0.37 |

**Figure S14. Overall freezing count comparisons between adult crickets and treatment groups. (A)** Overall and **(B)** sex-stratified analyses compare adult crickets with control, acarbose-, rapamycin-, and phenylbutyrate-treated cohorts. The two-way ANOVA results are presented above the table, summarizing the contributions of treatment, sex, and their interaction on overall freezing counts. Pairwise comparisons were calculated using Cohen’s *d* with Hedges’ *g* correction to adjust for small sample size bias, with effect sizes reported together with 95% CIs in the format (lower, upper). Adult crickets served as the reference group for all comparisons, and adjusted *P*-values were determined using Dunnett’s multiple comparisons post-hoc test (**P* < 0.05, ****P* < 0.001).

| A. B. 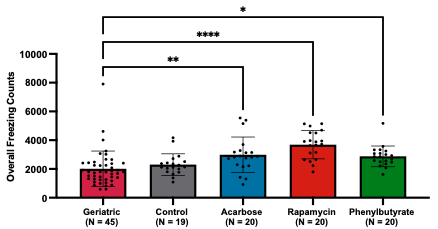 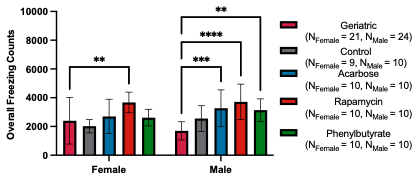 **Source** \| **SS** \| **df** \| **MS** \| **F** \| ***P***; Interaction \| 9.86e06 \| 4 \| 2.47e06 \| 2.26 \| 0.068; Sex \| 1.04e06 \| 1 \| 1.04e06 \| 0.95 \| 0.33; Group \| 4.43e07 \| 4 \| 1.11e07 \| 10.12 \| **<0.0001**; Residual \| 1.25e08 \| 114 \| 1.09e06 \| \|; Total \| 1.80e08 \| 123 \| \| \| | | | | | | | | |
| --- | --- | --- | --- | --- | --- | --- | --- | --- |
|  | Geriatric-Control | | Geriatric -Acarbose | | Geriatric-Rapamycin | | Geriatric-Phenylbutyrate | |
|  | *d*  (95% CI) | Adj. *P*-Value | *d*  (95% CI) | Adj. *P*-Value | *d*  (95% CI) | Adj. *P*-Value | *d*  (95% CI) | Adj. *P*-Value |
| Overall | -0.26  (-0.79, 0.28) | 0.76 | -0.78  (-1.32, -0.23) | **0.0035** | -1.43  (-2.01, -0.84) | **<0.0001** | -0.77  (-1.32, -0.23) | **0.012** |
| Female | 0.26  (-0.53, 1.04) | 0.83 | -0.20  (-0.96, 0.55) | 0.88 | -0.89  (-1.68, -0.11) | **0.0070** | -0.16  (-0.92, 0.59) | 0.96 |
| Male | -1.19  (-1.98, -0.40) | 0.10 | -1.79  (-2.65, -0.94) | **0.0004** | -2.34  (-3.26, -1.41) | **<0.0001** | -2.09  (-2.98, -1.20) | **0.0014** |

**Figure S15. Overall freezing count comparisons between historical geriatric crickets and treatment groups. (A)** Overall and **(B)** sex-stratified analyses present historical geriatric crickets relative to control, acarbose-, rapamycin-, and phenylbutyrate-treated cohorts. Results from the two-way ANOVA are provided above the table, detailing the effects of treatment group, sex, and their interaction on overall freezing counts. Pairwise comparisons were determined using Cohen’s *d* with Hedges’ *g* correction to minimize small sample size bias, with effect sizes displayed alongside 95% CIs in the format (lower, upper). Historical geriatric crickets served as the reference group for all comparisons, and adjusted *P*-values were calculated using Dunnett’s multiple comparisons post-hoc test (**P* < 0.05, ***P* < 0.01, ****P* < 0.001, *****P* < 0.0001).

| A. B. 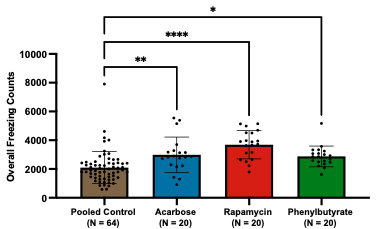 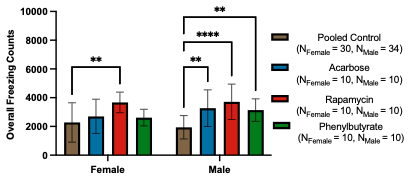 **Source** \| **SS** \| **df** \| **MS** \| **F** \| ***P***; Interaction \| 4.80e06 \| 3 \| 1.60e06 \| 1.42 \| 0.24; Sex \| 9.60e05 \| 1 \| 9.60e05 \| 0.85 \| 0.36; Group \| 4.36e07 \| 3 \| 1.45e07 \| 12.87 \| **<0.0001**; Residual \| 1.31e08 \| 116 \| 1.13e06 \| \|; Total \| 1.89e08 \| 123 \| \| \| | | | | | | |
| --- | --- | --- | --- | --- | --- | --- |
|  | Pooled Control-Acarbose | | Pooled Control-Rapamycin | | Pooled Control-Phenylbutyrate | |
|  | *d*  (95% CI) | Adj. *P*-Value | *d*  (95% CI) | Adj. *P*-Value | *d*  (95% CI) | Adj. *P*-Value |
| Overall | -0.77 (-1.28, -0.25) | **0.0044** | -1.45 (-2.00, -0.91) | **<0.0001** | -0.74 (-1.26, -0.23) | **0.015** |
| Female | -0.31 (-1.03, 0.41) | 0.61 | -1.09  (-1.85, -0.34) | **0.0014** | -0.27 (-0.99, 0.45) | 0.75 |
| Male | -1.40 (-2.17, -0.64) | **0.0021** | -1.24 (-1.99, -0.49) | **<0.0001** | -1.46 (-2.23, -0.69) | **0.0066** |

**Figure S16. Overall freezing count comparisons between pooled control crickets and treatment groups. (A)** Overall and **(B)** sex-stratified analyses show pooled controls compared with acarbose-, rapamycin-, and phenylbutyrate-treated cohorts. The results from the two-way ANOVA are displayed above the table, summarizing the effects of treatment, sex, and their interaction on overall freezing counts. Pairwise comparisons were computed using Cohen’s *d* with Hedges’ *g* correction to address small sample size bias, with effect sizes reported alongside 95% CIs in the format (lower, upper). Pooled controls were used as the reference group for all comparisons, and adjusted *P*-values were derived through Dunnett’s multiple comparisons post-hoc test (**P* < 0.05, ***P* < 0.01, *****P* < 0.0001).

| **Central Freezing Counts** | | | | | | | | | |
| --- | --- | --- | --- | --- | --- | --- | --- | --- | --- |
|  | Overall | | | Female | | | Male | | |
|  | Mean (SD) | *d*  (95% CI) | *P* | Mean (SD) | *d*  (95% CI) | *P* | Mean (SD) | *d*  (95% CI) | *P* |
| Control  (N_Female_ = 9, N_Male_ = 10) | 412.9 (304.0) |  |  | 346 (254.8) |  |  | 473 (344.4) |  |  |
| Acarbose  (N_Female_ = 10, N_Male_ = 10) | 401.2 (748.1) | 0.02  (-0.61, 0.65) | >0.99 | 154 (161.9) | 0.87  (−0.07, 1.81) | 0.88 | 649 (1009) | −0.22  (−1.10, 0.66) | 0.90 |
| Rapamycin  (N_Female_ = 10, N_Male_ = 10) | 1526 (1109) | -1.33  (-2.02, -0.63) | **<0.0001** | 1298 (926.0) | -1.31  (-2.30, -0.31) | **0.014** | 1753 (1274) | -1.31  (-2.28, -0.35) | **0.0005** |
| Phenylbutyrate  (N_Female_ = 10, N_Male_ = 10) | 547.3 (412.7) | -0.36  (-0.99, 0.27) | 0.89 | 591 (458.1) | -0.62  (-1.54, 0.30) | 0.79 | 503 (381.4) | -0.08  (-0.96, 0.80) | >0.99 |

**Table S6. Pairwise comparisons of central freezing counts between control and treatment groups.** Overall and sex-stratified analyses are shown for central freezing counts in control, acarbose-, rapamycin-, and phenylbutyrate-treated crickets. Effect sizes were computed using Cohen’s *d* with Hedges’ *g* correction to account for small sample size bias, with values reported together with 95% CIs in the format (lower, upper). Controls were used as the reference group for all comparisons, and adjusted *P*-values were calculated using Dunnett’s multiple comparisons post-hoc test.

| A. B. 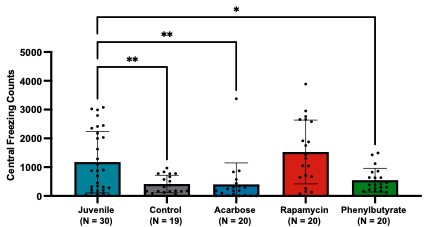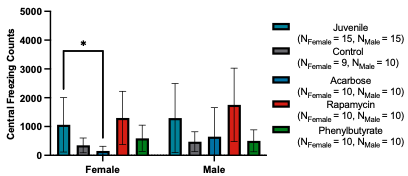 **Source** \| **SS** \| **df** \| **MS** \| **F** \| ***P***; Interaction \| 1.16e06 \| 4 \| 2.89e05 \| 0.41 \| 0.80; Sex \| 1.58e06 \| 1 \| 1.58e06 \| 2.26 \| 0.14; Group \| 2.19e07 \| 4 \| 5.47e06 \| 7.82 \| **<0.0001**; Residual \| 6.92e07 \| 99 \| 6.99e05 \| \|; Total \| 9.38e07 \| 108 \| \| \| | | | | | | | | |
| --- | --- | --- | --- | --- | --- | --- | --- | --- |
|  | Juvenile-Control | | Juvenile-Acarbose | | Juvenile-Rapamycin | | Juvenile-Phenylbutyrate | |
|  | *d*  (95% CI) | Adj. *P*-Value | *d*  (95% CI) | Adj. *P*-Value | *d*  (95% CI) | Adj. *P*-Value | *d*  (95% CI) | Adj. *P*-Value |
| Overall | 0.87  (0.27, 1.47) | **0.0086** | 0.80  (0.21, 1.39) | **0.0064** | -0.32  (-0.89, 0.25) | 0.43 | 0.71  (0.13, 1.29) | **0.037** |
| Female | 0.90  (0.03, 1.76) | 0.15 | 1.18  (0.31, 2.04) | **0.034** | -0.25  (-1.05, 0.56) | 0.91 | 0.57  (-0.24, 1.39) | 0.48 |
| Male | 0.82  (-0.01, 1.66) | 0.064 | 0.55  (-0.26, 1.37) | 0.20 | -0.36  (-1.17, 0.44) | 0.50 | 0.79  (-0.04, 1.62) | 0.08 |

**Figure S17. Central freezing count comparisons between juvenile crickets and treatment groups. (A)** Overall **and (B)** sex-stratified analyses evaluate juvenile crickets relative to control, acarbose-, rapamycin-, and phenylbutyrate-treated cohorts. Results from the two-way ANOVA are presented above the table, summarizing the effects of treatment group, sex, and their interaction on central freezing counts. Pairwise comparisons were calculated using Cohen’s *d* with Hedges’ *g* correction to control for small sample size bias, with effect sizes reported alongside 95% CIs in the format (lower, upper). Juvenile crickets served as the reference group for all comparisons, and adjusted *P*-values were obtained using Dunnett’s multiple comparisons post-hoc test (**P* < 0.05, ***P* < 0.01).

| A. B. 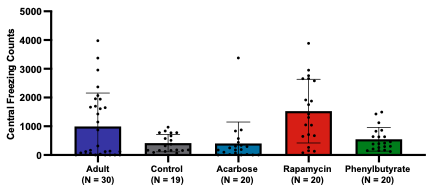 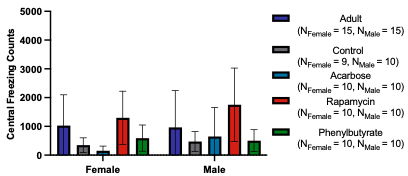 **Source** \| **SS** \| **df** \| **MS** \| **F** \| ***P***; Interaction \| 1.68e06 \| 4 \| 4.21e05 \| 0.55 \| 0.70; Sex \| 9.08e05 \| 1 \| 9.08e05 \| 1.19 \| 0.28; Group \| 1.90e07 \| 4 \| 4.75e06 \| 6.23 \| **0.0002**; Residual \| 7.55e07 \| 99 \| 7.63e05 \| \|; Total \| 1.05e08 \| 108 \| \| \| | | | | | | | | |
| --- | --- | --- | --- | --- | --- | --- | --- | --- |
|  | Adult-Control | | Adult-Acarbose | | Adult-Rapamycin | | Adult-Phenylbutyrate | |
|  | *d*  (95% CI) | Adj. *P*-Value | *d*  (95% CI) | Adj. *P*-Value | *d*  (95% CI) | Adj. *P*-Value | *d*  (95% CI) | Adj. *P*-Value |
| Overall | 0.62  (0.03, 1.20) | 0.083 | 0.57  (0.00, 1.15) | 0.068 | -0.46  (-1.03, 0.11) | 0.12 | 0.47  (-0.10, 1.04) | 0.24 |
| Female | 0.76  (-0.10, 1.61) | 0.22 | 1.00  (0.15, 1.85) | 0.057 | -0.26  (-1.06, 0.55) | 0.88 | 0.48  (-0.33, 1.29) | 0.59 |
| Male | 0.47  (-0.35, 1.28) | 0.48 | 0.26  (-0.55, 1.06) | 0.81 | -0.60  (-1.41, 0.22) | 0.10 | 0.43  (-0.37, 1.24) | 0.53 |

**Figure S18. Central freezing count comparisons between adult crickets and treatment groups. (A)** Overall and **(B)** sex-stratified analyses compare adult crickets to control, acarbose-, rapamycin-, and phenylbutyrate-treated cohorts. Results from the two-way ANOVA are shown above the table, detailing the effects of treatment group, sex, and their interaction on central freezing counts. Pairwise comparisons were computed using Cohen’s *d* with Hedges’ *g* correction to account for small sample size bias, with effect sizes displayed alongside 95% CIs in the format (lower, upper). Adult crickets served as the reference group for all comparisons, and adjusted *P*-values were determined using Dunnett’s multiple comparisons post-hoc test.

| A. B. 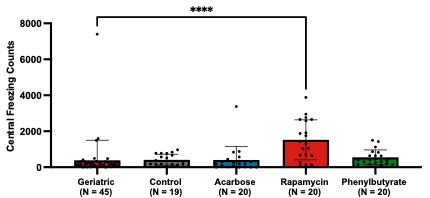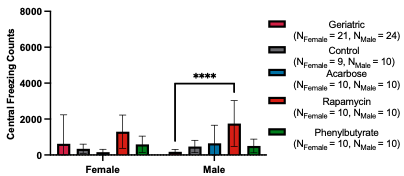 **Source** \| **SS** \| **df** \| **MS** \| **F** \| ***P***; Interaction \| 4.59e06 \| 4 \| 1.15e06 \| 1.47 \| 0.22; Sex \| 3.29e05 \| 1 \| 3.29e05 \| 0.42 \| 0.52; Group \| 2.04e07 \| 4 \| 5.11e06 \| 6.55 \| **<0.0001**; Residual \| 8.89e07 \| 114 \| 7.80e05 \| \|; Total \| 1.14e08 \| 123 \| \| \| | | | | | | | | |
| --- | --- | --- | --- | --- | --- | --- | --- | --- |
|  | Geriatric-Control | | Geriatric -Acarbose | | Geriatric-Rapamycin | | Geriatric-Phenylbutyrate | |
|  | *d*  (95% CI) | Adj. *P*-Value | *d*  (95% CI) | Adj. *P*-Value | *d*  (95% CI) | Adj. *P*-Value | *d*  (95% CI) | Adj. *P*-Value |
| Overall | -0.03  (-0.56, 0.51) | >0.99 | -0.01  (-0.54, 0.51) | >0.99 | -1.01  (-1.57, -0.46) | **<0.0001** | -0.17  (-0.69, 0.36) | 0.93 |
| Female | 0.20  (-0.59, 0.98) | 0.88 | 0.34  (-0.42, 1.10) | 0.49 | -0.46  (-1.22, 0.30) | 0.17 | 0.02  (-0.73, 0.78) | >0.99 |
| Male | -1.34  (-2.14, -0.53) | 0.83 | -0.84  (-1.60, -0.07) | 0.47 | -2.24  (-3.15, -1.33) | **<0.0001** | -1.37  (-2.17, -0.56) | 0.77 |

**Figure S19. Central freezing count comparisons between historical geriatric crickets and treatment groups. (A)** Overall and **(B)** sex-stratified analyses present historical geriatric crickets in comparison with control, acarbose-, rapamycin-, and phenylbutyrate-treated cohorts. Results from the two-way ANOVA are provided above the table, summarizing the contributions of treatment group, sex, and their interaction on central freezing counts. Pairwise comparisons were determined using Cohen’s *d* with Hedges’ *g* correction to minimize small sample size bias, with effect sizes reported together with 95% CIs in the format (lower, upper). Historical geriatric crickets served as the reference group for all comparisons, and adjusted *P*-values were obtained using Dunnett’s multiple comparisons post-hoc test (*****P* < 0.0001).

| A. B. 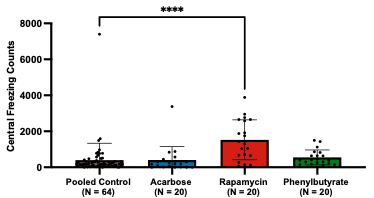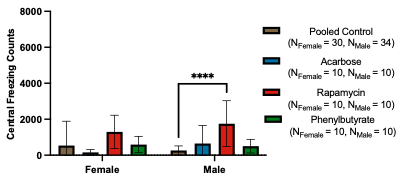 **Source** \| **SS** \| **df** \| **MS** \| **F** \| ***P***; Interaction \| 3.50e06 \| 3 \| 1.17e06 \| 1.50 \| 0.22; Sex \| 5.20e05 \| 1 \| 5.20e05 \| 0.67 \| 0.41; Group \| 2.05e07 \| 3 \| 6.82e06 \| 8.79 \| **<0.0001**; Residual \| 9.00e07 \| 116 \| 7.76e05 \| \|; Total \| 1.15e08 \| 123 \| \| \| | | | | | | |
| --- | --- | --- | --- | --- | --- | --- |
|  | Pooled Control-Acarbose | | Pooled Control-Rapamycin | | Pooled Control-Phenylbutyrate | |
|  | *d*  (95% CI) | Adj. *P*-Value | *d*  (95% CI) | Adj. *P*-Value | *d*  (95% CI) | Adj. *P*-Value |
| Overall | -0.01 (-0.51, 0.49) | >0.99 | -1.14 (-1.67, -0.61) | **<0.0001** | -0.18 (-0.68, 0.32) | 0.87 |
| Female | 0.32 (-0.40, 1.04) | 0.53 | -0.59  (-1.32, 0.14) | 0.057 | -0.04 (-0.76, 0.67) | >0.99 |
| Male | -0.73 (-1.45, -0.01) | 0.53 | -2.04 (-2.87, -1.22) | **<0.0001** | -0.82 (-1.55, -0.10) | 0.83 |

**Figure S20. Central freezing count comparisons between pooled control crickets and treatment groups. (A)** Overall and **(B)** sex-stratified analyses compare pooled controls with acarbose-, rapamycin-, and phenylbutyrate-treated cohorts. Results from the two-way ANOVA are displayed above the table, outlining the effects of treatment group, sex, and their interaction on central freezing counts. Pairwise comparisons were calculated using Cohen’s *d* with Hedges’ *g* correction to reduce small sample size bias, with effect sizes presented alongside 95% CIs in the format (lower, upper). Pooled controls served as the reference group for all comparisons, and adjusted *P*-values were derived using Dunnett’s multiple comparisons post-hoc test (*****P* < 0.0001).

| **Peripheral Freezing Counts** | | | | | | | | | |
| --- | --- | --- | --- | --- | --- | --- | --- | --- | --- |
|  | Overall | | | Female | | | Male | | |
|  | Mean (SD) | *d*  (95% CI) | *P* | Mean (SD) | *d*  (95% CI) | *P* | Mean (SD) | *d*  (95% CI) | *P* |
| Control  (N_Female_ = 9, N_Male_ = 10) | 1889 (722) |  |  | 1674 (452.3) |  |  | 2082 (879.2) |  |  |
| Acarbose  (N_Female_ = 10, N_Male_ = 10) | 2584 (1032) | -0.76  (-1.41, -0.11) | **0.029** | 2546 (1128) | −0.95  (−1.90, 0.00) | 0.063 | 2622 (985.7) | −0.55  (−1.45, 0.34) | 0.33 |
| Rapamycin  (N_Female_ = 10, N_Male_ = 10) | 2165 (776) | -0.36  (-0.99, 0.27) | 0.60 | 2372 (742.8) | -1.07  (-2.03, -0.11) | 0.17 | 1958 (789.2) | 0.14  (-0.74, 1.02) | 0.98 |
| Phenylbutyrate  (N_Female_ = 10, N_Male_ = 10) | 2328 (754) | -0.58  (-1.22, 0.06) | 0.24 | 2024 (618.1) | -0.61  (-1.53, 0.31) | 0.68 | 2633 (782.6) | -0.63  (-1.53, 0.26) | 0.32 |

**Table S7. Pairwise comparisons of peripheral freezing counts between control and treatment groups.** Overall and sex-stratified analyses are presented for peripheral freezing counts in control, acarbose-, rapamycin-, and phenylbutyrate-treated crickets. Effect sizes were calculated using Cohen’s *d* with Hedges’ *g* correction to minimize small sample size bias, with values reported alongside 95% CIs in the format (lower, upper). Controls served as the reference group for all comparisons, and adjusted *P*-values were obtained through Dunnett’s multiple comparisons post-hoc test.

| A. B. 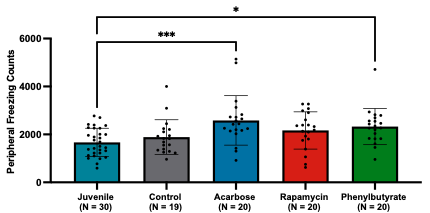 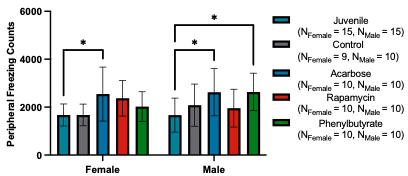 **Source** \| **SS** \| **df** \| **MS** \| **F** \| ***P***; Interaction \| 3.13e06 \| 4 \| 7.84e05 \| 1.33 \| 0.26; Sex \| 4.83e05 \| 1 \| 4.83e05 \| 0.82 \| 0.37; Group \| 1.23e07 \| 4 \| 3.07e06 \| 5.21 \| **0.0008**; Residual \| 5.84e07 \| 99 \| 5.90e05 \| \|; Total \| 7.43e07 \| 108 \| \| \| | | | | | | | | |
| --- | --- | --- | --- | --- | --- | --- | --- | --- |
|  | Juvenile-Control | | Juvenile-Acarbose | | Juvenile-Rapamycin | | Juvenile-Phenylbutyrate | |
|  | *d*  (95% CI) | Adj. *P*-Value | *d*  (95% CI) | Adj. *P*-Value | *d*  (95% CI) | Adj. *P*-Value | *d*  (95% CI) | Adj. *P*-Value |
| Overall | -0.34  (-0.92, 0.24) | 0.76 | -1.13  (-1.74, -0.53) | **0.0003** | -0.73  (-1.31, -0.15) | 0.10 | -0.98  (-1.58, -0.39) | **0.014** |
| Female | -0.01  (-0.83, 0.82) | >0.99 | -1.07  (-1.92, -0.22) | **0.023** | -1.16  (-2.02, -0.29) | 0.10 | -0.65  (-1.47, 0.17) | 0.65 |
| Male | -0.51  (-1.32, 0.30) | 0.51 | -1.11  (-1.97, -0.25) | **0.011** | -0.38  (-1.18, 0.43) | 0.78 | -1.26  (-2.13, -0.39) | **0.010** |

**Figure S21. Peripheral freezing count comparisons between juvenile crickets and treatment groups. (A)** Overall and **(B)** sex-stratified analyses evaluate juvenile crickets relative to control, acarbose-, rapamycin-, and phenylbutyrate-treated cohorts. Results from the two-way ANOVA are presented above the table, summarizing the effects of treatment group, sex, and their interaction on peripheral freezing counts. Pairwise comparisons were computed using Cohen’s *d* with Hedges’ *g* correction to account for small sample size bias, with effect sizes reported alongside 95% CIs in the format (lower, upper). Juvenile crickets served as the reference group for all comparisons, and adjusted *P*-values were calculated using Dunnett’s multiple comparisons post-hoc test (**P* < 0.05, ****P* < 0.001).

| A. B. 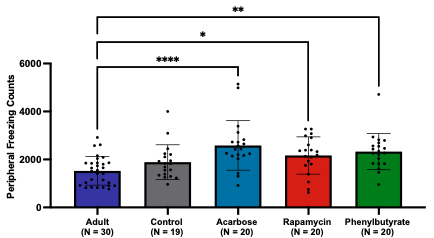 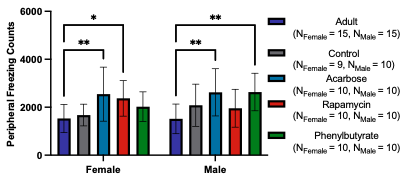 **Source** \| **SS** \| **df** \| **MS** \| **F** \| ***P***; Interaction \| 3.16e06 \| 4 \| 7.89e05 \| 1.34 \| 0.26; Sex \| 4.67e05 \| 1 \| 4.67e05 \| 0.79 \| 0.38; Group \| 1.64e07 \| 4 \| 4.10e06 \| 6.94 \| **<0.0001**; Residual \| 5.85e07 \| 99 \| 5.91e05 \| \|; Total \| 7.85e07 \| 108 \| \| \| | | | | | | | | |
| --- | --- | --- | --- | --- | --- | --- | --- | --- |
|  | Adult-Control | | Adult-Acarbose | | Adult-Rapamycin | | Adult-Phenylbutyrate | |
|  | *d*  (95% CI) | Adj. *P*-Value | *d*  (95% CI) | Adj. *P*-Value | *d*  (95% CI) | Adj. *P*-Value | *d*  (95% CI) | Adj. *P*-Value |
| Overall | -0.56  (-1.14, 0.03) | 0.33 | -1.31  (-1.93, -0.69) | **<0.0001** | -0.94  (-1.54, -0.35) | **0.018** | -1.20  (-1.81, -0.59) | **0.0018** |
| Female | -0.26  (-1.09, 0.57) | 0.98 | -1.17  (-2.03, -0.30) | **0.0063** | -1.25  (-2.12, -0.37) | **0.032** | -0.80  (-1.63, 0.03) | 0.35 |
| Male | -0.75  (-1.58, 0.08) | 0.23 | -1.37  (-2.25, -0.48) | **0.0025** | -0.62  (-1.44, 0.20) | 0.45 | -1.57  (-2.48, -0.66) | **0.0025** |

**Figure S22. Peripheral freezing count comparisons between adult crickets and treatment groups. (A)** Overall and **(B)** sex-stratified analyses compare adult crickets with control, acarbose-, rapamycin-, and phenylbutyrate-treated cohorts. Results from the two-way ANOVA are shown above the table, outlining the effects of treatment group, sex, and their interaction on peripheral freezing counts. Pairwise comparisons were determined using Cohen’s *d* with Hedges’ *g* correction to reduce small sample size bias, with effect sizes presented alongside 95% CIs in the format (lower, upper). Adult crickets served as the reference group for all comparisons, and adjusted *P*-values were obtained using Dunnett’s multiple comparisons post-hoc test (**P* < 0.05, ***P* < 0.01, *****P* < 0.0001).

| A. B. **Source** \| **SS** \| **df** \| **MS** \| **F** \| ***P***; Interaction \| 4.26e06 \| 4 \| 1.07e06 \| 1.60 \| 0.18; Sex \| 1.97e05 \| 1 \| 1.97e05 \| 0.30 \| 0.59; Group \| 1.57e07 \| 4 \| 3.93e06 \| 5.89 \| **0.0002**; Residual \| 7.60e07 \| 114 \| 6.67e05 \| \|; Total \| 9.61e07 \| 123 \| \| \| | | | | | | | | |
| --- | --- | --- | --- | --- | --- | --- | --- | --- |
|  | Geriatric-Control | | Geriatric -Acarbose | | Geriatric-Rapamycin | | Geriatric-Phenylbutyrate | |
|  | *d*  (95% CI) | Adj. *P*-Value | *d*  (95% CI) | Adj. *P*-Value | *d*  (95% CI) | Adj. *P*-Value | *d*  (95% CI) | Adj. *P*-Value |
| Overall | -0.33  (-0.87, 0.21) | 0.64 | -1.08  (-1.63, -0.52) | **0.0001** | -0.67  (-1.21, -0.13) | 0.059 | -0.88  (-1.43, -0.33) | **0.0073** |
| Female | 0.10  (-0.68, 0.88) | >0.99 | -0.73  (-1.50, 0.05) | 0.052 | -0.64  (-1.41, 0.13) | 0.19 | -0.28  (-1.04, 0.48) | 0.86 |
| Male | -0.84  (-1.60, -0.07) | 0.22 | -1.53  (-2.35, -0.71) | **0.0017** | -0.69  (-1.45, 0.06) | 0.43 | -1.73  (-2.57, -0.88) | **0.0015** |

**Figure S23. Peripheral freezing count comparisons between historical geriatric crickets and treatment groups. (A)** Overall and **(B)** sex-stratified analyses present historical geriatric crickets relative to control, acarbose-, rapamycin-, and phenylbutyrate-treated cohorts. Results from the two-way ANOVA are provided above the table, summarizing the effects of treatment, sex, and their interaction on peripheral freezing counts. Pairwise comparisons were calculated using Cohen’s *d* with Hedges’ *g* correction to correct for small sample size bias, with effect sizes reported alongside 95% CIs in the format (lower, upper). Historical geriatric crickets served as the reference group for all comparisons, and adjusted *P*-values were derived using Dunnett’s multiple comparisons post-hoc test (***P* < 0.01, ****P* < 0.001).

| A. B. **Source** \| **SS** \| **df** \| **MS** \| **F** \| ***P***; Interaction \| 2.79e06 \| 3 \| 9.32e05 \| 1.38 \| 0.25; Sex \| 6.68e04 \| 1 \| 6.68e04 \| 0.10 \| 0.75; Group \| 1.50e07 \| 3 \| 4.99e06 \| 7.38 \| **0.0001**; Residual \| 7.84e07 \| 116 \| 6.76e05 \| \|; Total \| 9.69e07 \| 123 \| \| \| | | | | | | |
| --- | --- | --- | --- | --- | --- | --- |
|  | Pooled Control-Acarbose | | Pooled Control-Rapamycin | | Pooled Control-Phenylbutyrate | |
|  | *d*  (95% CI) | Adj. *P*-Value | *d*  (95% CI) | Adj. *P*-Value | *d*  (95% CI) | Adj. *P*-Value |
| Overall | -1.03 (-1.55, -0.50) | **0.0002** | -0.46 (-0.97, 0.05) | 0.086 | -0.80 (-1.31, -0.28) | **0.011** |
| Female | -0.85 (-1.59, -0.11) | **0.024** | -0.49 (-1.21, 0.23) | 0.10 | -0.34 (-1.06, 0.38) | 0.70 |
| Male | -1.19 (-1.94, -0.45) | **0.0052** | -0.40 (-1.11, 0.31) | 0.70 | -1.29 (-2.05, -0.54) | **0.0047** |

**Figure S24. Peripheral freezing count comparisons between pooled control crickets and treatment groups. (A)** Overall and **(B)** sex-stratified analyses compare pooled controls with acarbose-, rapamycin-, and phenylbutyrate-treated cohorts. Results from the two-way ANOVA are shown above the table, describing the effects of treatment group, sex, and their interaction on peripheral freezing counts. Pairwise comparisons were computed using Cohen’s *d* with Hedges’ *g* correction to minimize small sample size bias, with effect sizes reported alongside 95% CIs in the format (lower, upper). Pooled controls served as the reference group for all comparisons, and adjusted *P*-values were calculated using Dunnett’s multiple comparisons post-hoc test (**P* < 0.05, ***P* < 0.01, ****P* < 0.001).

**Appendix 5. Treadmill Running Assay**

| **Maximum Running Time (s)** | | | | | | | | | |
| --- | --- | --- | --- | --- | --- | --- | --- | --- | --- |
|  | Overall | | | Female | | | Male | | |
|  | Mean (SD) | *d*  (95% CI) | *P* | Mean (SD) | *d*  (95% CI) | *P* | Mean (SD) | *d*  (95% CI) | *P* |
| Control  (N_Female_ = 10, N_Male_ = 10) | 43.08 (8.22) |  |  | 41.83 (7.43) |  |  | 44.33 (9.17) |  |  |
| Acarbose  (N_Female_ = 10, N_Male_ = 8) | 46.13  (6.41) | -0.40  (-1.05, 0.24) | 0.40 | 45.40  (6.92) | -0.48  (-1.37, 0.41) | 0.54 | 47.04  (6.04) | -0.32  (-1.26, 0.61) | 0.76 |
| Rapamycin  (N_Female_ = 10, N_Male_ = 10) | 59.98  (6.03) | -2.30  (-3.10, -1.50) | **<0.0001** | 60.40  (5.95) | -2.64  (-3.84, -1.44) | **<0.0001** | 59.57  (6.41) | -1.84  (-2.89, -0.80) | **<0.0001** |
| Phenylbutyrate  (N_Female_ = 10, N_Male_ = 10) | 58.93  (7.09) | -2.02  (-2.79, -1.26) | **<0.0001** | 56.92  (8.02) | -1.87  (-2.92, -0.82) | **<0.0001** | 60.95  (5.72) | -2.08  (-3.17, -0.99) | **<0.0001** |

**Table S1. Pairwise comparisons of maximum running time between control and treatment groups.** Overall and sex-stratified analyses are presented for maximum running time in control, acarbose-, rapamycin-, and phenylbutyrate-treated crickets. Effect sizes were calculated using Cohen’s *d* with Hedges’ *g* correction to minimize small sample size bias, with values reported alongside 95% confidence intervals (CIs) in the format (lower, upper). Controls served as the reference group for all comparisons, and adjusted *P*-values were obtained through Dunnett’s multiple comparisons post-hoc test.

| A. B. **Source** \| **SS** \| **df** \| **MS** \| **F** \| ***P***; Interaction \| 62.80 \| 4 \| 15.70 \| 0.32 \| 0.87; Sex \| 75.12 \| 1 \| 75.12 \| 1.51 \| 0.22; Group \| 5043 \| 4 \| 1261 \| 25.36 \| **<0.0001**; Residual \| 4376 \| 88 \| 49.72 \| \|; Total \| 9557 \| 97 \| \| \| | | | | | | | | |
| --- | --- | --- | --- | --- | --- | --- | --- | --- |
|  | Adult-Control | | Adult-Acarbose | | Adult-Rapamycin | | Adult-Phenylbutyrate | |
|  | *d*  (95% CI) | Adj. *P*-Value | *d*  (95% CI) | Adj. *P*-Value | *d*  (95% CI) | Adj. *P*-Value | *d*  (95% CI) | Adj. *P*-Value |
| Overall | 2.00  (1.24, 2.76) | **<0.0001** | 1.83  (1.07, 2.58) | **<0.0001** | -0.23  (-0.85, 0.40) | 0.91 | -0.06  (-0.68, 0.56) | >0.99 |
| Female | 2.12  (1.02, 3.21) | **<0.0001** | 1.70  (0.68, 2.73) | **0.0007** | -0.39  (-1.27, 0.50) | 0.82 | 0.11  (-0.77, 0.99) | >0.99 |
| Male | 1.75  (0.72, 2.79) | **<0.0001** | 1.77  (0.67, 2.86) | **0.0017** | -0.05  (-0.93, 0.83) | >0.99 | -0.26  (-1.14, 0.62) | 0.95 |

**Figure S1. Maximum running time comparisons between adult crickets and treatment groups. (A)** Overall and **(B)** sex-stratified analyses compare adult crickets with control, acarbose-, rapamycin-, and phenylbutyrate-treated cohorts. Results from the two-way analysis of variance (ANOVA) are shown above the table, outlining the effects of treatment group, sex, and their interaction on maximum running time. Pairwise comparisons were determined using Cohen’s *d* with Hedges’ *g* correction to reduce small sample size bias, with effect sizes presented alongside 95% CIs in the format (lower, upper). Adult crickets served as the reference group for all comparisons, and adjusted *P*-values were obtained using Dunnett’s multiple comparisons post-hoc test (***P* < 0.01, ****P* < 0.001, *****P* < 0.0001).

| A. B. **Source** \| **SS** \| **df** \| **MS** \| **F** \| ***P***; Interaction \| 69.47 \| 4 \| 17.37 \| 0.28 \| 0.89; Sex \| 59.74 \| 1 \| 59.74 \| 0.96 \| 0.33; Group \| 4937 \| 4 \| 1234 \| 19.87 \| **<0.0001**; Residual \| 5466 \| 88 \| 62.12 \| \|; Total \| 10532 \| 97 \| \| \| | | | | | | | | |
| --- | --- | --- | --- | --- | --- | --- | --- | --- |
|  | Mid-Age-Control | | Mid-Age-Acarbose | | Mid-Age-Rapamycin | | Mid-Age-Phenylbutyrate | |
|  | *d*  (95% CI) | Adj. *P*-Value | *d*  (95% CI) | Adj. *P*-Value | *d*  (95% CI) | Adj. *P*-Value | *d*  (95% CI) | Adj. *P*-Value |
| Overall | 1.58  (0.87, 2.28) | **<0.0001** | 1.34  (0.64, 2.05) | **<0.0001** | -0.24  (-0.86, 0.38) | 0.82 | -0.11  (-0.73, 0.51) | 0.98 |
| Female | 1.61  (0.60, 2.62) | **<0.0001** | 1.28  (0.31, 2.24) | **0.0028** | -0.29  (-1.17, 0.59) | 0.86 | 0.08  (-0.80, 0.95) | >0.99 |
| Male | 1.40  (0.42, 2.38) | **0.0007** | 1.27  (0.25, 2.29) | **0.014** | -0.16  (-1.04, 0.72) | 0.99 | -0.33  (-1.21, 0.55) | 0.85 |

**Figure S2. Maximum running time comparisons between mid-age crickets and treatment groups. (A)** Overall and **(B)** sex-stratified analyses compare mid-age crickets with control, acarbose-, rapamycin-, and phenylbutyrate-treated cohorts. Results from the two-way ANOVA are shown above the table, outlining the effects of treatment group, sex, and their interaction on maximum running time. Pairwise comparisons were determined using Cohen’s *d* with Hedges’ *g* correction to reduce small sample size bias, with effect sizes presented alongside 95% CIs in the format (lower, upper). Mid-age crickets served as the reference group for all comparisons, and adjusted *P*-values were obtained using Dunnett’s multiple comparisons post-hoc test (**P* < 0.05, ***P* < 0.01, ****P* < 0.001, *****P* < 0.0001).

| A. B. **Source** \| **SS** \| **df** \| **MS** \| **F** \| ***P***; Interaction \| 127.1 \| 4 \| 31.78 \| 0.49 \| 0.75; Sex \| 41.14 \| 1 \| 41.14 \| 0.63 \| 0.43; Group \| 5188 \| 4 \| 1297 \| 19.83 \| **<0.0001**; Residual \| 7325 \| 112 \| 65.40 \| \|; Total \| 12681 \| 121 \| \| \| | | | | | | | | |
| --- | --- | --- | --- | --- | --- | --- | --- | --- |
|  | Geriatric-Control | | Geriatric-Acarbose | | Geriatric-Rapamycin | | Geriatric-Phenylbutyrate | |
|  | *d*  (95% CI) | Adj. *P*-Value | *d*  (95% CI) | Adj. *P*-Value | *d*  (95% CI) | Adj. *P*-Value | *d*  (95% CI) | Adj. *P*-Value |
| Overall | 0.42  (-0.11, 0.95) | 0.25 | 0.09  (-0.46, 0.64) | 0.99 | -1.51  (-2.10, -0.92) | **<0.0001** | -1.35  (-1.92, -0.77) | **<0.0001** |
| Female | 0.62  (-0.15, 1.39) | 0.22 | 0.24  (-0.52, 0.99) | 0.91 | -1.44  (-2.28, -0.61) | **0.0003** | -0.99  (-1.78, -0.20) | **0.013** |
| Male | 0.21  (-0.53, 0.96) | 0.93 | -0.07  (-0.88, 0.73) | >0.99 | -1.48  (-2.31, -0.66) | **0.0001** | -1.67  (-2.51, -0.82) | **<0.0001** |

**Figure S3. Maximum running time comparisons between historical geriatric crickets and treatment groups. (A)** Overall and **(B)** sex-stratified analyses compare historical geriatric crickets with control, acarbose-, rapamycin-, and phenylbutyrate-treated cohorts. Results from the two-way ANOVA are shown above the table, outlining the effects of treatment group, sex, and their interaction on maximum running time. Pairwise comparisons were determined using Cohen’s *d* with Hedges’ *g* correction to reduce small sample size bias, with effect sizes presented alongside 95% CIs in the format (lower, upper). Geriatric crickets served as the reference group for all comparisons, and adjusted *P*-values were obtained using Dunnett’s multiple comparisons post-hoc test (**P* < 0.05, ****P* < 0.001, *****P* < 0.0001).

| A. B. **Source** \| **SS** \| **df** \| **MS** \| **F** \| ***P***; Interaction \| 78.18 \| 3 \| 26.06 \| 0.39 \| 0.76; Sex \| 34.54 \| 1 \| 34.54 \| 0.52 \| 0.47; Group \| 4990 \| 3 \| 1663 \| 25.03 \| **<0.0001**; Residual \| 7577 \| 114 \| 66.47 \| \|; Total \| 12680 \| 121 \| \| \| | | | | | | |
| --- | --- | --- | --- | --- | --- | --- |
|  | Pooled Control-Acarbose | | Pooled Control-Rapamycin | | Pooled Control-Phenylbutyrate | |
|  | *d*  (95% CI) | Adj. *P*-Value | *d*  (95% CI) | Adj. *P*-Value | *d*  (95% CI) | Adj. *P*-Value |
| Overall | -0.04  (-0.57, 0.48) | >0.99 | -1.65  (-2.21, -1.09) | **<0.0001** | -1.49  (-2.04, -0.94) | **<0.0001** |
| Female | 0.04  (-0.68, 0.75) | >0.99 | -1.67  (-2.47, -0.87) | **<0.0001** | -1.22  (-1.98, -0.46) | **0.0008** |
| Male | -0.14  (-.92, 0.63) | 0.97 | -1.55  (-2.33, -0.77) | **<0.0001** | -1.73  (-2.53, -0.93) | **<0.0001** |

**Figure S4. Maximum running time comparisons between pooled control crickets and treatment groups. (A)** Overall and **(B)** sex-stratified analyses compare pooled control crickets with acarbose-, rapamycin-, and phenylbutyrate-treated cohorts. Results from the two-way ANOVA are shown above the table, outlining the effects of treatment group, sex, and their interaction on maximum running time. Pairwise comparisons were determined using Cohen’s *d* with Hedges’ *g* correction to reduce small sample size bias, with effect sizes presented alongside 95% CIs in the format (lower, upper). Pooled control crickets served as the reference group for all comparisons, and adjusted *P*-values were obtained using Dunnett’s multiple comparisons post-hoc test (****P* < 0.001, *****P* < 0.0001).

| **Maximum Jumping Distance (cm)** | | | | | | | | | |
| --- | --- | --- | --- | --- | --- | --- | --- | --- | --- |
|  | Overall | | | Female | | | Male | | |
|  | Mean (SD) | *d*  (95% CI) | *P* | Mean (SD) | *d*  (95% CI) | *P* | Mean (SD) | *d*  (95% CI) | *P* |
| Control  (N_Female_ = 10, N_Male_ = 10) | 18.10  (8.02) |  |  | 18.80  (8.40) |  |  | 17.40  (8.00) |  |  |
| Acarbose  (N_Female_ = 10, N_Male_ = 8) | 21.39  (6.62) | -0.43  (-1.07, 0.21) | 0.40 | 19.60  (5.15) | -0.11  (-0.99, 0.77) | 0.99 | 23.63  (7.87) | -0.75  (-1.71, 0.21) | 0.21 |
| Rapamycin  (N_Female_ = 10, N_Male_ = 10) | 21.30  (6.41) | -0.43  (-1.05, 0.20) | 0.40 | 20.50  (3.92) | -0.25  (-1.13, 0.63) | 0.92 | 22.10  (8.36) | -0.55  (-1.44, 0.34) | 0.37 |
| Phenylbutyrate  (N_Female_ = 10, N_Male_ = 10) | 17.00  (8.72) | 0.13  (-0.49, 0.75) | 0.94 | 14.40  (10.06) | 0.45  (-0.43, 1.34) | 0.42 | 19.60  (6.65) | -0.29  (-1.17, 0.59) | 0.85 |

**Table S2. Pairwise comparisons of maximum jumping distance between control and treatment groups.** Overall and sex-stratified analyses are presented for maximum jumping distance in control, acarbose-, rapamycin-, and phenylbutyrate-treated crickets. Effect sizes were calculated using Cohen’s *d* with Hedges’ *g* correction to minimize small sample size bias, with values reported alongside 95% CIs in the format (lower, upper). Controls served as the reference group for all comparisons, and adjusted *P*-values were obtained through Dunnett’s multiple comparisons post-hoc test.

| A. B. **Source** \| **SS** \| **df** \| **MS** \| **F** \| ***P***; Interaction \| 238.3 \| 4 \| 59.56 \| 1.24 \| 0.30; Sex \| 40.27 \| 1 \| 40.27 \| 0.84 \| 0.36; Group \| 314.7 \| 4 \| 78.66 \| 1.64 \| 0.17; Residual \| 4225 \| 88 \| 48.01 \| \|; Total \| 4818 \| 97 \| \| \| | | | | | | | | |
| --- | --- | --- | --- | --- | --- | --- | --- | --- |
|  | Adult-Control | | Adult-Acarbose | | Adult-Rapamycin | | Adult-Phenylbutyrate | |
|  | *d*  (95% CI) | Adj. *P*-Value | *d*  (95% CI) | Adj. *P*-Value | *d*  (95% CI) | Adj. *P*-Value | *d*  (95% CI) | Adj. *P*-Value |
| Overall | 0.11  (-0.51, 0.73) | 0.99 | -0.47  (-1.11, 0.18) | 0.61 | -0.46  (-1.09, 0.17) | 0.62 | 0.26  (-0.36, 0.88) | 0.83 |
| Female | 0.21  -0.67, 1.09) | 0.97 | 0.14  (-0.74, 1.02) | >0.99 | -0.05  (-0.92, 0.83) | >0.99 | 0.73  (-0.18, 1.63) | 0.18 |
| Male | -0.02  (-0.89, 0.86) | >0.99 | -1.06  (-2.05, -0.07) | 0.18 | -0.73  (-1.64, 0.18) | 0.35 | -0.43  (-1.31, 0.46) | 0.87 |

**Figure S5. Maximum jumping distance comparisons between adult crickets and treatment groups. (A)** Overall and **(B)** sex-stratified analyses compare adult crickets with control, acarbose-, rapamycin-, and phenylbutyrate-treated cohorts. Results from the two-way ANOVA are shown above the table, outlining the effects of treatment group, sex, and their interaction on maximum jumping distance. Pairwise comparisons were determined using Cohen’s *d* with Hedges’ *g* correction to reduce small sample size bias, with effect sizes presented alongside 95% CIs in the format (lower, upper). Adult crickets served as the reference group for all comparisons, and adjusted *P*-values were obtained using Dunnett’s multiple comparisons post-hoc test.

| A. B. **Source** \| **SS** \| **df** \| **MS** \| **F** \| ***P***; Interaction \| 151.2 \| 4 \| 37.80 \| 0.63 \| 0.64; Sex \| 83.03 \| 1 \| 83.03 \| 1.38 \| 0.24; Group \| 388.1 \| 4 \| 97.03 \| 1.61 \| 0.18; Residual \| 5293 \| 88 \| 60.15 \| \|; Total \| 5915 \| 97 \| \| \| | | | | | | | | |
| --- | --- | --- | --- | --- | --- | --- | --- | --- |
|  | Mid-Age-Control | | Mid-Age-Acarbose | | Mid-Age-Rapamycin | | Mid-Age-Phenylbutyrate | |
|  | *d*  (95% CI) | Adj. *P*-Value | *d*  (95% CI) | Adj. *P*-Value | *d*  (95% CI) | Adj. *P*-Value | *d*  (95% CI) | Adj. *P*-Value |
| Overall | -0.11  (-0.73, 0.51) | 0.99 | -0.54  (-1.19, 0.11) | 0.28 | -0.54  (-1.17, 0.09) | 0.28 | 0.02  (-0.60, 0.64) | >0.99 |
| Female | -0.19  (-1.07, 0.68) | 0.98 | -0.39  (-1.27, 0.50) | 0.91 | -0.59  (-1.49, 0.30) | 0.76 | 0.33  (-0.55, 1.22) | 0.82 |
| Male | -0.03  (-0.91, 0.85) | >0.99 | -0.66  (-1.61, 0.29) | 0.24 | -0.51  (-1.40, 0.39) | 0.41 | -0.27  (-1.15, 0.61) | 0.88 |

**Figure S6. Maximum jumping distance comparisons between mid-age crickets and treatment groups. (A)** Overall and **(B)** sex-stratified analyses compare mid-age crickets with control, acarbose-, rapamycin-, and phenylbutyrate-treated cohorts. Results from the two-way ANOVA are shown above the table, outlining the effects of treatment group, sex, and their interaction on maximum jumping distance. Pairwise comparisons were determined using Cohen’s *d* with Hedges’ *g* correction to reduce small sample size bias, with effect sizes presented alongside 95% CIs in the format (lower, upper). Mid-age crickets served as the reference group for all comparisons, and adjusted *P*-values were obtained using Dunnett’s multiple comparisons post-hoc test.

| A. B. **Source** \| **SS** \| **df** \| **MS** \| **F** \| ***P***; Interaction \| 128.1 \| 4 \| 32.02 \| 0.64 \| 0.64; Sex \| 164.0 \| 1 \| 164.0 \| 3.26 \| 0.07; Group \| 1188 \| 4 \| 296.9 \| 5.89 \| **0.0002**; Residual \| 5642 \| 112 \| 50.38 \| \|; Total \| 7122 \| 121 \| \| \| | | | | | | | | |
| --- | --- | --- | --- | --- | --- | --- | --- | --- |
|  | Geriatric-Control | | Geriatric-Acarbose | | Geriatric-Rapamycin | | Geriatric-Phenylbutyrate | |
|  | *d*  (95% CI) | Adj. *P*-Value | *d*  (95% CI) | Adj. *P*-Value | *d*  (95% CI) | Adj. *P*-Value | *d*  (95% CI) | Adj. *P*-Value |
| Overall | -0.59  (-1.13, -0.05) | 0.11 | -1.14  (-1.72, -0.56) | **0.0011** | -1.14  (-1.70, -0.57) | **0.0008** | -0.42  (-0.96, 0.11) | 0.35 |
| Female | -0.89  (-1.67, -0.10) | 0.077 | -1.17  (-1.98, -0.37) | **0.037** | -1.39  (-2.22, -0.56) | **0.014** | -0.25  (-1.01, 0.51) | 0.91 |
| Male | -0.30  (-1.05, 0.44) | 0.88 | -1.21  (-2.06, -0.35) | **0.019** | -0.95  (-1.73, -0.18) | **0.046** | -0.66  (-1.42, 0.10) | 0.35 |

**Figure S7. Maximum jumping distance comparisons between historical geriatric crickets and treatment groups. (A)** Overall and **(B)** sex-stratified analyses compare historical geriatric crickets with control, acarbose-, rapamycin-, and phenylbutyrate-treated cohorts. Results from the two-way ANOVA are shown above the table, outlining the effects of treatment group, sex, and their interaction on maximum jumping distance. Pairwise comparisons were determined using Cohen’s *d* with Hedges’ *g* correction to reduce small sample size bias, with effect sizes presented alongside 95% CIs in the format (lower, upper). Geriatric crickets served as the reference group for all comparisons, and adjusted *P*-values were obtained using Dunnett’s multiple comparisons post-hoc test (**P* < 0.05, ***P* < 0.01, ****P* < 0.001).

| A. B. **Source** \| **SS** \| **df** \| **MS** \| **F** \| ***P***; Interaction \| 69.24 \| 3 \| 23.08 \| 0.44 \| 0.72; Sex \| 218.3 \| 1 \| 218.3 \| 4.18 \| **0.04**; Group \| 933.2 \| 3 \| 311.1 \| 5.96 \| **0.0008**; Residual \| 5949 \| 114 \| 52.18 \| \|; Total \| 7170 \| 121 \| \| \| | | | | | | |
| --- | --- | --- | --- | --- | --- | --- |
|  | Pooled Control-Acarbose | | Pooled Control-Rapamycin | | Pooled Control-Phenylbutyrate | |
|  | *d*  (95% CI) | Adj. *P*-Value | *d*  (95% CI) | Adj. *P*-Value | *d*  (95% CI) | Adj. *P*-Value |
| Overall | -0.87  (-1.41, -0.33) | 0.0054 | -0.86  (-1.38, -0.34) | **0.0041** | -0.23  (-0.74, 0.27) | 0.70 |
| Female | -0.71  (-1.44, 0.02) | 0.15 | -0.86  (-1.60, -0.12) | 0.068 | 0.01  (-0.70, 0.72) | >0.99 |
| Male | -1.07  (-1.88, -0.27) | **0.023** | -0.84  (-1.57, -0.11) | 0.055 | -0.53  (-1.25, 0.18) | 0.39 |

**Figure S8. Maximum running time comparisons between pooled control crickets and treatment groups. (A)** Overall and **(B)** sex-stratified analyses compare pooled control crickets with acarbose-, rapamycin-, and phenylbutyrate-treated cohorts. Results from the two-way ANOVA are shown above the table, outlining the effects of treatment group, sex, and their interaction on maximum jumping distance. Pairwise comparisons were determined using Cohen’s *d* with Hedges’ *g* correction to reduce small sample size bias, with effect sizes presented alongside 95% CIs in the format (lower, upper). Pooled control crickets served as the reference group for all comparisons, and adjusted *P*-values were obtained using Dunnett’s multiple comparisons post-hoc test (**P* < 0.05, ***P* < 0.01).

**Appendix 6. Percent Weight Change (%)**

| **Percent Weight Change (%)** | | | | | | | | | |
| --- | --- | --- | --- | --- | --- | --- | --- | --- | --- |
|  | Overall | | | Female | | | Male | | |
|  | Mean (SD) | *d*  (95% CI) | *P* | Mean (SD) | *d*  (95% CI) | *P* | Mean (SD) | *d*  (95% CI) | *P* |
| Control  (N_Female_ = 10, N_Male_ = 10) | 7.15 (23.97) |  |  | 9.25 (24.33) |  |  | 5.83 (24.18) |  |  |
| Acarbose  (N_Female_ = 10, N_Male_ = 8) | 18.96 (28.37) | 0.44  (0.00, 0.89) | 0.10 | 29.21 (27.26) | 0.75  (0.07, 1.42) | **0.033** | 5.85 (24.67) | 0.00  (-0.61, 0.61) | >0.99 |
| Rapamycin  (N_Female_ = 10, N_Male_ = 10) | 12.94 (24.23) | 0.24  (-0.20, 0.67) | 0.60 | 25.78 (24.46) | 0.66  (-0.03, 1.36) | 0.11 | 2.78 (18.96) | -0.14  (-0.70, 0.43) | 0.95 |
| Phenylbutyrate  (N_Female_ = 10, N_Male_ = 10) | 13.53 (24.88) | 0.26  (-0.17, 0.69) | 0.52 | 24.03 (28.75) | 0.54  (-0.15, 1.22) | 0.17 | 5.13 (17.74) | -0.03  (-0.59, 0.53) | >0.99 |

**Table S1. Pairwise comparisons of percent weight change between control and treatment groups.** Overall and sex-stratified analyses are presented for percent weight change in control, acarbose-, rapamycin-, and phenylbutyrate-treated crickets. Effect sizes were calculated using Cohen’s *d* with Hedges’ *g* correction to minimize small sample size bias, with values reported alongside 95% confidence intervals (CIs) in the format (lower, upper). Controls served as the reference group for all comparisons, and adjusted *P*-values were obtained through Dunnett’s multiple comparisons post-hoc test.

|  | Mean (SD) | | *d* (95% CI) | *P* |
| --- | --- | --- | --- | --- |
|  | Female | Male |  |  |
| Control | 9.25 (24.33) | 5.83 (24.18) | 0.14 (-0.78, 0.51) | >0.99 |
| Acarbose | 29.21 (27.26) | 5.85 (24.67) | -0.88 (-1.52, -0.23) | **0.0087** |
| Rapamycin | 25.78 (24.46) | 2.78 (18.96) | -1.05 (-1.69, -0.41) | **0.0079** |
| Phenylbutyrate | 24.03 (28.75) | 5.13 (17.74) | -0.80 (-1.41, -0.19) | **0.036** |

**Table S2. Group means and effect sizes for percent weight change comparing sexes within each treatment group.** Mean values with standard deviations (SD) are reported for percent weight change in control, acarbose-, rapamycin-, and phenylbutyrate-treated crickets. Pairwise comparisons are quantified using Cohen’s *d* with Hedges’ *g* correction to account for small sample size bias. Effect sizes are reported alongside 95% CIs in the format (lower, upper). Females served as the reference group for all comparisons. Adjusted *P*-values were calculated using Bonferroni’s multiple comparisons post-hoc test.

**Appendix 7. Lifespan**

A.	B. C. D.

|  | Control | | Phenylbutyrate | | Acarbose | | Rapamycin | |
| --- | --- | --- | --- | --- | --- | --- | --- | --- |
|  | Male | Female^1^ | Male | Female^1^ | Male | Female^1^ | Male | Female^1^ |
| Survival (days) | 97.0 | 112.0 | 83.0 | 116.5 | 87.5 | 93.5 | 132.5 | 136.0 |
| *HR* (95% CI) | 1.51 (0.75, 3.04) | | 3.22 (1.08, 9.65) | | 1.05 (0.37, 3.02) | | 0.93 (0.45, 1.94) | |
| Adj. *P* -value | 0.92 | | **0.0068** | | >0.99 | | >0.99 | |

^1^Reference groups for effect size estimations and statistical testing.

**Figure S1. Post-treatment lifespan following intermittent drug exposure. (A)** Sex comparison within control group showing no difference in lifespan. **(B)** Phenylbutyrate-treated females outlived their male counterparts. **(C–D)** Sex comparisons within the acarbose and rapamycin groups revealed no sex-specific differences in post-treatment lifespan.
